# Comparison of marker selection methods for high throughput scRNA-seq data

**DOI:** 10.1101/679761

**Authors:** Anna C. Gilbert, Alexander Vargo

## Abstract

Here, we evaluate the performance of a variety of marker selection methods on scRNA-seq UMI counts data. We test on an assortment of experimental and synthetic data sets that range in size from several thousand to one million cells. In addition, we propose several performance measures for evaluating the quality of a set of markers when there is no known ground truth. According to these metrics, most existing marker selection methods show similar performance on experimental scRNA-seq data; thus, the speed of the algorithm is the most important consideration for large data sets. With this in mind, we introduce RANKCORR, a fast marker selection method with strong mathematical underpinnings that takes a step towards sensible multi-class marker selection.

## Background

In single cell RNA sequencing (scRNA-seq), mRNA data are collected from individual cells, allowing for detailed descriptions of specific cell types and states. In recent years, the development of high throughput microfludic sequencing protocols has allowed for the collection of genetic information from up to one million individual cells in a single experiment. Additionally, the incorporation of unique molecular identifier (UMI) technology makes it possible to process these raw sequencing data into integer valued read counts.

Due to factors that are both biological (e.g. transcriptional bursting) and technical (e.g. 3’ bias in UMI based sequencing protocols) in nature, scRNA-seq counts data exhibit high variance and are very sparse (often, approximately 90% of the reads are 0). These characteristics, in combination with the integer valued quality of the counts and the high dimensionality of the data (often, 20,000 genes show nonzero expression levels in an experiment), make it so that scRNA-seq data does not match many of the models that underly common data analysis techniques. For this reason, many specialized tools have been developed to attempt to answer biological questions with scRNA-seq data.

One such biological question that has generated a significant amount of study (and produced a multitude of tools) in the scRNA-seq literature is the problem of finding *marker genes*. Marker genes are genes that can be used to distinguish the cells in one group apart from all other cells or from other specific cell types. In the (sc)RNA-seq literature, marker selection is almost always accomplished via differential expression (see [1], [2], [3], [4], [5] for some examples of methods that we examine in greater detail in the Methods; see [6] for a survey and evaluation of more methods). In order to find the genes that are useful for separating two populations of cells, a statistical test is applied to each gene in the data set to determine if the distributions of gene expression are different between the two populations: the genes with the most significance are selected as marker genes.

Marker selection has also received extensive study in the computer science literature, where is is known as the problem of “feature selection.” There are generally two main classes of feature selection algorithms: greedy algorithms that select features one-by-one, computing a score at each step to determine the next marker to select (for example, forward-or backward-stepwise selection, see Section 3.3 of [7]; mutual information based methods, see e.g. [8]; and other greedy methods e.g. [9]), and slower algorithms that are based on solving some regularized convex optimization problem (for example LASSO [10], Elastic Nets [11], and other related methods [12]).

In this work, we evaluate the performance of a diverse set of feature selection methods when they are applied to a representative sample of different types of UMI counts data sets, both experimentally generated and synthetic. We consider feature selection algorithms from the computer science literature along with complex statistical differential expression methods from the scRNA-seq literature: see the marker selection methods for a detailed list. Unlike the review [6], which also focuses on comparing differential expression methods for scRNA-seq data, we focus on large datasets generated by UMI based protocols. We consider data sets of multiple different sizes, data sets that contain cell differentiation trajectories, and data sets with well-differentiated cell types. Moreover, we consider cell type classifications that are both biologically motivated as well as cell clusters that are algorithmically created.

We also propose several metrics that can be used to evaluate the quality of marker set when a ground truth marker set is unknown. Using these evaluation metrics, there are generally only small differences between the different marker selection algorithms that are examined here, though all of the algorithms perform significantly better than random marker selection. Similar results are observed on synthetic data, where a set of ground truth markers is known. This suggests that fast and simple marker selection methods should be preferred over high complexity algorithms.

A major setback of many feature selection and differential expression algorithms is that they are not designed to handle data that contain more than two cell types. Using a differential expression method, for example, one strategy is to pick a fixed number (e.g. 10) of the statistically most significant genes for each cell type; there may be overlap in the genes selected for different cell types. This strategy does not take into account the fact that some cell types are more difficult to characterize than others, however: one cell type may require more than 10 markers to separate from the other cells, while a different cell type may be separated with only one marker. Setting a significance threshold for the statistical test does not solve this problem: a cell type that is easy to separate from other cells will often exhibit several high significance markers, while a cell type that is difficult to separate might not exhibit any high significance markers.

In this work we introduce a feature selection algorithm that takes steps to mitigate this issue. This method, which we call RANKCORR, is inspired by the method in [13] and relies on the ranking the scRNA-seq counts before marker selection. Below, we provide some intuition as to why ranking scRNA-seq data is a useful strategy for understanding scRNA-seq counts data. The method in [13] is based on convex optimization (which would generally be quite slow on large data sets); it is possible to find a quick deterministic solution to this optimization, however, so that RANKCORR runs quickly. RANKCORR is able to handle over one million cells in an amount of time that is competitive with simple statistical methods.

In the remainder of this background section, we introduce mathematical notation that will appear throughout this manuscript, discuss some of the instinct behind why it might be reasonable to rank scRNA-seq data, and finish with some mathematical background on feature selection and some existing feature selection algorithms. The benchmarking of feature selection methods appears in the Results and Discussion.

### Notation and definitions

Let ℝ denote the set of real numbers, ℤ denote the set of integers, and ℕ denote the set of natural numbers.

Consider an scRNA-seq experiment that collects gene expression information from *n* cells. After processing, for each cell that is sequenced, a vector *x* ∈ ℝ*^p^* is obtained: *x*_*j*_ represents the number of copies of a specific mRNA that was observed during the sequencing procedure (and we are assuming that *p* different mRNAs were detected during the experiment). When all *n* cells are sequenced, this results in *n* vectors in ℝ*^p^*, which we arrange into a data matrix *X* ∈ ℝ^*n*×*p*^. The entry *X*_*i,j*_ represents the number of counts of gene *j* in cell *i*. Note that this is the *transpose* of the data matrix that is common in the scRNA-seq literature.

Let [*n*] = {1, *…, n*}. For a matrix *X*, let *X*_*i*_ denote column *i* of *X*. Given a vector *x*, let 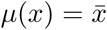 denote the average of the elements of *x* and let *σ*(*x*) represent the standard deviation of the elements in *x*; that is, 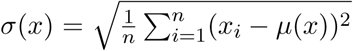. For a vector *x* ∈ ℝ*^p^*, let *S*(*x*) denote the support of *x*; that is, *S*(*x*) = {*i* : *x*_*i*_ ≠ 0} ⊂ [*p*].

Given a vector *x* ∈ ℝ*^p^* and a parameter *β* ∈ ℝ, we define the soft-thresholding operator *T*_*β*_(*x*): ℝ*^p^* → ℝ*^p^* by

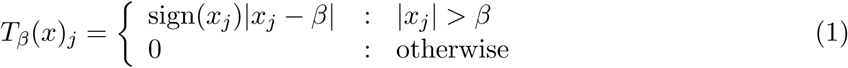

We say that *T*_*β*_(*x*) is a soft-thresholding of the vector *x*.

We use the notation ∥*x*∥*_p_* to represent the *p*-norm of the vector *x*. For example, ∥*x*∥_2_ is the standard Euclidean norm of *x* and 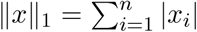. The notation ∥*x*∥_0_ represents the number of nonzero elements in *x*.

### RANKing scRNA-seq data

Consider a vector *x* ∈ ℝ*^n^*. For a given index *i* with 1 ≤ *i* ≤ *n*, let *S*_*i*_(*x*) = {*𝓁* ∈ [*n*] : *x*_*l*_ < *x*_*i*_} and *E*_*i*_(*x*) = {*𝓁* ∈ [*n*] : *x*_*l*_ = *x*_*i*_} (note that *i* ∈ *E*_*i*_(*x*)). Define a transformation Φ: ℝ*^n^* → ℝ*^n^* by

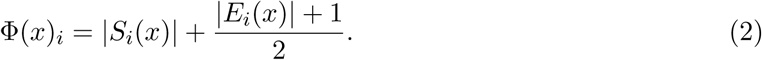

We refer to Φ as the *rank transformation* due to the fact that *Φ*(*x*)*_i_* is the index of *x*_*i*_ in an ordered version of *x*. If multiple elements in *x* are equal, we assign their rank to be the average of the ranks that would be assigned to those elements.

In the definition above, the codomain of Φ is given as ℝ*^n^*. From the defining equation (2), however, we see that each entry of Φ(*x*) (*x* ∈ ℝ*^n^*) is either an integer or a half-integer.

**Example:** Let *n* = 5, and consider the point *x* = (17, 17, 4, 308, 17). Then Φ(*x*) = (3, 3, 1, 5, 3). This value will be the same as the rank transformation applied to any point in *x* ∈ ℝ^5^ with *x*_3_ *< x*_1_ = *x*_2_ = *x*_5_ *< x*_4_.

The rank transformation is non-local, non-linear, and highly distorts the geometry of ℝ*^n^*. There is still much more formal analysis to cover in regards to the connection between the rank transformation and scRNA-seq data; some of this analysis will appear in an upcoming work [14]. Here we provide some intuition as to why the rank transformation produces intelligible results on scRNA-seq UMI counts data.

scRNA-seq data is very sparse and has a high dynamic range. Thus, when looking at the expression of a fixed gene *g* across a population of cells, it is intuitive to separate the cells in which *g* is expressed from the cells in which no expression of *g* is observed. Among the cells in which *g* is observed, it is interesting to distinguish between low expression of *g* and high expression of *g*. The actual values of the counts in cells with high expression (say a count of 500 vs a count of 1000) are often not especially important.

Capturing this idea, methods in the scRNA-seq literature often normalize the counts matrix *X* via a log transformation, taking *X*_*ij*_ ↦ log(*X*_*ij*_ + 1). This is a nonlinear transformation that helps to reduce the gaps between the largest entries of *X* while leaving unchanged the entries that were originally 0 (and preserving much of the gap between “no expression” and “some expression”). With this in mind, the rank transformation can be considered as a more aggressive log transformation. Under the rank transformation, the largest count will be brought adjacent to the second-largest - no gap will be preserved. On the other hand, since there are so many entries that are 0, the gap between no expression (a count of 0) and some expression will be significantly expanded (in the equation (2), the set *E*_*i*_(*x*) will be large for any *i* such that *x*_*i*_ = 0). See Figure 1 for a visualization of these ideas on experimental scRNA-seq data.

**Figure 1:**
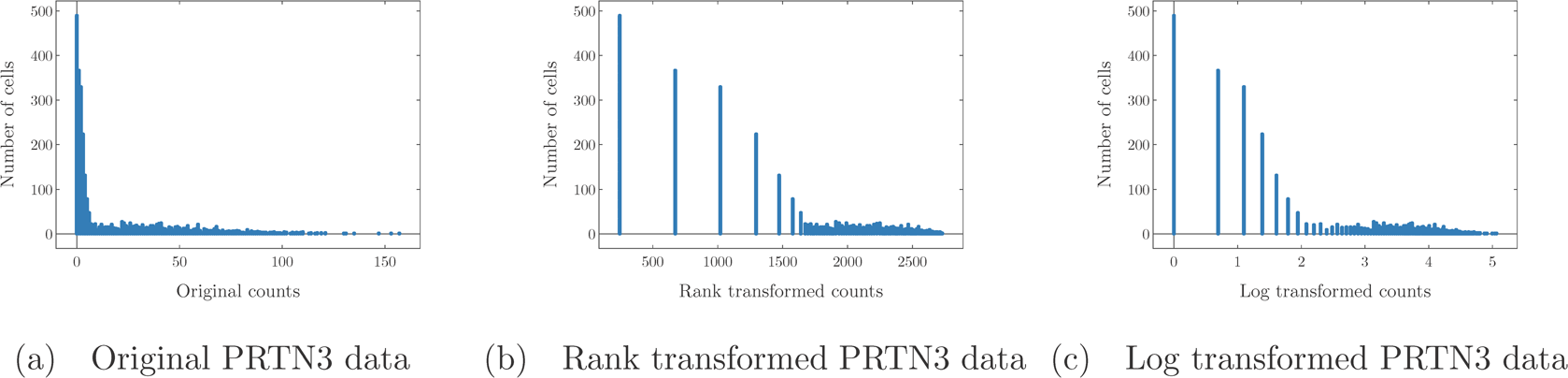
Counts of gene PRTN3 in bone marrow cells in the Paul data set (See the Methods). Each point represents a cell; the horizontal axis shows the number of reads and the vertical axis counts the number of cells with a fixed number of reads. No library size or cell size normalization has been carried out in these pictures in order to facilitate a comparison of the methods depicted herein. Note that the tail of the log transformed data is subjectively longer, while the gap between zero counts and nonzero counts appears larger in the rank transformed data

Stratifying the gene expression in this way intuitively seems useful for determining which genes are important in identifying cell types: a gene that shows expression in many of the cells of a given cell type can be used to separate that cell type from all of the others and thus is a useful marker gene. Thus, by enforcing a large separation between expression and no expression (when compared to the separation between low expression and high expression), it will be easier to identify markers. For these reasons, and since the rank transformation has shown promise in other scRNA-seq tools (for example NODES [15]) we have decided to involve the rank transform in the creation of the RANKCORR marker selection algorithm.

### Mathematical development of a marker selection algorithm

Let *X* ∈ **R**^*n*×*p*^ be a scRNA-seq count matrix (*n* cells, *p* genes). Label the cells with the numbers in [*n*]. Given a subset *S* ⊂ [*n*] of cells of a specific cell type, define *τ* ∈ {±1}*^n^* such that *τ*_*i*_ = +1 if cell *i* is in the subset corresponding to the cell type (that is, if *i* is in *S*) and *τ*_*i*_ = −1 otherwise. If there is a vector *ω* ∈ ℝ*^p^* such that

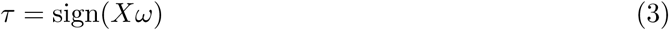

then *ω* gives us a linear separator between the cells that are in the subtype *S* and the cells that are not in *S* (if cell *i* is in *S*, then ⟨*x*_*i*_, *ω* ⟩ *>* 0; otherwise, ⟨*x*_*i*_, *ω*⟩ < 0). In this case, the nonzero entries of *ω* are marker genes for the type *S* - they are the features that separate the given cell type from the other cells.

Ideally, we would like to only select a small number of marker genes. To account for this, we enforce sparsity in *ω*. This corresponds to seeking a sparse linear separator between the two classes: we seek as few genes as possible that are “responsible” for the separation between the subset and the other cells. Unfortunately, it is computationally infeasible to find an optimal sparse separating hyperplane *ω* (see [16]).

In [17], the authors present the convex optimization problem (4) that uses *X, τ* and an input sparsity parameter *s* to give a “good” (in a technical sense) approximation 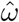 to the true sparse *ω* (assuming that *ω* exists).

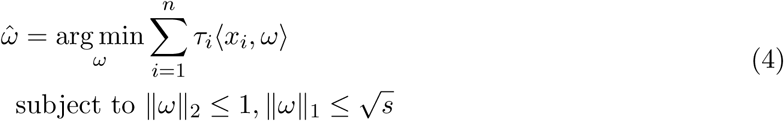

The sparsity parameter *s* controls the size of the set that the approximation 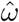 will be chosen from. In particular, if a true signal *ω* has ∥*ω*∥_0_ ≤ *s*, then *ω* will be in the feasible range of the optimization when the input sparsity parameter is *s* or larger. In practice, larger values of *s* will result in larger number of features that are selected.

In both [18] and [19], the authors show that the solution 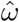 to (4) is given by a normalized soft thresholding of the vector

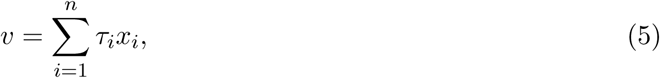

where *x*_*i*_ represents the *i*-th row of *X*. That is,

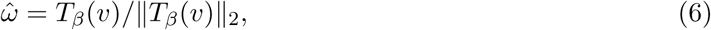

where *β* is a parameter that depends on *s*. Due to this result, it is possible to very quickly find the solution 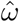 to the optimization (4) (see the Select algorithm in the Methods), making the approximation 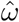 from (4) an appealing method to use for the recovery of *ω* when dealing with large data sets (though other approximation algorithms exist in the literature; e.g. [19], [20], [21]).

The results of [17] (and its extension [22]) depend on the assumption that the rows of *X* are independently generated from a (sub-)Gaussian distribution with mean 0 and variance 1. The rows of an scRNA-seq data matrix *X* do not look like standard Gaussian vectors and, importantly, do not have mean 0. In order to apply the results of [17] to sparse biological data, therefore, the authors of [13] propose standardizing the columns of *X* before running the optimization (4). That is, they create *X*^*std*^ by 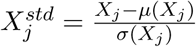 and then let 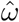 be the solution generated from *X*^*std*^ and *τ*. Note that, in this case, the vector *v* from (5) is defined by

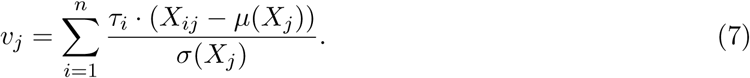

That is, the *j*-th entry of the vector *v* is something that looks a lot like the empirical Pearson correlation between the vector *τ* and column *x*_*j*_ (in fact, *v*_*j*_ is proportional to this correlation). Thus, when we soft threshold this vector *v* from (7), the remaining nonzero entries correspond to the genes that have the (absolute) largest correlation with the vector of cluster labels *τ* .

More accurately, the method in [13] uses a quasi-standardization of the matrix *X* that depends on two hyperparameters. The specific details are not important here; it is enough to establish that [13] proposes a feature selection algorithm for sparse biological data that works by solving the optimization 4 using a quasi-standardized version of *X*. We refer to this algorithm as Spa for the remainder of this work; the markers selected by Spa are those that have high correlation with the cell type labels. As discussed in the introduction, the method that we introduce in this work is inspired by Spa.

## Results and Discussion

### RANKCORR

A fast feature selection algorithm using the rank transformation

RANKCORR, a new method for marker selection based on the rank transformation, is based on the ideas presented in [13]. It essentially works by applying the rank transformation to the columns (genes) of the data matrix as well as the vector *τ* ∈ {±1}*^n^* of class labels and then solving the optimization (4) on the rank transformed data. When compared to the quasi-standardization of genes presented in [13], RANKCORR has the benefit that there are no hyperparameters to tune. In addition, all normalization is done via the rank transformation - there are no extra normalization steps to consider. In experiments, we see that RANKCORR runs much more quickly than the method Spa and generally produces better results. See the Methods for the full details of the algorithm.

In the Methods, we also present a fast algorithm called Select that allows us to quickly jump to the solution of the optimization (4) without the use of optimization software. We use this algorithm in our implementations of both Spa and RANKCORR. Select is the main reason that RANKCORR is a fast algorithm, as it prevents us from needing a general optimization package. This also means that our implementation of Spa is faster than it would have appeared in past work, including [13] (where Spa is introduced).

It can be shown (see the Methods) that RANKCORR will select the genes that have the highest (absolute) Spearman correlation with the vector of class labels. Note that choosing the genes that have the highest absolute Spearman correlation with the vector *τ* could be carried out in a greedy fashion without solving the optimization (4) (similar to a differential expression method) - for this reason, it is not useful to apply RANKCORR to a data set that contains only two cell types. The sparsity parameter *s* does not directly control the number of markers that are selected, however: for example, for a fixed parameter *s* and fixed data matrix *X*, changing the vector *τ* will change the number of markers that are selected. Thus, fixing *s* across all clusters allows for nontrival multi-class marker selection to be carried out in a one-vs-all (OvA) approach.

The effect of fixing *s* is complex; refer to upcoming work [14] for a full description of how this works. In the following, we try to give a bit of intuition. We refer to the norm of the Spearman rank correlation between a gene and the vector *τ*_*j*_ ∈ {±1}*^n^* of cluster labels for cluster *j* as a gene’s score for cluster *j*.

When fixing the value of *s* while choosing markers for each cluster, RANKCORR will select at least *s* markers for each cluster. The RANKCORR algorithm will select exactly *s* markers for cluster *j* only if there are exactly *s* genes tied with the same (absolute) highest score for cluster *j*, which will essentially never occur in noisy experimental data. After selecting *s* markers, the algorithm will continue to choose more markers for the cluster; the number of markers selected depends on the gaps between the scores for different genes. If several genes have a similar score for cluster *j* and one of them is selected as a marker by the algorithm, it is more likely that they will all be selected. If, on the other hand, there are large gaps between consecutive scores, then fewer markers will be selected.

So, if a cluster *j* can be separated from the other clusters using a small number of marker genes, we would expect for these marker genes to exhibit high scores for cluster *j* compared to the rest of the genes. In this case, RANKCORR will select a subset of these high score genes, and maybe a few more depending on the value of *s*. Issues can occur if *s* is selected to be too large, but this is not a problem that we will address here. On the other hand, if cluster *j* is hard to separate from the other cells we would expect it not to exhibit any particularly high scoring genes, and we would see more of a linear continuum of scores (without any large gaps between scores). In this case, more genes will be selected. This allows for the researcher to set a baseline number of markers to select for each cluster, while providing room for RANKCORR to select more genes depending on the distribution of scores for the cluster.

As mentioned above, this discussion was meant to be intuitive and does not formally cover all of the possible cases. To summarize the main points: in RANKCORR, we solve the optimization (4) for each of the cell types in the data set, using rank transformed UMI counts and the same value of *s* for each of the cell types. Thus we may get a different number of markers for each cell type. In this way, we don’t need to specify the number of markers that we would like to obtain for each cell type - we will instead select an informative number for each cluster. Please refer to the Methods or the upcoming work [14] for more information.

### Comparison and benchmarking of marker selection methods in scRNA-seq data

#### Marker set evaluation metrics

There is currently no definitive ground truth set of markers for any experimental scRNA-seq data set. Known markers for cell types have usually been determined from bulk samples, and treating these as ground truth markers neglects the individual cell resolution of single cell sequencing. Moreover, we would argue that the set of known markers is incomplete and that other genes could be used as effectively as (or more effectively than) known markers for many cell types. Indeed, finding new, better markers for rare cell types is one of the coveted promises of single cell sequencing. Since a goal of marker selection is to discover heretofore unknown markers, we cannot evaluate the efficacy of a marker selection algorithm by testing to see if the algorithm recovers a set of previously known markers on experimental scRNA-seq data sets. For this reason, and since our experimental data sets come with an associated clustering into cell types, we evaluate the quality of the selected markers by measuring how much information the selected markers provide about the given clustering.

We examine two general procedures to evaluate how much information a set of selected markers provides about a given clustering when ground truth markers are not known: a supervised classification procedure, in which we train a classifier on the data contained in the selected markers using the ground truth clustering as the target output; and an unsupervised clustering procedure, where we cluster the cells using the information in the set of markers without any reference to the ground truth clustering. Different methods (algorithms) have been developed to accomplish each procedure, and for each method we consider several different performance metrics: a summary of these marker set metrics is found in rightmost column of Table 1, along with the abbreviations that we will use to refer to the metrics. It is important to note that these metrics do not capture the full picture and only represent summary statistical information about the markers that are selected. See the marker evaluation metrics section for further information, including information about how the parameters for the Louvain clustering algorithm are selected.

**Table 1:**
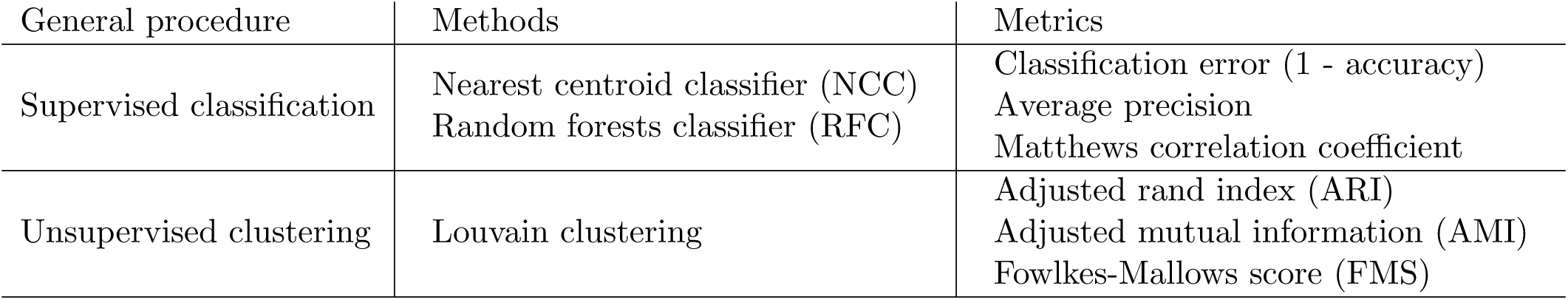
Evaluation metrics for marker sets on experimental data. The “Average precison” metric is a weighted average of precision over the clusters. The Matthews correlation coefficient is a summary statistic that incorporates all information from the confusion matrix. See [23] for more information about the classification metrics and [24] for more information about the clustering metrics.

Following Table 1, for each set of markers that we select, we use the markers to classify the cells twice (using the NCC and the RFC) and to cluster the cells once (using the Louvain clustering algorithm). For both classifications, we examine three evaluation metrics, and we examine an additional three evaluation metrics for the clustering. This results in a total of nine metrics for the evaluation of a set of markers. The three supervised classification metrics (error rate, precision, and Matthew’s correlation coefficient) generally provide similar information, however. Thus, for most data, the results from five metrics (NCC classification error, RFC classification error, ARI, AMI, and FMS) are presented in this manuscript.

We compute each of the metrics using 5-fold cross-validation. Cross-validation is a common technique from the computer science literature that is designed to reduce overfitting - see Section 7.10 of [7]. This means that we always select markers on 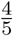 of the cells and then use those markers to classify (or cluster) the other 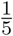 of the cells. The timing information reported in the following sections represents the time needed to select markers on one fold.

#### Summary of marker selection algorithms and experimental data sets

We consider the marker selection methods listed in the leftmost column of Table 2. See the marker selection methods section for implementation details. We apply these methods to some subset of the data sets listed in Table 3; see the experimental data for more information. Each data set in Table 3 comes with a “ground truth” clustering; we use the marker selection methods to determine markers for the clusters and apply the evaluation metrics from Table 1 to evaluate the quality of the selected markers. The large number cells in these data sets reflects the fact that modern scRNA-seq data sets tend to be larger.

**Table 2:** Performance summary of marker selection methods tested in this paper. All designations of “good” performance and “poor” performance are compared to the other methods and are based on observations of graphs presented in later sections of this work. “Class” refers to the supervised classification metrics and “Clust” refers to the unsupervised clustering metrics. We also test the SCDE and D^3^E methods; we find that they are too slow to run on the Paul data set (the smallest data set considered in this work), however. The Spa method does not show any particular highlights and is thus excluded from this table as well.

| Method | ZEISEL | PAUL | ZHENGFI<sup>L</sup>T |
| --- | --- | --- | --- |
| RANKCORR | Class: Good, < 100 markers<br>Clust: Good, < 200 markers | Clust: FMS good, < 100 markers<br>Good, > 100 markers | Class: Good, < 120 markers<br>Clust: Poor, > 200 markers |
| t-test | Class: Good always<br>Clust: Good, > 200 markers | Clust: FMS good, < 100 markers<br>Good, > 100 markers | Class: Good, < 120 markers<br>Clust: Good, > 50 markers |
| Wilcoxon | Clust: Good, > 200 markers | Class: Poor, many markers<br>Clust: AMI good, < 100 markers<br>Good, > 100 markers | Class: Poor, > 50 markers<br>Clust: Good, > 50 markers |
| edgeR<br>edgeRdet | Class: Poor, < 50 markers | Class: edgeR poor, < 80 markers<br>Clust: edgeR good, > 100 markers | only run edgeR<br>Class: Poor, < 50 markers<br>Good, > 100 markers<br>Clust: Good, ∼ 20 markers |
| MAST<br>MASTdet | Class: Good, < 100 markers<br>Clust: Good, < 50 markers<br>Good, > 200 markers | Clust: MAST good, > 100 markers<br>MASTdet poor, > 100 markers | only run MAST<br>Class: Good, < 120 markers |
| scVI | Class: Poor always<br>Clust: Good, between 100 and 200 | Clust: Poor always<br>Class: Poor always | Did not run;<br>requires too much RAM |
| Elastic Nets | Class: Poor, < 50 markers<br>Clust: Good, between 100 and 200 | Class: Poor, < 80 markers<br>Good, between 100 and 400<br>Clust: Poor, > 100 markers | Did not run;<br>too slow |
| Log. Reg. | Class: Poor for < 50 markers<br>Good, > 150 markers<br>Clust: Poor always | Class: Good, between 80 and 250<br>Clust: Poor, > 100 markers | Class: Poor, < 150 markers<br>Clust: Poor always |

**Table 3:** Data sets considered in this work. See experimental data for more information. ZhengFilt contains a subset of the data in ZhengFull. ZhengSim is a collection of simulated data sets created with the Splatter R package; see the generating synthetic data section in the Methods for more information.

| Data set | cells | non-zero genes |
| --- | --- | --- |
| ZEISEL | 3005 | 4999 |
| PAUL | 2730 | 3451 |
| ZHENGFI<sup>L</sup>T | 68579 | 5000 |
| ZHENGFULL | 68579 | 20387 |
| 10xMOUSE | 1.3 million | 24015 |
| ZHENGSI<sup>M</sup> | 5000 | varies |

A summary of the performance of the methods is included in Table 2. This table notes areas where the methods perform noticeably better or worse than the majority of the other methods according to the data that is graphically presented in later sections. It is important to emphasize that the performances of the different methods are generally quite similar; thus, the annotations in Table 2 are quite specific and do not cover all possible sizes of marker sets for all metrics. The configurations that aren’t mentioned (e.g. the classification metrics on the Paul data set) indicate that the algorithms perform similarly in those regions.

An important note is that the relative speeds of the algorithms are not indicated in Table 2. Since the algorithms generally exhibit similar performances under the metrics considered in this work, fast algorithms have a significant advantage over the other algorithms. The fastest methods are RANKCORR, the t-test, and Wilcoxon: see Tables 4, 5, 6, and 7 for this timing information. It is thus these three methods that show a clear advantage over the other methods for working with future experimental data. Logistic regression also runs quickly on the smaller data sets, but does not scale as well as the three methods mentioned above, and significantly slows down on the larger data sets. In addition, logistic regression shows inconsistent performance, and is often one of the worst performers when selecting small numbers of markers. All of the other methods are significantly slower or require large computational resources compared to the size of the data set.

**Table 4:**
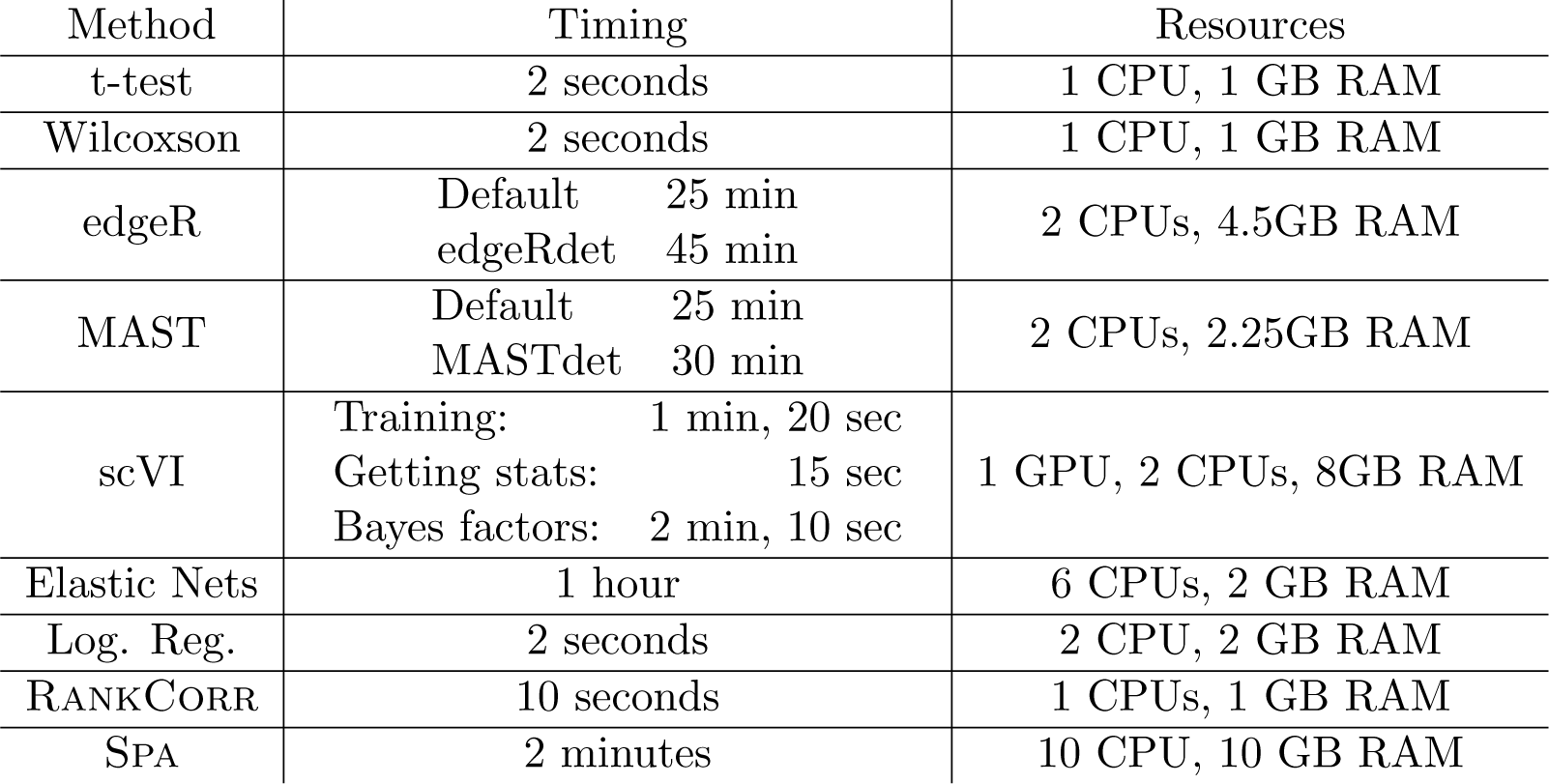
Timing and computation resources required on the Zeisel data set. Times are reported to compute the markers for one fold and are approximate.

| Method | Timing |  | Resources |
| --- | --- | --- | --- |
| t-test | 2 seconds |  | 1 CPU, 1 GB RAM |
| Wilcoxon | 2 seconds |  | 1 CPU, 1 GB RAM |
| edgeR | Default | 25 min | 2 CPUs, 4.5GB RAM |
|  | edgeRdet | 45 min |  |
| MAST | Default | 25 min | 2 CPUs, 2.25GB RAM |
|  | MASTdet | 30 min |  |
| scVI | Training: | 1 min, 20 sec | 1 GPU, 2 CPUs, 8GB RAM |
|  | Getting stats: | 15 sec |  |
|  | Bayes factors: | 2 min, 10 sec |  |
| Elastic Nets | 1 hour |  | 6 CPUs, 2 GB RAM |
| Log. Reg. | 2 seconds |  | 2 CPU, 2 GB RAM |
| RANKCORR | 10 seconds |  | 1 CPUs, 1 GB RAM |
| SPA | 2 minutes |  | 10 CPU, 10 GB RAM |

**Table 5:** Timing and computation resources required on the Paul data set. Apart from D^3^E, times are reported to compute the markers for one fold and are approximate. The reported time for D^3^E is for one *cluster*. There are 19 clusters in the Paul dataset, and we would need to collect markers for all 19 clusters in each of the 5 folds to perform the full cross validation analysis.

| Method | Timing | Resources |
| --- | --- | --- |
| t-test | 2 seconds | 1 CPU, 1 GB RAM |
| Wilcoxon | 3 seconds | 1 CPU, 1 GB RAM |
| edgeR | Default: 40 min<br>edgeRdet: 1 hour 5 min | Default: 1 CPU, 6 GB RAM<br>edgeRdet: 1 CPU, 5 GB RAM |
| MAST | Default: 35 min<br>MASTdet: 40 min | 1 CPU, 2 GB RAM |
| scVI | Training: 1 min, 20 sec<br>Getting stats: 10 sec<br>Bayes factors: 2 min, 50 sec | 1 GPU, 2 CPUs, 8GB RAM |
| D<sup>3</sup>E | 25 min for one cluster* | 10 CPUs, 2GB RAM |
| Elastic Nets | 55 minutes | 2 CPUs, 1 GB RAM |
| Log. Reg. | 10 seconds | 1 CPU, 1 GB RAM |
| RANKCORR | 10 seconds | 1 CPU, 1 GB RAM |
| SPA | 1.5 minutes | 10 CPUs, 5.5 GB RAM |

**Table 6:** Timing and computation resources required on the ZhengFilt data set. Times are reported to compute the markers for one fold and are approximate. The Logistic Regression method did not converge on every cluster in some of the folds. The memory requirements for these data are overestimates.

| Method | Timing | Resources |
| --- | --- | --- |
| t-test | 15 seconds | 1 CPU, 4 GB RAM |
| Wilcoxon | 30 seconds | 1 CPU, 8 GB RAM |
| Log. Reg. | 10 minutes | 3 CPUs, 8 GB RAM |
| edgeR | 16 hours | 1 CPU, 80 GB RAM |
| MAST | 10 hours | 1 CPU, 30 GB RAM |
| RANKCORR | 2 minutes | 1 CPU, 8 GB RAM |

**Table 7:** Timing and computation resources required on the ZhengFull data set. Times are reported to compute the markers for one fold and are approximate. The Logistic Regression method did not converge on every fold. The memory requirements for these data are overestimates.

| Method | Timing | Resources |
| --- | --- | --- |
| t-test | 2.5 minutes | 1 CPU, 8 GB RAM |
| Wilcoxon | 3 minutes | 1 CPU, 8 GB RAM |
| Log. Reg. | 30 minutes | 4 CPU, 8 GB RAM |
| RANKCORR | 12 minutes | 2 CPUs, 8 GB RAM |

The ZhengFull and 10xMouse data sets do not appear in Table 2. These data sets were too large for the majority of the methods to handle; thus, data was collected only for the RANKCORR, t-test, logistic regression, and Wilcoxon methods (the fastest methods). We include the 10xMouse data specifically as a stress test for the methods to see which could handle the largest data sets. It is impressive that these methods are able to run on such a large data set in a reasonable amount of time. The ZhengSim data also does not appear in the table, partially due to the aforementioned speed issues and partially due to the fact that we can examine different evaluation metrics when we know a ground truth set of markers.

### The marker selection methods perform well on the well-clustered Zeisel data set

The Zeisel data set contains 9 clusters that are generally well separated (since neuronal cells are usually fully differentiated). Thus, we expect that both the classification metrics and the clustering metrics should be quite good for all methods. The Zeisel data set, in some ways, acts as a biological verification of the marker selection methods.

The time and computational resources required for the marker selection methods on Zeisel are shown in Table 4. The t-test, Wilcoxson, logistic regression, and RANKCORR methods all run quickly and require few resources. The MAST algorithms, the edgeR algorithms, elastic nets, and Spa all take a significant amount of time on this small data set. scVI runs quite quickly, but requires steep computational resources (a GPU and 8GB of RAM for a small data set). All of the marker selection methods perform quite well on the Zeisel data set; thus, the faster methods are preferable here.

#### Supervised classification

The classification error rate (1 - accuracy) of the nearest centroid and random forests classifiers on the Zeisel data set are presented in Figure 2. As expected, the error rates are very low: it requires only 100 markers (an average of 11 markers per cluster) to reach an error rate lower than 5% for most methods using the random forests classifier. Related to the high accuracy, the average precisions are also very high and the precision curves do not provide new information. The same is true of the Matthews correlation coefficient curves. Thus, both the precision and Matthews correlation curves are not presented here: they can be found in Additional File 1, Figures 1 and 2 (for the NCC and RFC data respectively). Selecting random markers produced poor results, so those curves are also omitted from the plots displayed here; the marker selection methods all perform better than random on the Zeisel data set. See Additional File 1, Figures 1 and 2 for random marker selection data.

**Figure 2:**
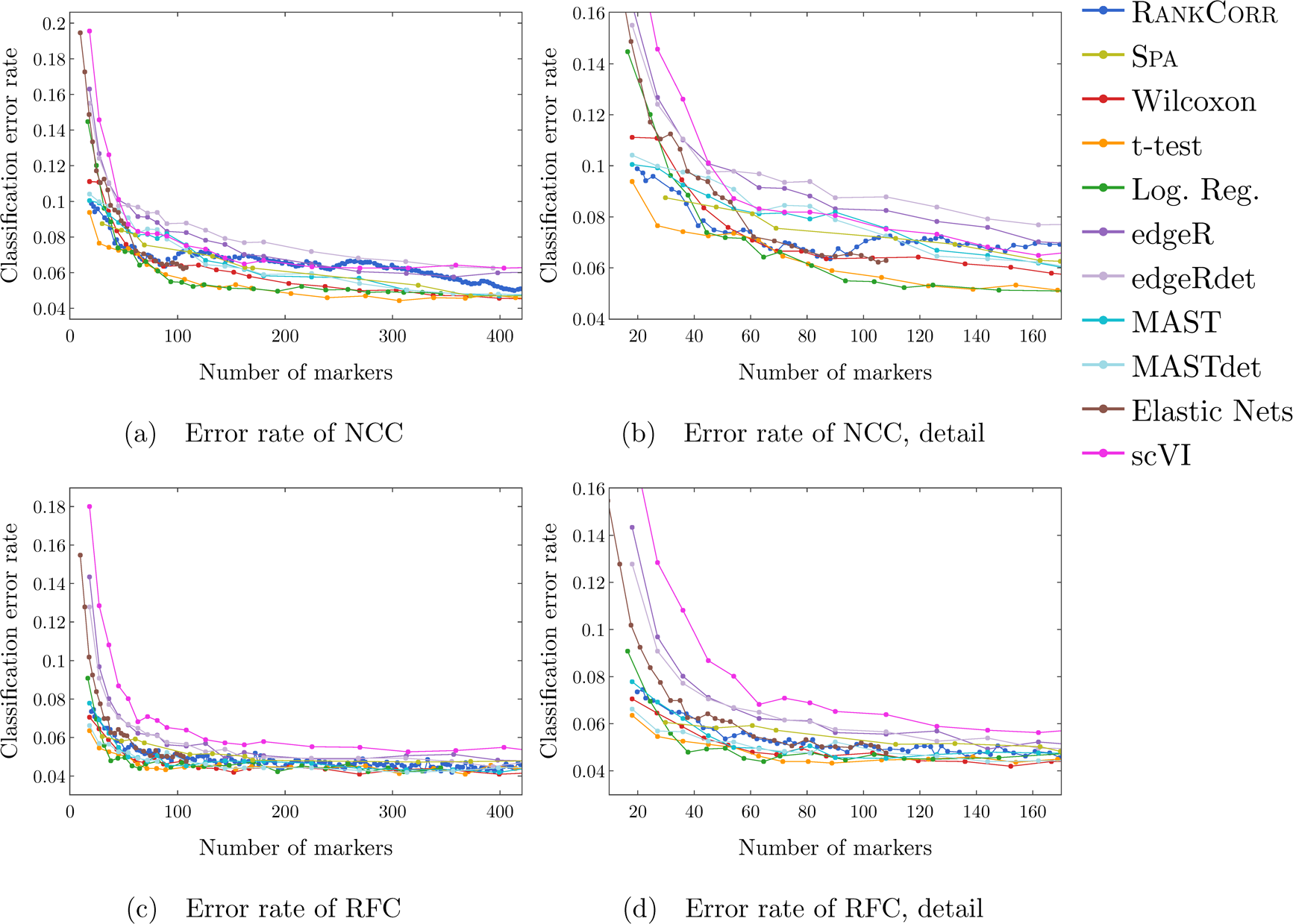
Error rate of both the nearest centroids classifier (NCC; (a) and (b)) and the random forests classifier (RFC; (c) and (d)) on the Zeisel data set. Figure (b) (respectively (d)) is a detailed image of the error rate of the different methods using the NCC (respectively RFC) when smaller numbers of markers are selected.

Examining the performance of the NCC (Figures 2(a) and 2(b)), we see that the t-test, RANKCORR, and the MAST methods are the best methods when selecting small numbers of markers (an average of 2-3 markers per cluster; less than 30 total markers selected). In this same domain, both edgeR algorithms, as well as logistic regression, elastic nets, and scVI all perform the worst, with error rates of around 15% or higher. The error rates of both the logistic regression and elastic nets methods quickly drop: when selecting about 30 to 100 total markers, the best methods are RANKCORR, the t-test, logistic regression, Wilcoxon, and elastic nets. Only a small number of markers were selected by elastic nets on the Zeisel data set; thus, the elastic nets curve ends before the others. As higher numbers of markers are selected, the RANKCORR method moves a bit (up to 3%) away from the best methods, but then catches up with the best methods for very large numbers of markers. The t-test and logistic regression are consistently the highest performing, while both edgeR methods and scVI are consistently the lowest performing, with scVI slightly besting the edgeR methods.

For the RFC (Figures 2(c) and 2(d)), apart from scVI, all of the methods show very similar performance after 150 markers have been selected. scVI stands out from the other methods as the worst across for all levels of selected markers. Selecting less than 30 total markers, the elastic nets, edgeR, and edgeRdet methods also show worse performance than the other methods; the other methods are generally within 1.5% of each other in this domain. The two edgeR methods remain slightly worse than the other methods until about 120 total markers are selected, though edgeR always performs better than scVI according to the RFC (this behavior is flipped from the NCC). RANKCORR is always within 1% of the best method. Importantly, when only examining approximately 2 markers per cluster, RANKCORR produces a low error rate.

Based on these plots, the most pertinent conclusion is that most of the methods tested here produce markers that provide a significant amount of information about the ground truth clustering associated with the Zeisel data set. It is notable that the two edgeR algorithms, scVI, elastic nets, and (to a certain extent) logistic regression methods perform significantly worse than the other methods when selecting small numbers of markers according to these classification metrics (though it is tough to say that any of the methods perform poorly, since all error rates are quite low). On the other hand, the RANKCORR and t-test methods, along with the two MAST algorithms, are probably the best choices for selecting small numbers of markers on the Zeisel data set according to these data. Both of the edgeR algorithms and scVI generally perform suboptimally on the Zeisel data set overall. It is tough to say that any method is best for large numbers of markers, though the t-test and logistic regression methods look best for the NCC.

Most importantly, these data suggest that when examining a data set that is well clustered, it is useful to examine several marker selection algorithms to get different perspectives on which genes are most important. This provides an advantage for marker selection algorithms that can run using only small amounts of resources.

#### Unsupervised clustering

The ARI, AMI, and FM scores, reported in Figure 3, are high for all methods, as expected. The three plots show similar characteristics, so we analyze them together here. All of the methods perform much better than random marker selection: see Additional File 1, Figure 3 for the performance of random marker selection under the unsupervised clustering metrics.

**Figure 3:**
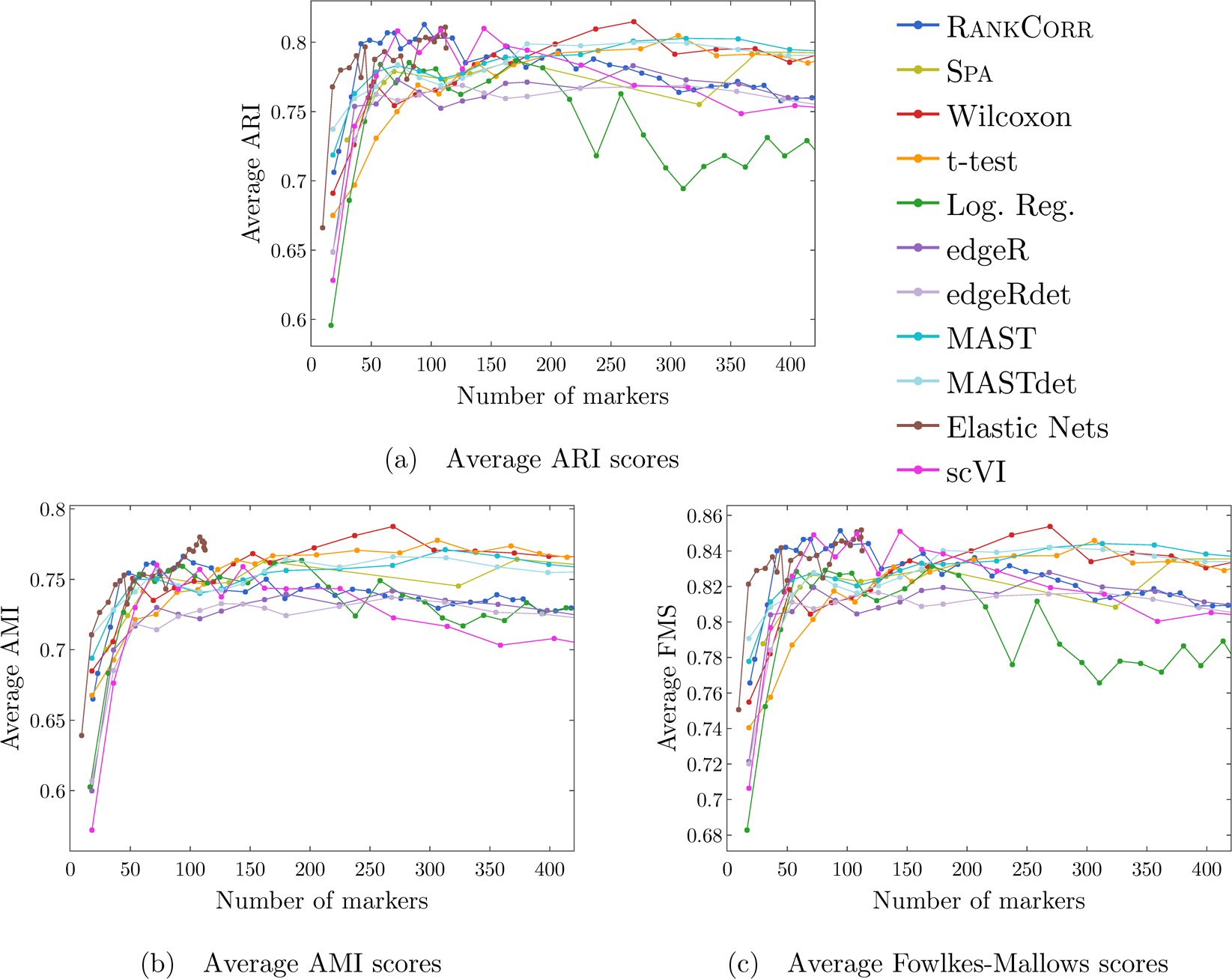
Clustering performance metrics vs total number of markers selected for marker selection methods on the Zeisel data set. The ARI score is shown in (a), the AMI score is shown in (b), and the Fowlkes-Mallows score is shown in (c). The clustering is carried out using 5-fold cross validation and scores are averaged across folds.

When less than 50 markers are selected (about 5 unique markers per cluster), the elastic nets, MAST, MASTdet, and RANKCORR methods perform significantly better than the other methods, with scores often about 0.05 to 0.1 higher than the other methods in this range. As seen in the classification metrics, the two edgeR algorithms, logistic regression, and scVI perform poorly in this range. Unlike the classification metrics, the t-test shows low ARI and FM scores for less than 50 markers selected as well.

Then, when between 50 and 150 total markers are selected, the RANKCORR, Elastic Nets, and scVI methods perform at the highest levels. All of the other methods exhibit similar performance in this regime, though the edgeR methods continue to show the worst performance. The t-test and Wilcoxon methods produce low scores when 50 total markers are selected but steadily improve through this range until they rival the (optimal) performance of scVI and RANKCORR when approximately 150 total markers are selected.

At around 200 markers selected, the methods split into two groups that are separated by about to 0.05 depending on the score. The lower performing band consists of the RANKCORR, scVI, edgeR, and edgeRdet methods, while the higher performing band consists of the Wilcoxon, t-test, MAST, and MASTdet methods. Elastic Nets selects less than 200 markers on this data set, so it is not present in either of these bands. Logistic regression falls into the lower group on the AMI curve but performs significantly worse than all of the other methods in the ARI and FM score plots. Spa starts in the lower group and moves into the upper group when around 300 to 350 total markers are selected; it is thus possible that a small number of genes are responsible for the separation between the methods here.

It is noteworthy that logistic regression method is one of the worst methods when large numbers of markers are selected. In the supervised classification trials, the logistic regression method was one of the top performers when selecting large numbers of markers, so this is a striking change. When no information about the ground truth clustering is provided, the performance of logistic regression method on the Zeisel data sets drops considerably. On the other hand, any gaps between the scVI curve and the other curves are eliminated in this unsupervised clustering trial. In fact, when between 100 and 200 markers are selected, the scVI method exhibits the highest ARI and FM score of any of the methods. This is also a reversal from the classification metrics, where scVI was consistently one of the worst methods.

Interestingly, both the RANKCORR and scVI methods hit a peak performance when selecting between 50 and 200 markers, then their performance decreases as more markers are selected. On both the ARI and FM score curves, the optimal scores for RANKCORR and scVI are as high as the top scores for any of the other methods, even though both RANKCORR and scVI move into the lower performing group as more markers are selected. The other two methods in the lower performing group, edgeR and edgeRdet, consistently show some of the worst performance for all sizes of selected marker sets. It is thus unclear what causes this overfitting.

Again, all of the marker selection methods perform well on the Zeisel data set. The separation of performance between the methods for high levels of markers selected, and the fact that Spa quickly jumps between the two bands suggests that there are some important markers that are not selected by all of the methods. This further emphasizes that researchers should compare the results of multiple marker selection algorithms on their data sets, so fast marker selection algorithms are better.

## Discussion

RANKCORR is the only method that shows high performance for small numbers of markers in both the clustering and classification metrics. Most researchers will be looking for small numbers of markers for their data sets; thus RANKCORR stands out as a promising method on the Zeisel data set. Note also that RANKCORR clearly outperforms Spa in the clustering metrics and is competitive with Spa in the classification metrics: RANKCORR is both faster than Spa and selects a generally more informative set of markers than Spa on the Zeisel data set. Therefore, the performance on the Zeisel data set is evidence for the fact that RANKCORR is a progression of Spa for sparse UMI counts scRNA-seq data.

For large numbers of markers on the Zeisel data set, the t-test stands out, since it is in the top performing band in the clustering metric plots and one of the best methods for the NCC classification metric plots. Examining the other statistical and machine learning methods: Wilcoxon also generally does quite well on the Zeisel data set - it never stands out as one of the best methods, but it also consistently stays relevant. Logistic regression is inconsistent, as it mostly shows poor performance for small numbers of markers selected. For large numbers of markers selected, it shows optimal performance under the supervised classification metrics and poor performance under the unsupervised clustering metrics. Elastic nets shows good performance for between 50 and 100 total markers selected using all metrics, but inconsistent performance for less than 50 markers selected.

The other marker selection methods are designed for (sc)RNA-seq data and show a variety of performance levels. The two edgeR methods are consistently some on the lowest performing for the Zeisel data set. On the other hand, the two MAST methods are generally some of the better performers, consistently exhibiting good scores when lower numbers of markers are selected and landing in the top tier of methods for large numbers of markers selected under the clustering metrics. Like edgeR, scVI generally shows worse performance than the other methods, except that it tied for the best method for low numbers of markers selected under the clustering metrics. The main issue with these methods, however, is that they all require significantly higher resources (either time or computational) than other methods that exhibit higher scores.

### Marker selection algorithms struggle with the cell types defined along the cell differentiation trajectory in the Paul data set

The Paul data set consists of bone marrow cells and contains 19 clusters (some of which are very small). Since the clusters lie along a cell differentiation trajectory, it is reasonable that it would be difficult to separate the clusters or to accurately reproduce the clustering. The Paul data set thus represents an adversarial example for these marker selection algorithms, and we expect the methods to show significantly lower scores than the ones that were produced on the Zeisel data set.

The time and computational resources required for the marker selection methods on Paul are shown in Table 5. As Paul is a data set that is similarly sized to Zeisel, the timing information is similar. In particular, the t-test, Wilcoxson, logistic regression, and RANKCORR methods all run quickly and require few resources. The other algorithms all either take a significant amount of time on this small data set or require high computational resources. We additionally implemented the D^3^E method on Paul: it took 25 minutes using 10 CPUs to compute the markers for one cluster (vs all of the others) in one fold. This means that it would take about 21 hours on 10 cores to compute the data for the entirety of the Paul data set. Paul is meant as a small example of the types of data sets that we are focused on in this work and D^3^E would be too slow to run on larger data sets. We thus did not run all of the computations required for D^3^E on Paul and excluded it from further analysis. See Section the marker selection methods for more details. All of the marker selection methods perform quite similarly on the Paul data set; thus, as with the Zeisel data set, the faster methods are preferable.

#### Supervised classification

Figure 4 shows the performance of marker selection algorithms on the Paul data set as evaluated by the supervised classification metrics. Due to the cell differentiation trajectory in the data set, it is not surprising to see relatively high clustering error rates: the rates are always larger than 30% for the NCC and reach a minimum of around 27% with the RFC. Since there are 19 clusters, this is still much better then classifying the cells at random. In addition, selecting markers at random produces high error rates: the minimum error rates when selecting less than random 550 markers are around 40% for the NCC and around 48% for the RFC. Thus, although the error rates produced by the marker selection algorithms are quite high, they are significantly lower than the error produced by random marker selection. For the sake of clarity, Figure 4 does not contain the curves corresponding to random marker selection: see Additional File 1, Figures 4 and 5 for classification error rate figures that include the random markers.

**Figure 4:**
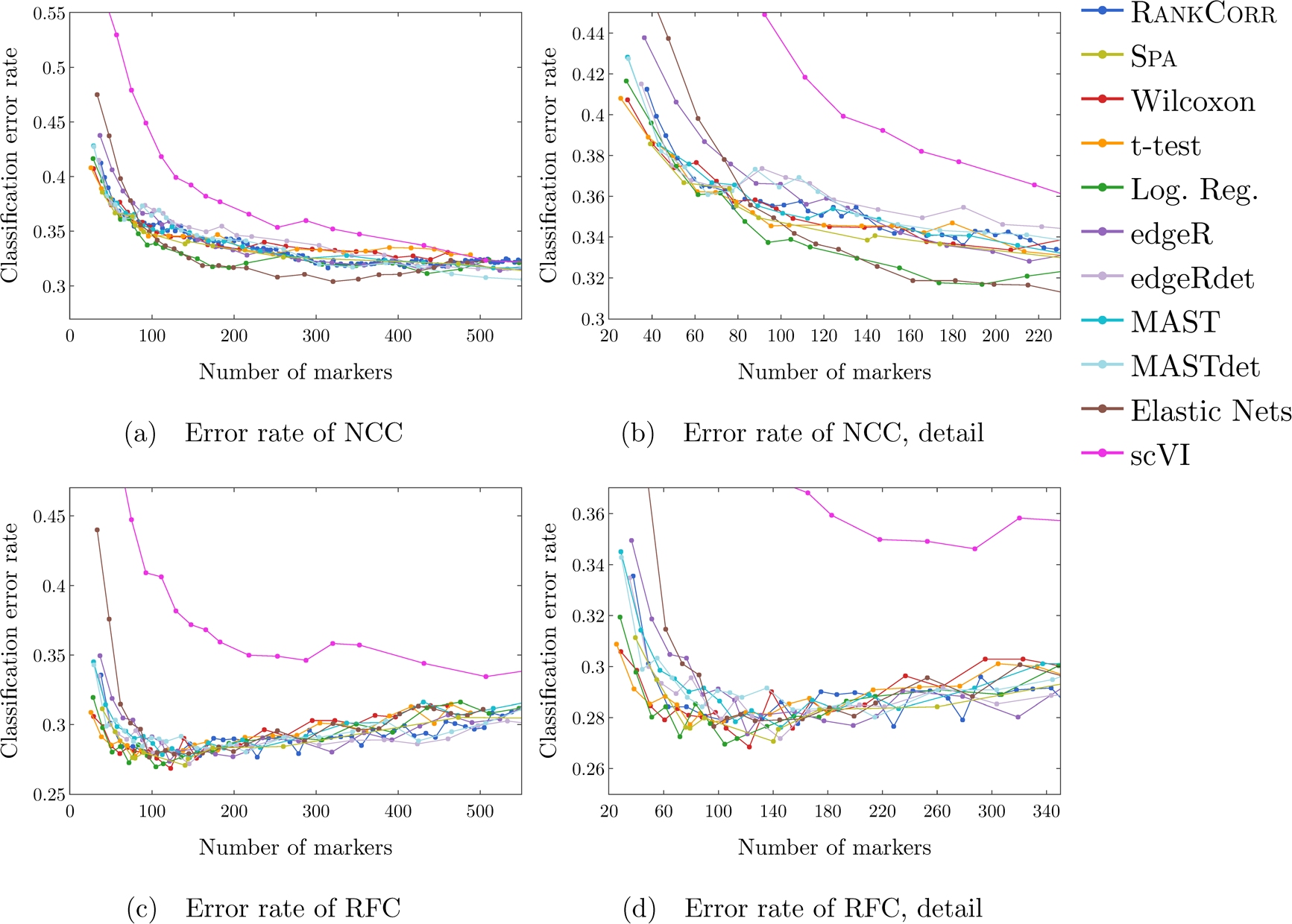
Error rates of both the nearest centroids classifier (NCC; (a) and (b)) and the random forests classifier (RFC; (c) and (d)) on the Paul data set. Figure (b) (respectively (d)) is a detailed image of the error rate of the different methods using the NCC (respectively RFC) when smaller numbers of markers are selected. Figure (b) details up to 220 total markers to make clear how similar the methods perform for when small numbers of markers are selected. Figure (d) examines up to 350 total markers to detail the performance of the methods when small numbers of markers are selected as well as get an idea for the increasing behavior and noisy nature of the curves.

**Figure 5:**
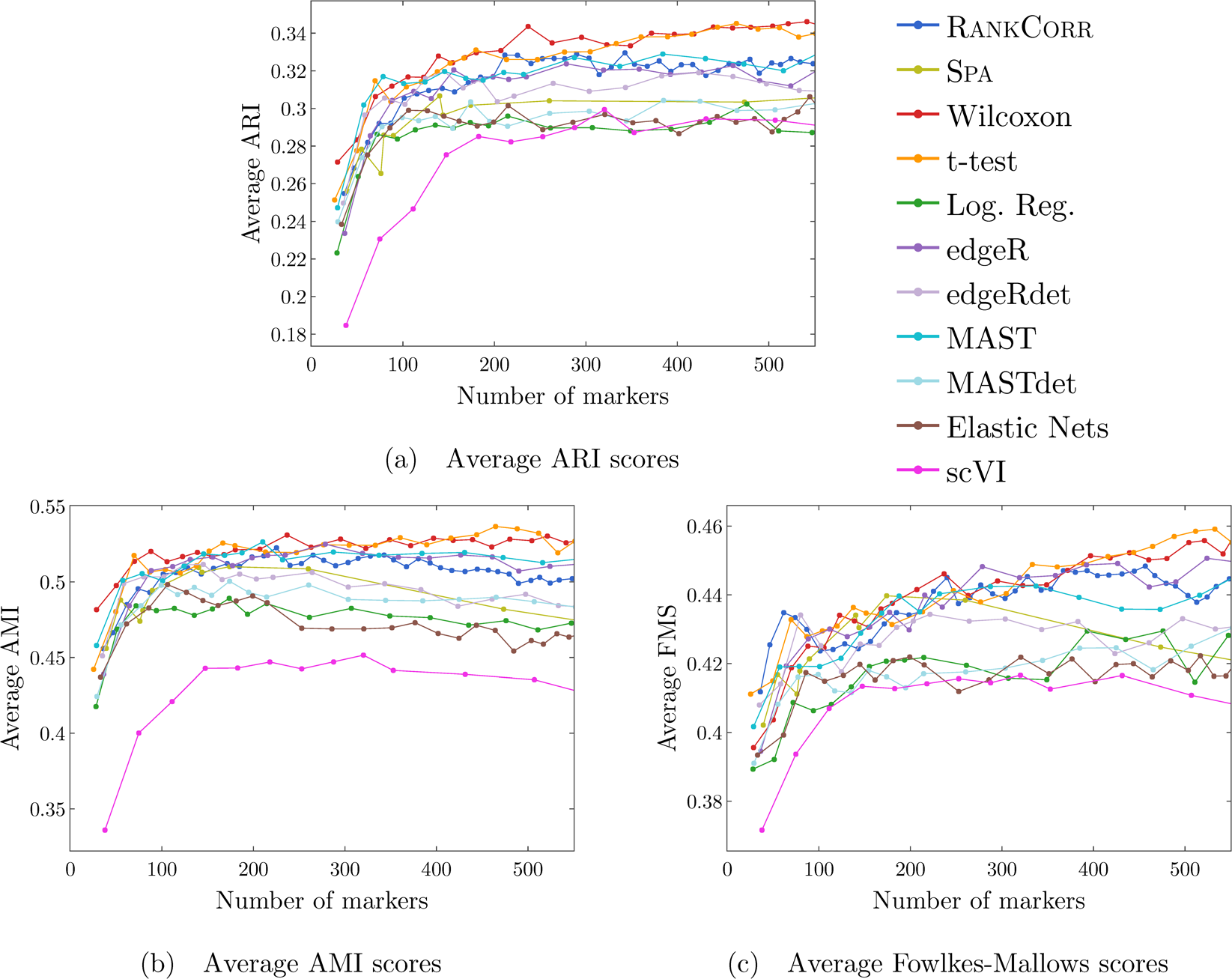
Clustering performance metrics vs total number of markers selected for marker selection methods on the Paul data set. The ARI score is shown in (a), the AMI score is shown in (b), and the Fowlkes-Mallows score is shown in (c). The clustering is carried out using 5-fold cross validation and scores are averaged across folds.

Similar to results on the Zeisel data set, the precision and Matthews correlation coefficient curves do not provide extra information; indeed, it is difficult to distinguish several of the precision and Matthews correlation plots from the classification error rate plots. See Additional File 1, Figures 4 and 5 to view these plots.

We start with a couple of general observations. First, scVI produces significantly higher error rates than the other methods using both the NCC and the RFC. For this reason, in the discussions for the remainder of this section, we will generally ignore scVI and focus on analyzing the other methods. scVI does not perform well on the Paul data set. Secondly, as we increase the number of markers, all of the methods appear to converge to similar performance levels - this is also reasonable, since we will approach the baseline error rate obtained by classifying based on all of the genes as we increase the number of markers.

Next we consider the performance of the methods when classified by the NCC. When selecting less than 80 total markers (an average of approximately 4 markers per clusters), the methods other than elastic nets and edgeR perform quite similarly, with a maximum difference of about 2.5% between those curves (mostly, all of these curves are within 1% of each other). See Figure 4(b). Elastic nets performs worse than edgeR which performs significantly worse than the group of other methods, and all of the error rates are decreasing quickly in this region.

The t-test shows the smallest error rate when a very small number of markers (an average of 2 markers per cluster) are selected, but, again, it is not much better than the other methods. It appears that the t-test gets relatively worse than the other methods as more markers are selected, until it is nearly as bad as scVI when around 400 unique markers are selected. On the other hand, when approximately 80 total markers are selected, the edgeRdet and MASTdet methods show a small spike in error rate. From this point, the edgeRdet and MASTdet error rates continually decrease until they are some of the best methods when around 500 unique markers are selected.

The error rates of the remaining methods (other than edgeRdet and MASTdet) meet at around 80 to 100 total markers selected and then mostly appear to fall into one band. Two methods, logistic regression and elastic nets, show a lower classification error rate for intermediate numbers of markers: logistic regression appears to perform optimally when we are looking for about 80-250 total markers (250 markers translates into an average of around 13 unique markers per cluster, and is obtained when we select 25 marker per cluster using logistic regression). On the other hand, elastic nets overcomes its poor start to perform well for between 100 and 400 total markers (this corresponds to selecting between 6 markers per cluster and 55 markers per cluster for the elastic net metod due to duplicates). RANKCORR performs well when selecting any number of markers, with a difference of less than 3% from the best method at any point along this curve. Since the nearest centroid classifier is quite heuristic, this is overall a small difference.

When we use the random forest classifier (see Figure 4(c)), there is a variety of different performances when less than 80 total markers are selected. Elastic nets performs the worst in this range. EdgeR is better than elastic nets but still worse than the other methods. The Wilcoxon, t-test, and logistic regression methods show the best performance in this regime. All of the other methods (RANKCORR, MAST, MASTdet, edgeRdet, and Spa) show nearly identical performance that is slightly worse than logistic regression. The difference in error rate between the methods in this middle band and the best method is at most 4% and decreases as more markers are selected.

All of the methods then show similar performance when around 100 total markers are selected. Interestingly, the methods appear to perform the best when approximately 100-125 total markers are selected. This suggests some level of overfitting might occur that was not visible in the NCC data.

When more than 140 total markers are selected, all of the methods fall into one group. Many of the curves show high variance without exhibiting clear monotonic behavior; however, the general trend is that the curves are increasing in this range. The the classification error rate of the RFC shows significant variance between classification attempts using the same set of markers; see the discussion about supervised classification in the Methods. Thus, the differences between many of the methods in Figure 4(c) could mostly be due to noise. If anything, it appears that RANKCORR, edgeR, and edgeRdet tend to outperform the other methods as the number of markers increases; Wilcoxon and the t-test again appear to perform the worst (this trend is more pronounced in the RFC precision curve, see Figure 5(c) in Additional File 1) and are joined by logistic regression, MAST, and elastic net when approximately 350 total markers are selected. The maximum difference between the curves in this range is only 3%, however. Therefore, the only definitive conclusion that can be drawn is that all of the methods perform comparably on the Paul data set with respect to the RFC classification metric as larger numbers of markers are selected.

Based on these data, it is difficult to conclude that any method is optimal on the Paul data set. According to the NCC, the methods other than scVI are nearly identical when selecting less than 80 total markers, with logistic regression and the t-test outperforming the other methods when 100-400 total markers selected. On the other hand, the RFC shows that Wilcoxon, the t-test, and logistic regression outperform the other methods when selecting less than 80 markers, with little to no differences between the methods for large numbers of markers selected. These two pictures do not support one another.

On the other hand, there several methods that consistently perform worse on the Paul data set than the majority of the other methods. scVI consistently performs poorly for all amounts of selected markers, while both elastic nets and edgeR show suboptimal performance when less than 80 total markers are selected. Both the NCC and the RFC metrics also suggest that Wilcoxon tends to perform slightly worse than the other methods for large amounts of selected markers. Other than these few exceptions, all of the marker selection methods tested show similar performance under the classification metrics on the Paul data set. This provides further support for the notion that faster, lighter marker selection methods should be preferred over the others, as similar information is obtained from all methods.

A corollary of this is that, on the Paul data set, the RANKCORR method performs comparably to the other methods by the measures that are considered here. It does indeed generally outperform the other methods that have been specifically designed for RNA-seq data.

#### Unsupervised clustering

The dependence of the ARI, AMI, and FM scores on the number of markers selected is plotted for the different marker selection algorithms in Figure 5. There are several features that are common to all three of these plots.

First of all, the values of the scores are all in low to medium ranges for all marker selection algorithms (recall that larger ARI, AMI, and FM scores indicate stronger agreement between the ground truth clustering and the unsupervised clustering; see the Methods). This was expected to some extent, as it should intuitively be difficult to reproduce a clustering that has been assigned along a continuous path. Again, choosing markers randomly performs noticeably worse than all of the marker selection methods and thus the data corresponding to random markers is not plotted in Figure 5 for the sake of clarity. See Additional File 1, Figure 6 for these data.

**Figure 6:**
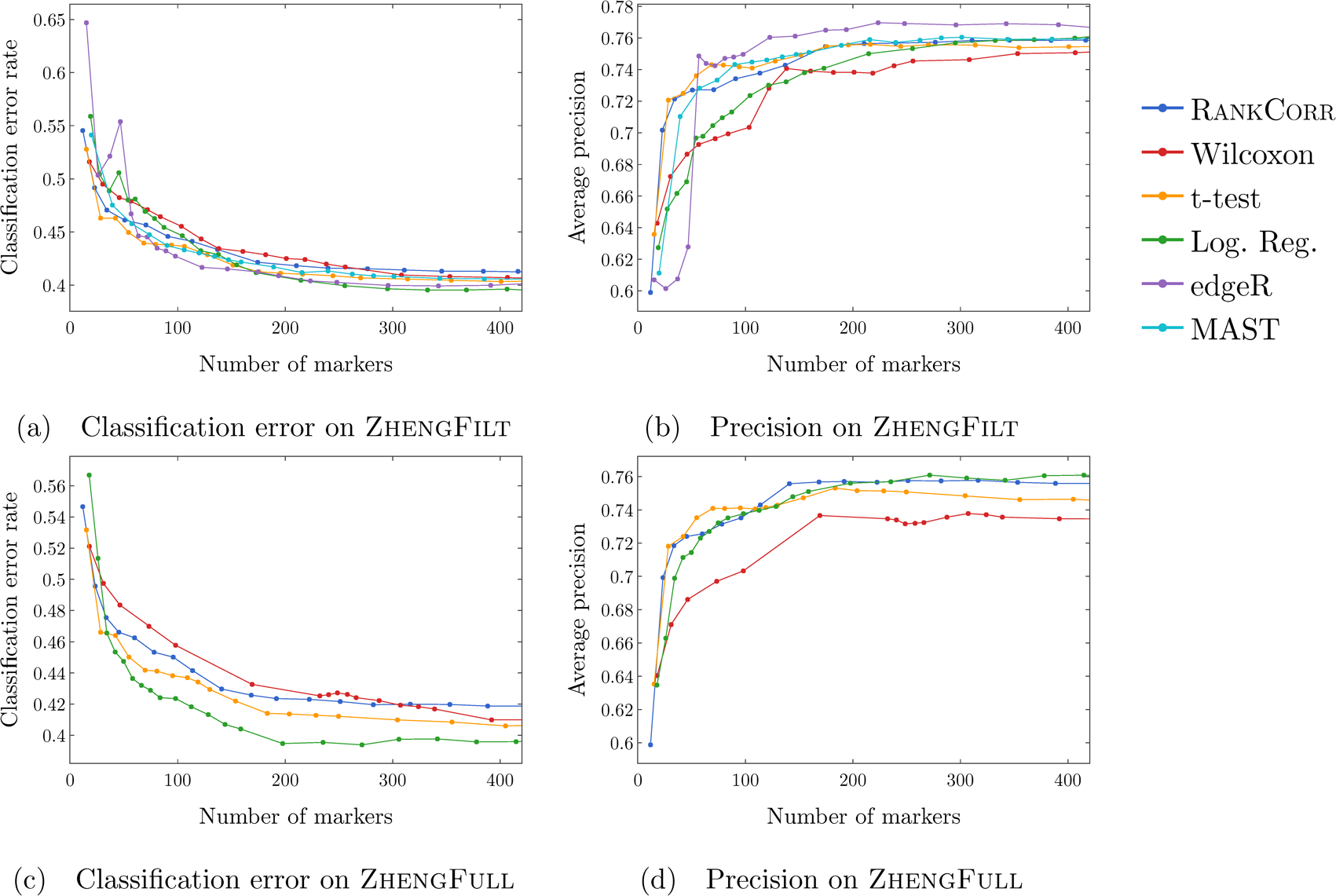
Accuracy and precision of the nearest centroids classifier on the Zheng data sets using the bulk labels. The top row correspond to the ZhengFilt data set and the bottom row corresponds to the ZhengFull data set.

The ARI values are especially low on the Paul data set, and the methods consistently produce lower ARI values than AMI values. This is a change from the Zeisel data set, where the ARI scores were higher than the AMI scores (and the FMSs were the highest of all). The aspects of the data sets that change the relative ordering of the metrics are unclear; it must be the data sets that influence this change, however, since the change persists across the marker selection algorithms.

Another observation is that, in all of these plots, the majority of the methods (apart from scVI) produce similar values of the ARI, AMI, and FM scores when less than 100 markers are selected. As more markers are selected, the methods tend to split up. In all three plots, the Wilcoxon, t-test, MAST, edgeR and RANKCORR methods tend to perform better than the other methods and provide the upper edge of the set of curves. On the other hand, the scVI, logistic regression, elastic nets, and MASTdet curves tend to be the worst performers by these metrics. The edgeRdet curve and the Spa curve generally stay in the middle between the two sets of curves discussed above. Although this stratification is consistent across the three scores, there is almost always less than a 0.05 score difference between the best and the worst scores - that is, all of the marker selection methods exhibit similar performance. Note that the value of *k* was chosen to optimize the RANKCORR method (see the Louvain parameter selection discussion in the Methods) and thus it is not entirely surprising that RANKCORR appears near the top.

The final note is that, on the ARI and FM curves, scVI performs comparably to the other methods when between 200 and 500 total markers are selected - unlike the supervised classification metrics, there is no gap (or only a small gap) between the performance of scVI and the performance of the other methods in this domain. This is a significant change from the classification metrics, where scVI was always much worse than all of the other methods.

There are also several observations that are unique to the different metrics in Figure 5. For the ARI scores (Figure 5(a)), although the methods (other than scVI) show similar performance when less than 100 total markers are selected, the two general groups discussed above also persist for small numbers of markers. That is, the Wilcoxon, MAST, and t-test methods perform slightly better than the other methods in this regime. The maximum difference between any of the scores (other than logistic regression) is around 0.03; i.e. the curves are more tightly grouped in this domain. Moreover, when more than 300 total markers are selected, the Wilcoxon and t-test methods appear to pull slightly ahead of all of the other methods according to the ARI values.

On the AMI plot (Figure 5(b)), the Wilcoxon method outperforms all of the other methods when less than 100 markers are selected, and produces a high AMI score even when selecting two markers per cluster. As more markers are selected, the grouping discussed above is less clear, with the methods separating from each other. The general order of performance is maintained, however.

Finally, on the FM score plot (Figure 5(c)), there is more of a separation between the methods when less than 100 markers are selected. The RANKCORR and t-test methods perform the best in this range, and both methods actually exhibit local maxes when around 60 total markers are selected. The edgeRdet also exhibits this peak, but does not start out as strongly as RANKCORR and the t-test. As more markers are selected, scVI comes close to the lower performing band of methods, but it peels away and moves towards the performance of random marker selection as we select more than 600 markers (see Additional File 1, Figure 6(d))).

The unsupervised clustering metrics provide a more consistent stratification of the marker selection methods that are tested here. The RANKCORR, Wilcoxon, t-test, MAST, and edgeR methods are often the top performing methods when selecting many different total numbers of markers. RANKCORR, the t-test, and Wilcoxon deserve recognition for their higher performance when selecting small numbers of markers according to the AMI and FM scores. That said, there is not a massive separation between any of the methods (apart from scVI), as with the classification metrics.

#### Discussion

As mentioned in the discussion of the supervised classification metrics, the most appropriate conclusion that can be drawn from these data is that most of the marker selection methods show similar levels of performance on the Paul data set. One method, scVI, stands out with worse performance than the other methods. All of the scores produced by all of the methods on the Paul data set are significantly worse than the metrics on the Zeisel data set; however, the methods perform considerably better than random marker selection. The notion of distinct cell types does not fit well with a cell differentiation trajectory; the poor score levels reflect the necessity to come up with a better mathematical description of a trajectory for the purposes of marker selection.

Although elastic nets and edgeR show poor performance for low numbers of markers selected according to the classification metrics, this is not observed in the clustering metrics. Moreover, edgeR is one of the better methods according to the clustering metrics. In a similar fashion, Wilcoxon is one of the better methods for selecting a large number of markers according to the clustering metrics, but it tends to be one of the worst methods for selecting a large number of markers according to the classification metrics.

These disparities emphasize the notion that the evaluation metrics considered in this work do not capture the full “usefulness” of a set of markers and simply present different ways of looking at the data. Designing a metric for benchmarking marker selection algorithms is itself a difficult task, and the optimal metric to consider could depend on the data set in question (as we see with the change in the relative orders of the AMI, ARI and FMS plots between the Paul and Zeisel data sets).

The RANKCORR algorithm is one of only three methods that always performs nearly optimally under every metric examined here; the others are the t-test and MAST. In addition, RANKCORR always performs well when selecting small numbers of markers, and shows exceptional performance in this regime under the Fowlkes-Mallows clustering metric. Combined with the facts that RANKCORR is fast to run and requires low computational resources, this shows that RANKCORR is a worthwhile marker selection method to add to computational toolboxes to use alongside other fast methods.

### Results on the ZhengFull and ZhengFilt data sets

The data set in [25] that we examine contains data from over 68 thousand PBMCs; this is more than 30 times the number of cells in either the Paul or Zeisel data sets. These data are more representative of the sizes of the data sets that we are interested in working with. We mostly focus on ZhengFilt, a version of the data set that includes only the information from the top 5000 most variable genes. We also consider the performance of the fastest algorithms (RANKCORR, logistic regression, Wilcoxon, and the t-test) on ZhengFull, the data set containing all of the genes, to check for any differences. The ground truth clustering that we consider is a labeling obtained by correlation with bulk profiles (biologically motivated “bulk labels”). There are 11 cell types in this clustering. See the section describing the data sets (in the **??**) for more information.

The Zheng data sets consist of PBMCs; therefore, there are some distinct clusters (e.g. B cells), as well as some clusters that are highly overlapping (e.g. different types of T cells). There are not any specific cell differentiation trajectories (that we are aware of), but the overlapping clusters provide a similar challenge for the marker selection methods. Thus, we expect to see performance between that of Paul and Zeisel.

The timings of the methods on ZhengFilt are in Table 6. The gap between the slow methods and the fast methods has increased considerably, and it would not be computationally feasible to run MAST and edgeR on data sets that are much larger. Certainly, running MAST and edgeR would not be possible on a personal machine. The timings of the methods on the ZhengFull data set are shown in Table 7. Even though the data set contains over 27 thousand genes, it is still possible to run the fast methods on a personal computer. Here we begin to see that logistic regression scales worse than the other methods, and it is already becoming slow and computationally heavy on “only” 68 thousand cells.

#### Supervised classification

As in previous data sets the Matthews correlation coefficient curves provide very similar information to the classification error curves and thus they can be found in Additional File 1, Figures 7-10. Unlike the previous data sets, the precision curves are slightly different in some occasions, and thus they are presented here. We start by reporting the classification metrics using the NCC.

**Figure 7:**
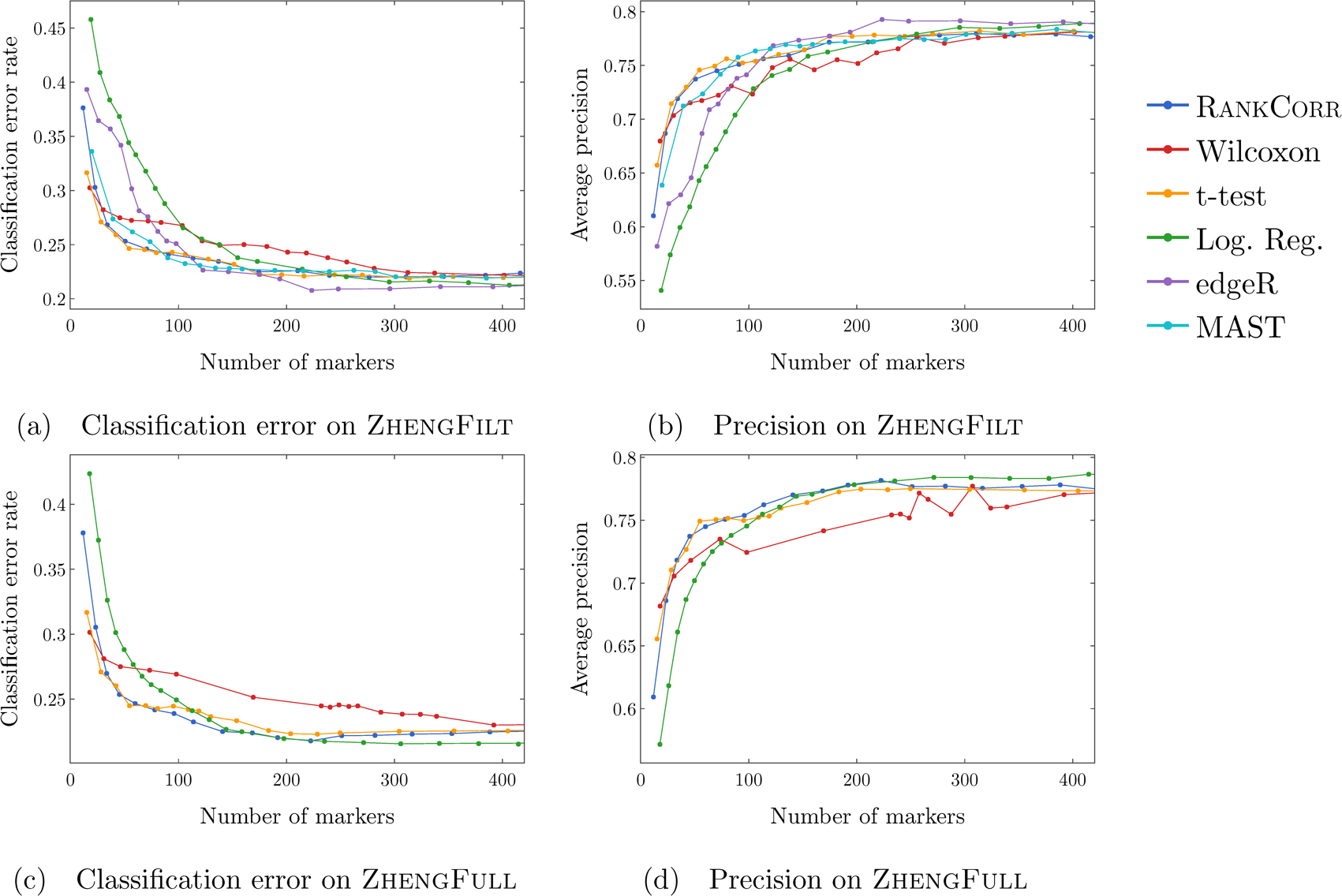
Accuracy and precision of the random forests classifier on the Zheng data sets using the bulk labels. The top row correspond to the ZhengFilt data set and the bottom row corresponds to the ZhengFull data set.

Figure 6 focuses on the performance of the methods when the NCC is used for classification. Overall, the classification error rates for these data are quite high, and don’t level off (to a minimum value of approximately 40%) until around 200 markers are selected; this corresponds to an average of around 18 unique markers per cluster. For very small numbers of markers selected, the classification error rates are quite high (around 55%), and these error rates drop relatively slowly until approximately 200 markers are selected (note that this is still significantly better than random classification on 11 classes). The methods generally perform at approximately the same level on the ZhengFilt data set, though edgeR and logistic regression both show a spike in error rate with about 50 markers are selected. On ZhengFull, the logistic regression method shows the lowest error rates when large numbers of markers are selected, clearly outperforming the other methods.

The precision of these methods is significantly higher than the accuracy. Although the values of precision start low for small numbers of markers selected, they increase quite rapidly until they level off when about 100-150 total markers are selected. In this case, the Wilcoxon method consistently shows the lowest (worst) precision. RANKCORR and logistic regression generally compete for the best performance on ZhengFull, and edgeR shows the highest precision on ZhengFilt when more than 50 markers are selected. Strangely, edgeR starts with very low precision and decreases in precision for a bit before spiking again when around 50 markers are selected. This fits with the error rate curve.

Neither the classification accuracy nor the precision changes by very much when we filter from the full gene set (Figures 6(c) and 6(d)) to the 5000 most variable genes (Figures 6(a) and 6(b)). In general, this filtering very slightly increases both the accuracy and precision of the t-test, Wilcoxon, and RANKCORR methods, while it worsens the performance of the Logistic Regression method. This suggests that enough marker genes are kept by this variable gene filtering process to maintain accurate marker selection.

We examine the performance of the classification metrics using the RFC in Figure 7. The error rates using the RFC are decreased significantly compared to the NCC, and level off to approximately 22% when large numbers of markers are selected. The error again does not completely level off until around 200 total markers are selected, but there is a steeper initial descent: the RANKCORR and t-test methods reach an error rate of slightly less than 25% when 55 total markers are selected (an average of 5 markers per cluster) on both the ZhengFull and ZhengFilt data sets, and MAST performs similarly on ZhengFilt. Overall performance is similar between ZhengFull and ZhengFilt, again showing that enough markers are maintained by the variable gene filtering.

On ZhengFilt, the RANKCORR, t-test, and MAST methods outperform the other methods in terms of error rate when less than 120 total markers are selected. Wilcoxon starts off with a very low error rate when around 2 markers are selected per cluster, but it does not improve with the other methods, and it shows the highest error rates and lowest precisions when more than 150 total markers are selected (an average of 14 unique markers per cluster). edgeR starts with a relatively high error rate and low precision, and improves more slowly than the top algorithms, catching up to the best methods when around 120 total markers are selected. Logistic regression is the worst performer when small numbers of markers are selected. It shows an error rate that is 12% to 15% higher than the RANKCORR, MAST, Wilcoxon, and the t-test when selecting 25-30 unique markers (an average of 2-3 markers per cluster).

When more than 150 total markers are selected, Wilcoxon stands out as the method with the poorest performance. The other methods, however, perform very similarly in this range. EdgeR shows the lowest error rates and highest precisions when more than 200 total markers are selected, but the improvement is only 1% to 2% at most.

On ZhengFull, similar patterns are visible, and the advantage shown by logistic regression on ZhengFull under the NCC has disappeared. In particular, RANKCORR and the t-test consistently show the lowest error rates and highest precisions. Logistic regression starts poorly but catches up to RANKCORR when around 120 total markers are selected. On the other hand, Wilcoxon shows poor performance when more than 100 total markers are selected; this trend is more pronounced than on ZhengFilt.

These RFC results are close to our expectations. The steep initial descent in error rates could appear as the large groups of cells are separated from each other (e.g. B cells from T cells) and the slower improvement from 100 to 200 of total markers selected could be the methods fine-tuning the more difficult clusters (e.g. Regulatory T from Helper T). The error rates are between those observed in Paul and Ziesel. On the other hand, the error rates for the NCC classifier are much higher than expected. We thus consider the results using the RFC as more informative than the results using the NCC on the ZhengFilt data set.

#### Unsupervised clustering

We focus on the ZhengFilt data set for the clustering metrics. This is due to the fact that the classification metrics are changed only slightly between ZhengFilt and ZhengFull as well as the fact that Louvain clustering on the large Zheng data set is itself time and resource intensive.

The clustering metrics on the ZhengFilt data set are presented in Figure 8. All three scores are generally quite low, though they are again mostly much higher than random marker selection. The performance of random marker selection can be found in Additional File 1, Figure 11. Several patterns are present in all three plots.

**Figure 8:**
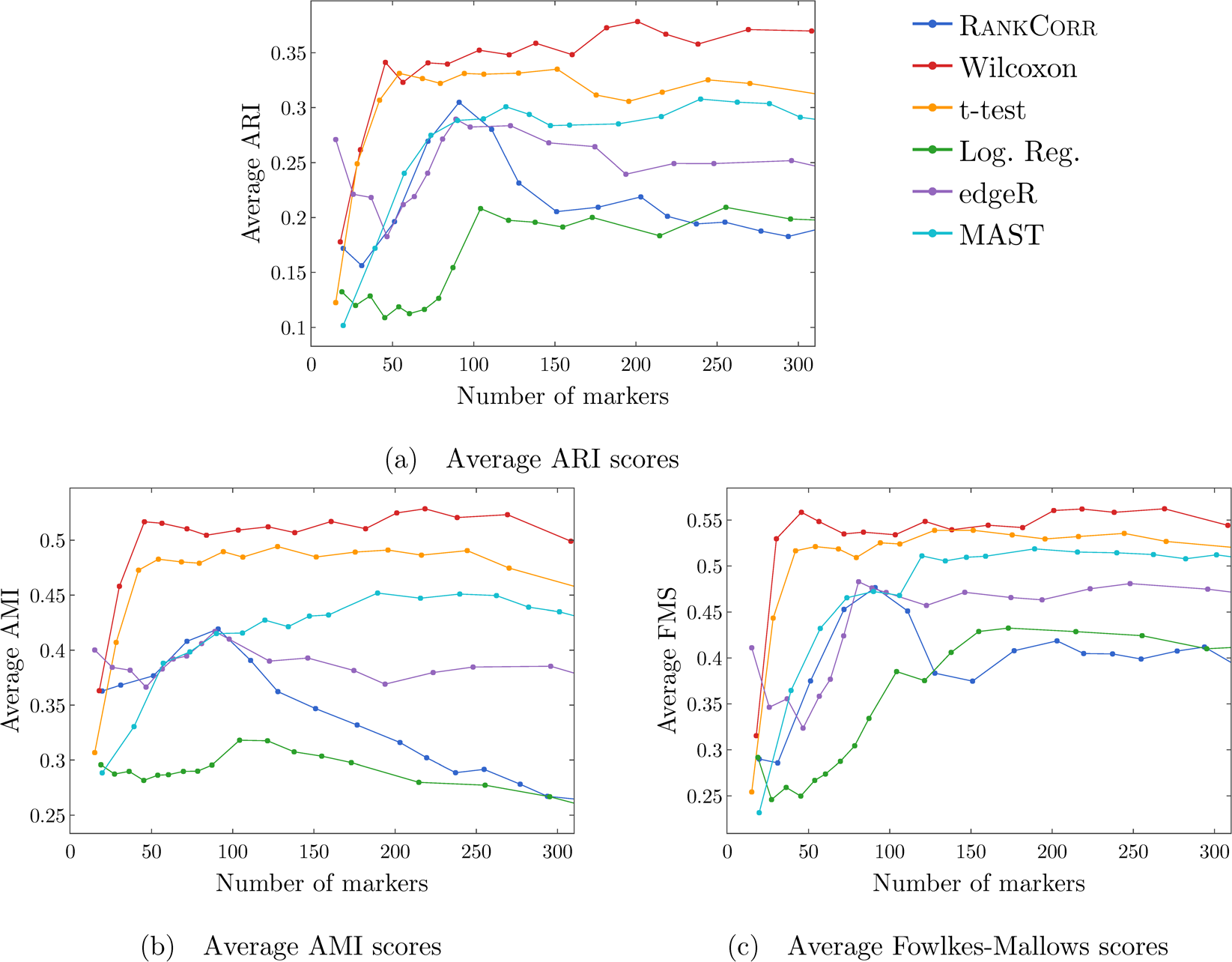
Clustering metrics on the ZhengFilt data set.

**Figure 9:**
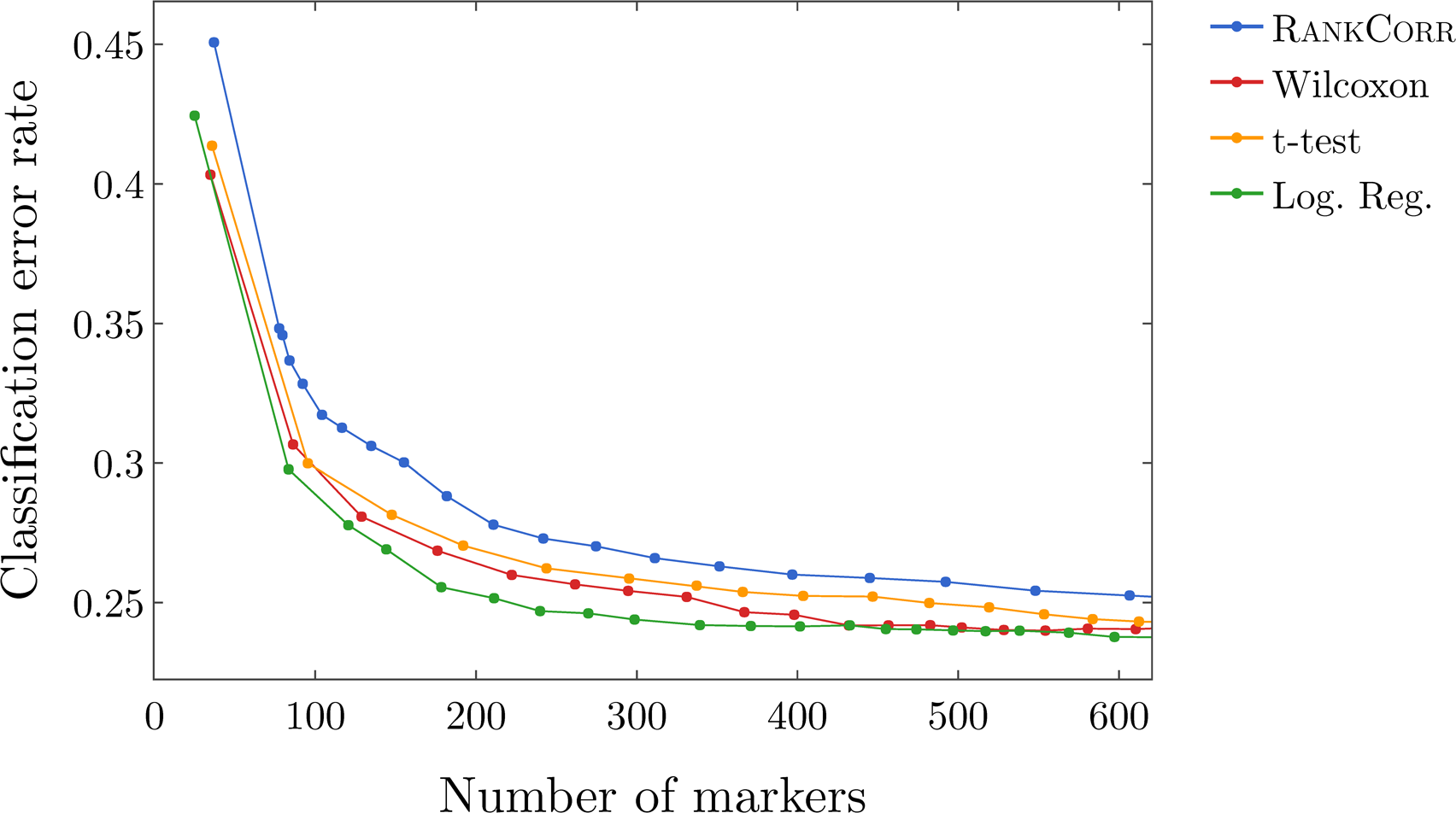
Classification error rate under the NCC vs number of markers on the full 10xMouse data set.

**Figure 10:**
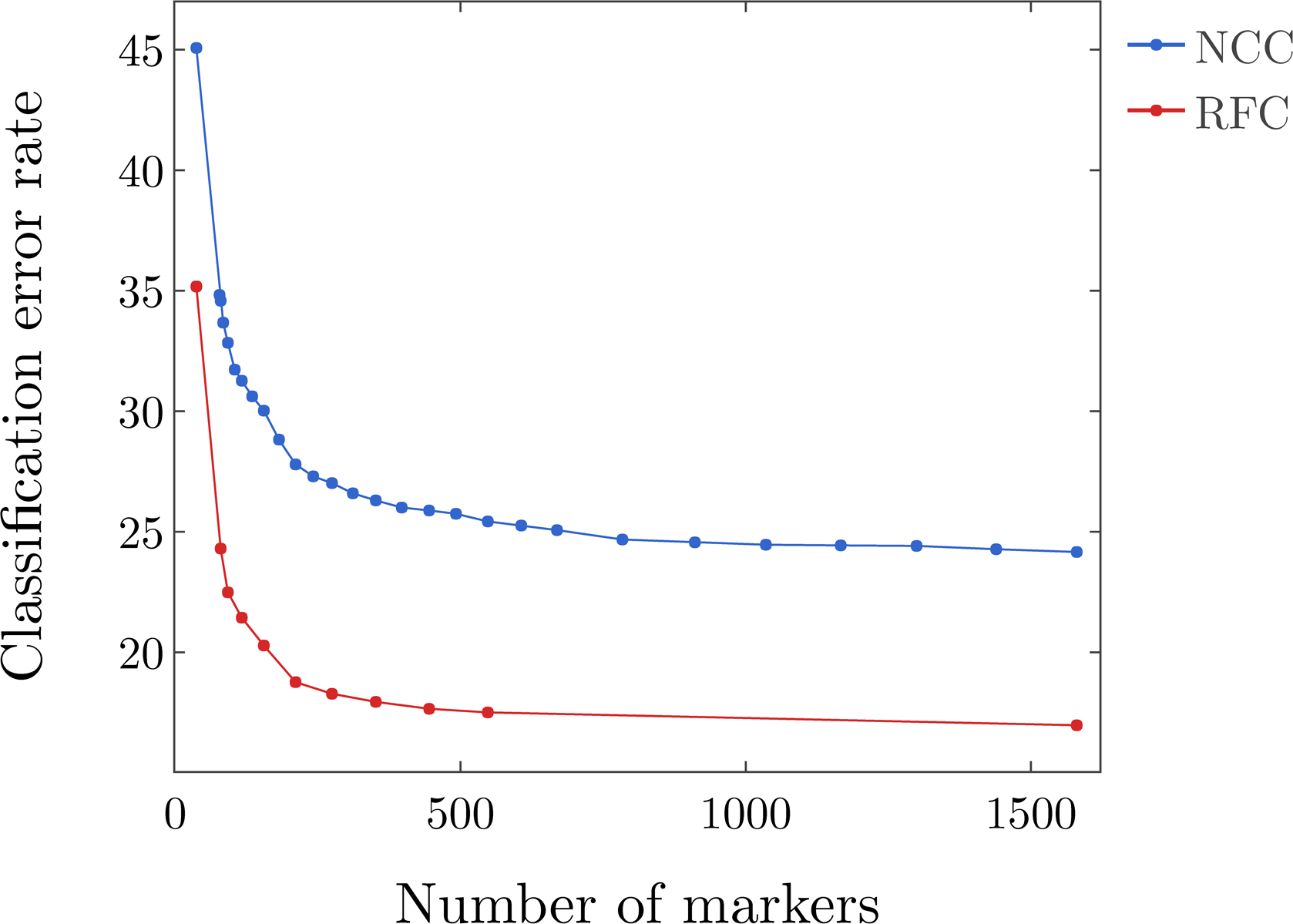
A comparison of the nearest centroid classifier (NCC) and the random forest classifier (RFC) using the RANKCORR method on the 10xMouse data set

**Figure 11:**
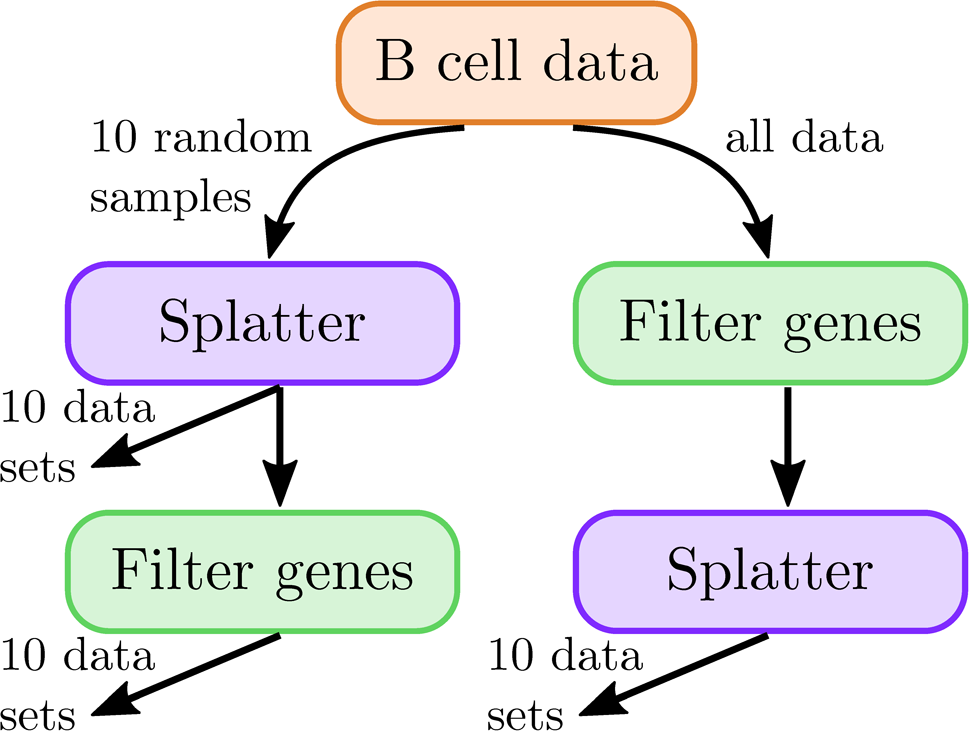
Set up of the simulated data. We consider 3 conditions. On the left side of this diagramme, we produce 10 data sets from using all genes in simulation, and 10 more from filtering after simulation (these data sets containing a subset of the information from the “all genes used for simulation” data sets). On the right hand side, we produce 10 data sets by filtering genes before simulation.

First, all of the methods start very low scores when small numbers of markers are selected; the ARI values for most methods are between 0.1 and 0.17 when around 20 total markers are selected. The Wilcoxon and t-test methods produce scores that increase rapidly until 50 total markers are selected, then their scores level off. These two methods consistently produce the highest scores, with Wilcoxon the best overall. On the other hand, the scores for logistic regression don’t improve significantly until around 75 total markers are selected and then remain low (either worst or second worst) for high numbers of selected markers. The edgeR, MAST, and RANKCORR methods perform similarly to each other when between 50 and 100 total markers are selected, and produce values that are above logistic regression but below Wilcoxon in this range. The three methods then split up, with MAST doing better than edgeR which does better than RANKCORR. The scores for the RANKCORR method decrease after about 100 total markers are selected until the RANKCORR curve meets up with the logistic regression curve. EdgeR always starts with the highest scores for small numbers of total markers selected, but its scores decrease until about 50 total markers are selected, where it meets up with the MAST and RANKCORR curves.

#### Discussion

Apart from the t-test, the methods show inconsistent performance when comparing the clustering metrics to the classification methods. It is possible that changing the number of nearest neighbours considered in the Louvain clustering would produce more consistent data. Although the clustering metrics did not appear to change significantly when altering the number of nearest neighbours on the previous data sets (see the Louvain parameter selection information in the Methods), the ZhengFilt data set is much larger than those previous data sets; the larger number of cells may necessitate the use of information from more nearest neighbors to recreate the full clustering structure.

It is also possible that the bulk labels that are used for the ground truth are difficult to reproduce through the Louvain algorithm. We generated a clustering that visually looked like the bulk labels via the Louvain algorithm (see Figure 19); the ARI, AMI, and FMS values for the generated louvain clustering compared to the bulk labels are in the ranges produced by the Wilcoxon and t-test methods (not larger than the scores here). In addition, the top ARI and AMI scores (produced by the Wilcoxon and the t-test methods) are comparable to (or only slightly better than) the scores on the Paul data set (Figure 5). This runs counter to our expectations: the Paul data set contains a cell differentiation trajectory, with no real clusters that are easy to separate out, while the Zheng data sets contain several clusters that are well separated. It is possible that the bulk labels produce clusters that are more mixed than it appears in a UMAP plot.

In any case, the disparity between the different types of scores emphasizes the fact that the classification and clustering metrics provide different ways of looking at the information contained in a selected set of markers. Methods that perform well according to both types of metrics should be preferred.

That said, the t-test produces the overall best results on the ZhengFilt data set. It performs well under the classification metrics, especially for small numbers of total markers selected. In addition, it is one of the best methods under the clustering metrics, where it is competitive with the Wilcoxon method.

Other methods of note are logistic regression, edgeR, and Wilcoxon. Logistic regression appears to be the best method on the ZhengFull data set using the NCC; however, according to the RFC, logistic regression performs quite poorly on ZhengFull when a small number of markers are selected. Since the error rates are overall quite similar between the ZhengFull and ZhengFilt data sets, but the running times of the methods are significantly faster on ZhengFilt, it seems worthwhile to select the most variable genes before marker selection, and logistic regression is never optimal on ZhengFilt according to the classification metrics. Combining this with the poor performance in the clustering metrics, logistic regression is not a recommended method for the Zheng data sets.

The edgeR method exhibits the top performance on the ZhengFilt data set after more than 50-100 unique markers are selected according to the classification metrics. The classification metrics show edgeR as one of the worst methods when choosing less than 50 unique markers, however. This is in direct contradiction to the clustering metrics, where edgeR is always the best method for the smallest (*∼* 20) total numbers of markers selected, and it then shows performance in the middle of the other methods as larger numbers of markers are selected. Since many researchers are interested in selecting smaller numbers of markers, edgeR’s advantages according to the classification metrics are somewhat irrelevant. Thus, edgeR can not be particularly recommended on ZhengFilt either.

Likewise, Wilcoxon consistently shows the worst performance according to the classification metrics as larger numbers of markers are selected (though it does quite well when selecting around 2 markers per cluster, or around 20 unique markers). On the other hand, it is the best method according to all of the clustering metrics except when selecting a very small number of markers. It is thus also difficult to recommend Wilcoxon, despite it’s dominant performance according to the clustering metrics.

The other two methods, RANKCORR and MAST, show good performance on the ZhengFilt data set under the classification metrics, and are generally competitive with the t-test, especially when selecting smaller numbers of markers. They both show middling performance under the clustering metrics, however. Unlike logistic regression, wilcoxon, and edgeR, there are no areas where they perform especially poorly. They are still outperformed by the t-test according to the clustering metrics, however.

### Marker selection on the 1 million cell 10xMouse data set

Here, we consider the 10xMouse data set which consists of 1.3 million mouse neurons generated using 10x protocols [26]. The “ground truth” clustering that we examine in this case was algorithmically generated without any biological verification or interpretation (see the section about data sets in the Methods). We include this data set as a stress test for the methods and therefore we do not perform any variable gene selection before running the marker selection algorithms (to keep the data set as large as possible). We also only consider the four fastest and lightest methods (RANKCORR, the t-test, Wilcoxon, and logistic regression) as these are the only methods considered in this work that could possibly produce results in a reasonable amount of time on this data set.

Table 8 contains the timing information for the methods on the data set. Logistic regression is by far the slowest method, and it is difficult to call the time and resources required by logistic regression “reasonable.” It is not clear that logistic regression could be used on any larger data sets (when they appear), while the other three methods could be used quite easily. RANKCORR is the second slowest, but it quite easy to parallelize, and the total required CPU time is only around four times that required by Wilcoxon.

**Table 8:** Timing and computation resources required on the 10xMouse data set. Times are reported to compute the markers for one fold and are approximate. Logistic regression did not converge for some clusters.

| Method | Timing | Resources |
| --- | --- | --- |
| t-test | 2 hours | 1 CPU, 80 GB RAM |
| Wilcoxon | 5 hours 30 minutes | 1 CPU, 80 GB RAM |
| Log. Reg. | 10 hours 30 minutes | 8 CPU, 80 GB RAM |
| RANKCORR | 2 hours | 10 CPUs, 80 GB RAM |

Figure 9 shows the classification error of the four methods on the 10xMouse data set using the NCC. There are 39 clusters in the “ground truth” clustering that we examine here - thus, the error rates produced by all of the methods are much lower than the error rate expected from random classification. In Figure 9, we see that the logistic regression method performs the best overall, and that RANKCORR consistently shows the highest error rate. The largest difference between the RANKCORR curve and the logistic regression curve is only around 3%, however. In addition, as mentioned above, logistic regression is the slowest method by far on this data set - extra accuracy is not worth much if the method is not able to finish running.

Due the fact that a biologically motivated/interpreted clustering may be quite different from the clustering used here, and considering that the classification error rate is not necessarily an accurate indication of the relative performance of methods that show similar error rates (see the following section about synthetic data), it is only possible to conclude that all four methods examined here show similar performance on the 10xMouse dataset. The RANKCORR method produces interpretable markers, runs in a competitive amount of time, and takes a step towards selecting a smart set of markers for each cluster (rather than the same number of markers per cluster).

As with previous data sets, the precision and Matthews correlation coefficient curves do not provide extra information; they can be found in Additional File 1, Figure 12. Moreover, the implementation of the RFC in `scikit-learn` was quite slow on the large 10xMouse data set, and thus we do not compare the methods via the RFC. From the smaller data sets, we might expect that the random forest classifier produces curves that are shaped similarly to the ones in Figure 9 but are shifted down to a lower error rate. This is indeed what we see for the RANKCORR method: a comparison of the RFC and the NCC is shown in Figure 10. Each point on the RFC curve in Figure 10 took over 3 hours on 10 CPU cores to generate; the largest point took over 15 hours. The Louvain clustering method was also too slow to compute any clustering error rates for the markers selected here. This is a situation where the marker selection algorithms are faster than almost all of the evaluation metrics (emphasizing the continued need for good marker set evaluation metrics along).

**Figure 12:**
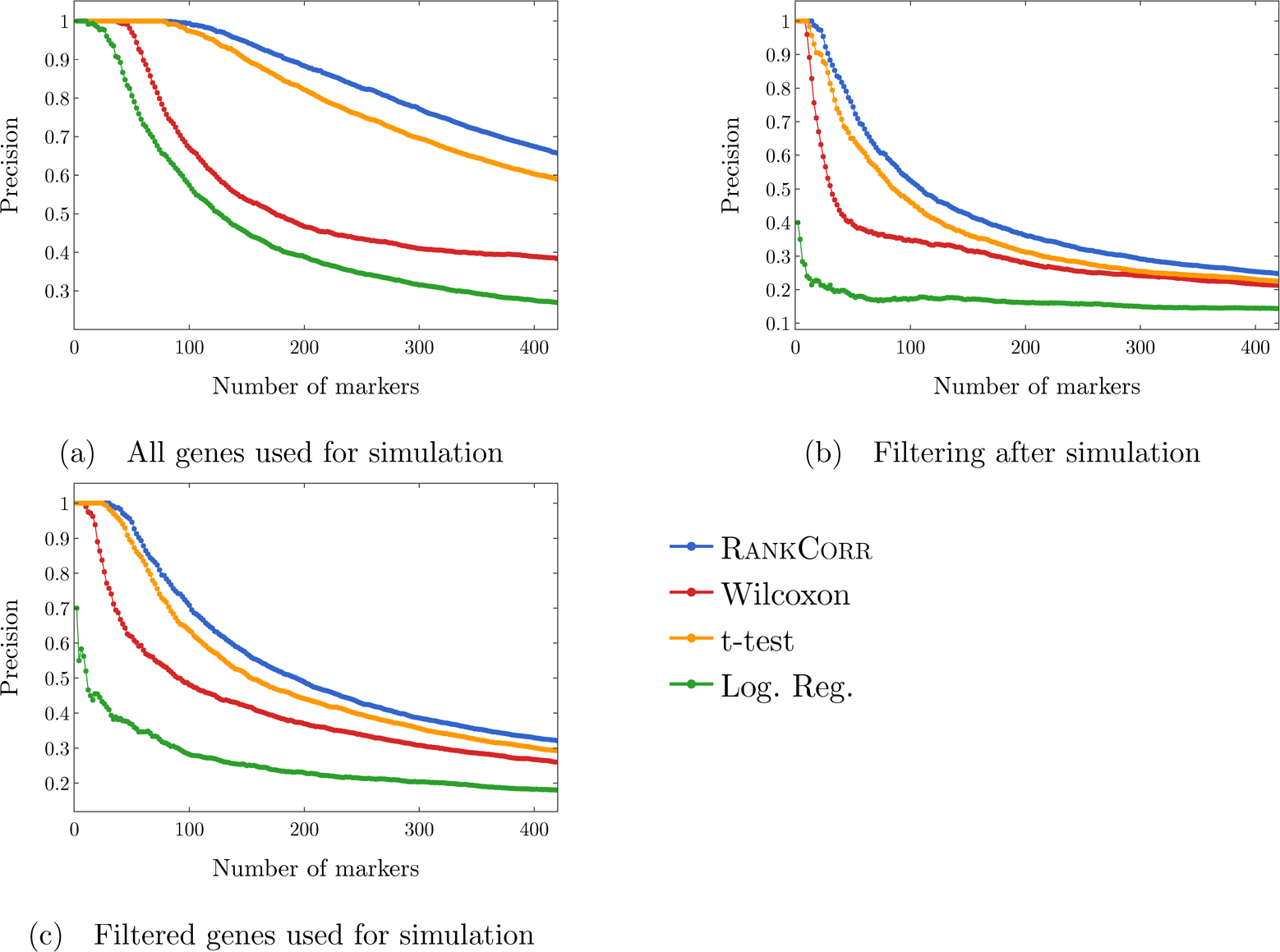
Precision of the marker selection methods versus the number of markers selected for the first 500 markers selected. Each sub-figure corresponds to a simulation method and the four lines correspond to the different marker selection algorithms. The RANKCORR method consistently shows the highest precision across all three simulation methods.

### Comparison of marker selection methods on synthetic data

We have evaluated several of the marker selection algorithms on some synthetic data sets that are designed to look like experimental scRNA-seq data. The data are generated using the Splatter R package [27]; see the synthetic data generation section of the Methods for specifics. In short, we use cells from the data set consisting of purified (CD19+) B cells that is introduced in [25] to estimate the simulation parameters. We generate 20 different simulated data sets from the CD19+ B cells. Each data set consists of 5000 cells that are split into two groups; 10% of the genes are differentially expressed. We also examine the effects of filtering down to the 5000 most variable genes: in 10 samples, we filter before simulating (and simulate 5000 genes); in the other 10 samples, we simulate without filtering (and simulate as many nonzero genes as there were in the input data, usually around 20000 genes). From each data set that was simulated without filtering, we produce another data set by filtering down to the 5000 most variable genes. This results in a total of 30 data sets; Figure 11 summarizes this design.

Since the differentially expressed genes are chosen at random by Splatter, many of the genes that are labeled as “differentially expressed” in the output data have low expression levels (often they are expressed in less than 10 cells). Thus, here we report the precision, true and false positive rates (TPR and FPR), and classification error rates for the marker selection algorithms. We do not consider the recall since we do not (in general) want marker selection methods to be selecting genes with very low expression levels - and, in fact, these genes are usually not selected as markers by any of the methods that we test here. The values of precision, TPR, and FPR are computed without cross-validation, since each entire set (of genes) in each data set is test data - there is no training to be done. The classification error rate is still computed with 5-fold cross-validation.

Since the speed of a marker selection algorithm has been observed as an important factor for use on actual data, we focus here only on the fastest methods: RANKCORR, the t-test, Wilcoxon, and logistic regression.

### Simulated data illuminates the precise performance characteristics of marker selection methods

In Figure 12, we examine the precision of the marker selection algorithms for the first 400 unique genes selected. Apart from the t-test curves, each curve represents the average across all 10 simulated data sets that are relevant to the curve. There were issues with selecting stable sets of genes using the t-test in one of the trials in each simulation condition (due to tied t-test scores); thus, the t-test precision, TPR, and FPR curves each represent the average of the 9 data sets that are relevant to the curve.

Precision is an important metric for marker selection - it is desirable for the algorithms to select genes that actually separate the two data sets (rather than genes that are statistically identical across the two populations). Thus, it is promising to see that RANKCORR produces the highest precision in marker selection across all of the simulation methods. The t-test is consistently second, the Wilcoxon method is consistently third, and logistic regression consistently exhibits the lowest precision.

In Figure 12, we see that the methods generally start off with high precision that decreases as more markers are selected (each data set contains more than 400 differentially expressed genes). In both of the filtered simulation conditions, all of the methods get close to a precision of 0.1 (indicating random gene selection) when 400 markers are selected, and all of the curves are still decreasing at this point. There are around 2000 differentially expressed genes in the un-filtered simulation condition, so the fact that the precision drops significantly when selecting up to 400 markers indicates proportionally similar behavior to the filtered data sets.

The ROC curves in Figure 13 also show similar behavior: the curves increase (above the main diagonal) quite rapidly for a short period of time, but then remain close to the diagonal overall. These ROC curves are somewhat expected: in the simulations that come from all of the genes, many of the “differentially expressed” genes show low levels of expression. Thus, we would expect that the methods end up close to the diagonal as intermediate to large numbers of total markers are selected (since finding these low expression markers should be close to random selection). The filtered data sets could have solved this problem; however, the filtering method used here preserves the relative proportions of low- and high-expression genes and (possibly for this reason) do not affect the ROC curves very much.

**Figure 13:**
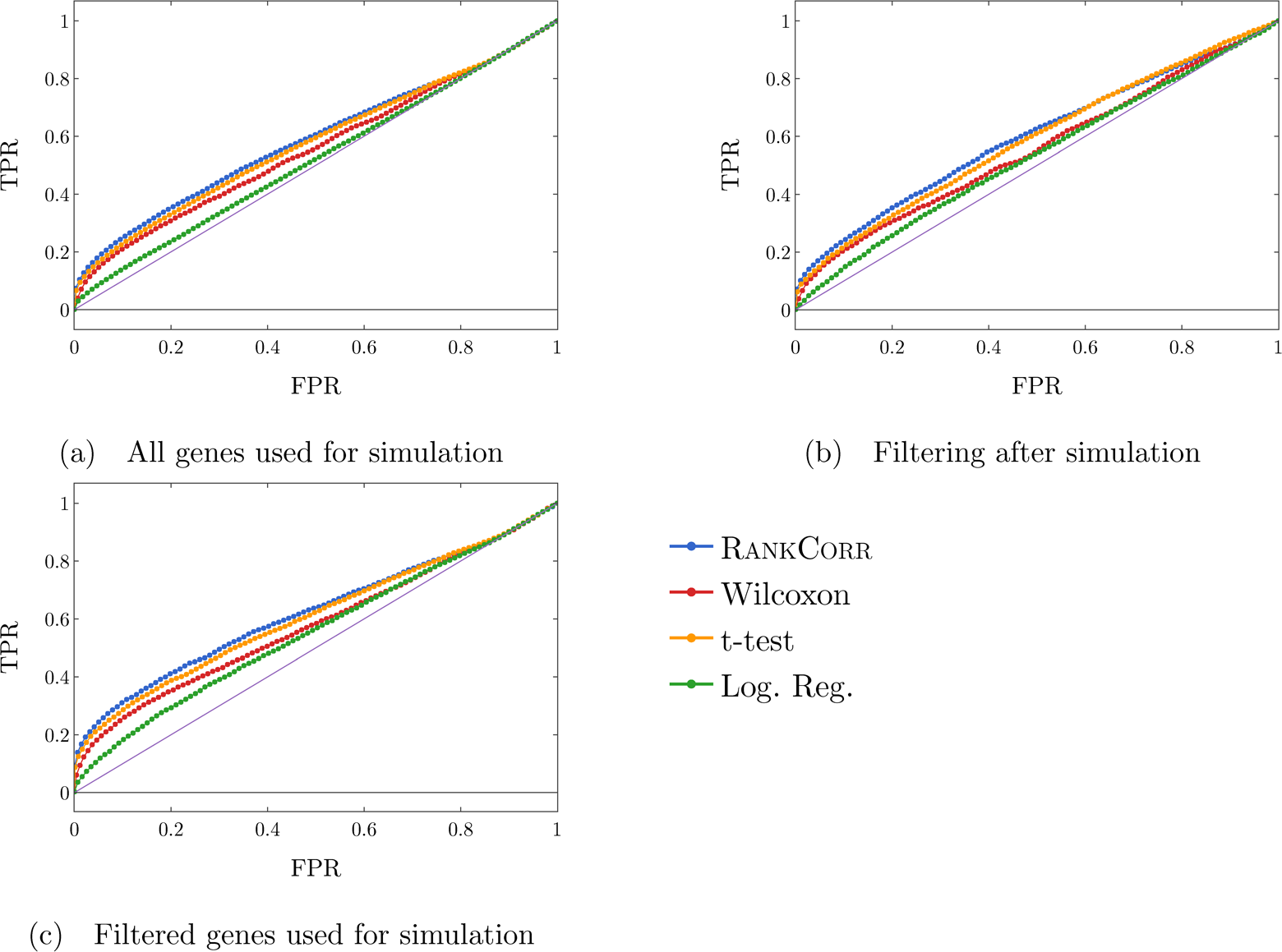
ROC curves. Each sub-figure corresponds to a simulation method and the four lines correspond to the different marker selection algorithms. The solid (purple) line is the diagonal TPR = FPR.

The ROC curves never get very far above the diagonal, however, and the precision curves (for the filtered conditions) are already quite low when only 400 markers are selected. This cannot only be explained by low expression markers. One explanation for these extra difficulties could be the differential expression parameters used in the Splat simulation. With these default parameters, the gene mean for some of the “differentially expressed” genes are only slightly different between the two clusters. Thus, although the simulation parameters may show that these genes are differentially expressed, detecting the differential expression by any method will be very difficult. In any case, these genes would not be good markers for real world examples, as it would be very difficult to tell two clusters apart based on the expression levels of these types of genes without collecting a lot of data.

In any case, it appears that the methods are able to easily identify a small set of differentially expressed genes but then rapidly start to have difficulties as more genes are selected. It would probably be worthwhile to come up with a more precise definition of a marker gene (versus a differentially expressed gene) in order to eliminate the types of problems discussed here. At the very least, it seems that a marker gene should be associated in some way with a measure of how useful of a marker it is (again, a differentially expressed gene that shows low expression is not a particularly useful marker). These types of definitions are saved for future work. The important point for this work is that the RANKCORR method consistently shows the highest value of TPR

### Potential issues with simulated data are revealed through marker algorithm performance

From comparing Figures 12(a-c) across the simulation conditions, we see that the highest precision for each of the marker selection methods is obtained by using all genes for simulation, without any filtering. This is somewhat reasonable, since there are approximately four to five times more differentially expressed genes in the data sets generated from the unfiltered original data than in the simulations generated from filtered data - even though many of the differentially expressed genes show low levels of expression, we still are drawing from a much larger pool.

It is tougher to explain why filtering after simulating produces lower precision scores than not filtering at all. Since the t-test (for example) works by choosing genes based on a p-value score, and the genetic information is not changed by the filtering process (p-values would be the same in both the unfiltered and filtered after simulation data sets), it must be the case that many of the differentially expressed genes are removed from the data set when we filter after simulation. The highly variable genes selected by the filtering method used here (see the Methods) are not required to have high expression; thus, there is no obvious reason that many differentially expressed genes should be filtered out.

Simulating scRNA-seq data is itself a difficult task, and the filtering process is quite heuristic, so it is unclear where the issues arise along the pipeline. At the very least, it is clear that this gene filtering process does not commute with the simulation process, since filtering before simulation shows higher precision than filtering after simulation. There is still more work to be done in fully understanding how the filtering process impacts real scRNA-seq and how to accurately simulate scRNA-seq data.

### The classification error rate is an informative but coarse metric

Finally, we examine the classification error rate in Figure 14. It is interesting to note that the classification error rate is quite high: with only two clusters, we are still misclassifying a minimum of around 10% of cells. This suggests that the simulated data are not well separated - the differential expression introduced by the Splat method is not strong enough to easily separate the two clusters. Moreover, apart from the curves corresponding to the Logistic Regression method, all of the curves look to be fairly constant after a small number of markers have been selected (approximately 50 for the simulations based on all genes and approximately 30 for the simulations based on filtered data). This further supports the discussion from the above sections - only a small portion of the differentially expressed genes provide useful information about the given clustering, and these are the genes that are chosen first.

**Figure 14:**
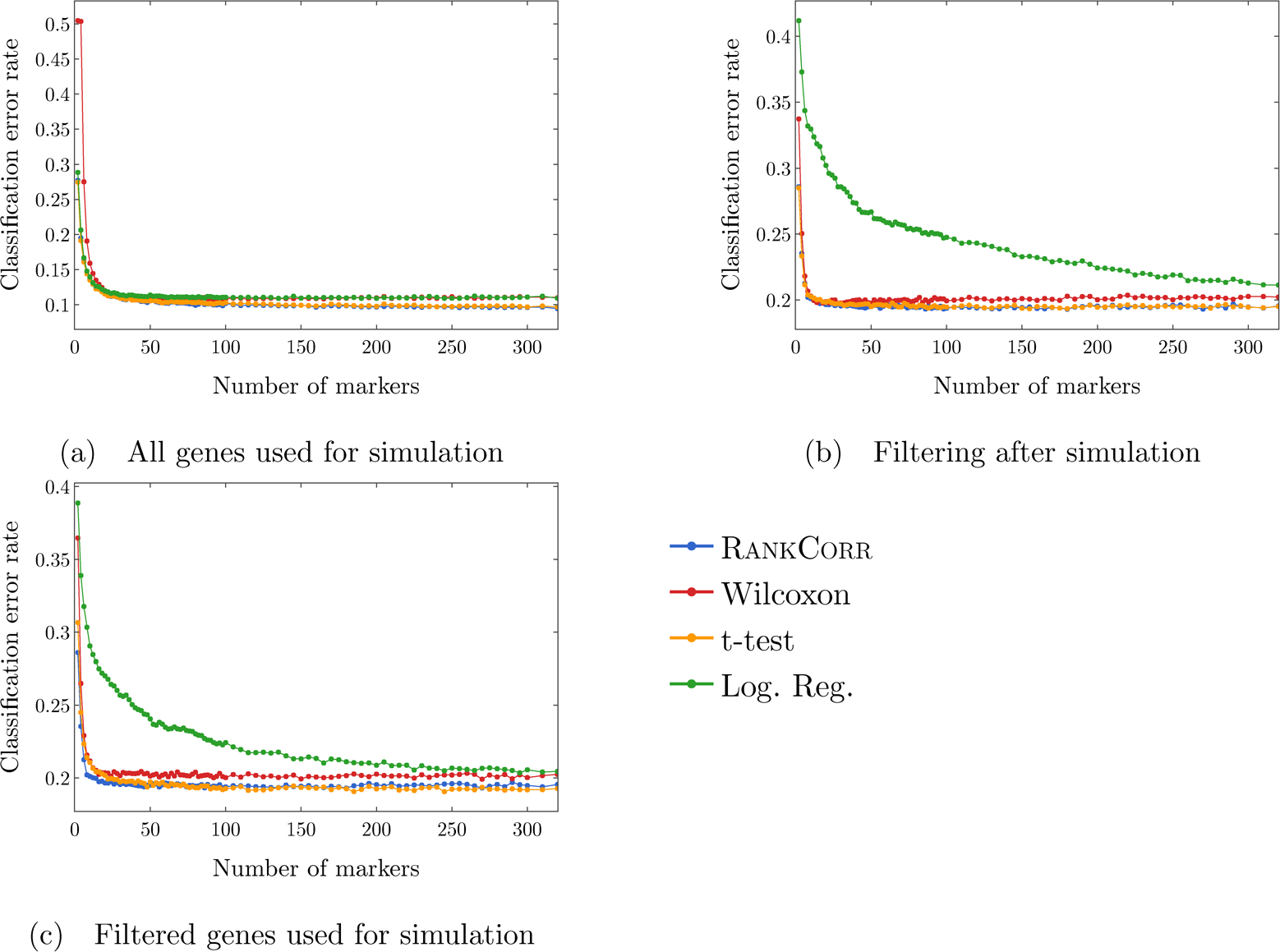
Clustering error rates using the Random Forest classifier for the first 500 markers chosen by each method. The sub-figures correspond to different simulation conditions. The RANKCORR algorithm consistently produces the smallest values of the clustering error rate.

Note that the methods that show higher precision in Figure 12 also generally show a lower classification error in Figure 14. On the other hand, logistic regression shows poor precision levels on the filtered data sets and also appears significantly worse than the other methods in the classification error rate curves. Thus, according to these experiments, the classification error rate seems to be a coarse but reasonable measure of how well a set of markers describes the data set. In this example, if one methods performs worse than another method according to the classification error rate curves (Figure 14), then the same relationship holds in the precision curves (Figure 12). Some large differences in precision are eliminated in the classification error rate curves, however, and thus the classification error rate should always be considered with a grain of salt.

## Conclusions

Across a wide variety of data sets (large and small; datasets containing cell differentiation trajectories; datasets with well separated clusters; biologically defined clusters; algorithmically defined clusters) and looking at many different performance metrics, it is tough to conclusively say that any of the methods tested here selects better markers than any of the others. Indeed, the marker selection method that was “best” depended on the data set that was being examined as well as the evaluation metric in question. Even then, the difference in performance between the best marker selection algorithm and the worst was often quite small.

Thus, the major factors that differentiate the methods examined in this work are the computational resources (both physical and temporal) that the methods require. Since the algorithms show similar overall quality, researchers should prefer marker selection methods that are fast and light.

In addition to this, as technology advances, the trend is towards the generation of larger and larger data sets. High throughput sequencing protocols are becoming more efficient and cheaper, and other statistical and computational methods are improved when many samples are collected. Through imputation and smoothing methods, a detailed description of the transciptome space can be revealed even when low amounts of reads are collected in individual cells. Thus, the speed of a marker selection algorithm will only become more important.

The RANKCORR, Wilcoxon, t-test, and logistic regression methods run the fastest of all of the methods considered in this work. They run considerably faster and/or lighter than any of the complex statistical methods that have been designed specifically for scRNA-seq data. Logistic regression does not scale well with the size of the data set, however, and it requires an amount of resources that is not competitive with the other three methods on the largest data sets. Moreover, logistic regression exhibits poor performance on several of the data sets considered in this work, especially when selecting small numbers of markers. Thus, as a general guideline, RANKCORR, Wilcoxon, and the t-test are the optimal marker selection algorithms to consider for the analysis of large, sparse UMI counts data. This recommendation is further bolstered by the fact that these three algorithms tend to perform well in the experiments that we have considered here, especially when selecting lower numbers of markers.

The RANKCORR algorithm, introduced in this work, is the slowest of the three recommended algorithms. Despite this, it provides added interpretability in the multi-class marker selection scenario. Specifically, RANKCORR attempts to select an informative number of markers for each cluster (rather than just a fixed number for each cluster), generally selecting more markers for clusters that we are less certain about. This would prove useful when investigating the selected markers for individual clusters after selecting a full set of markers for the data. The work of properly selecting sets of markers in a multi-class scenario has not been completed, however, and RANKCORR only proposes one step. Nonetheless, as a fast and efficient marker selection algorithm, RANKCORR is a useful tool to add into computational toolboxes.

RANKCORR also involves taking a rank transform of scRNA-seq counts data. The rank transformation has other uses in scRNA-seq; it is thus useful to understand the further properties of the rank transformation. These properties will be explored in upcoming work.

### The difficulties of benchmarking and the importance of simulated data

Benchmarking marker selection algorithms on scRNA-seq data is inherently a difficult task. The lack of a ground truth set of markers requires for us to devise performance evaluation metrics that will illuminate the information contained in a selected set of genes. We have examined several natural evaluation metrics in this work; these metrics sometimes produce conflicting results, however. Our experiments herein make it clear that these metrics provide different ways to view the information contained in a set of genes rather than capturing the full picture provided by of a set of markers.

Having a ground truth set of markers available makes the evaluation of marker selection algorithms much more explicit. In the analysis on synthetic data here, for example, it becomes apparent that the methods rapidly select a set of markers that provide a lot of information about the clustering, then essentially start picking things by chance. This type of behavior can only be revealed by a study with a known ground truth.

On the other hand, simulating scRNA-seq data is itself a difficult problem. The simulated data that we consider in this work behaves strangely when we filter it by selecting highly variable genes. In particular, the filtering process seems to remove many of the useful differentially expressed genes in the simulated data. This type of behavior was not observed in the experimental data, where working only on high variance genes had little impact on the marker set evaluation metrics. Better simulation methods, and mathematical results formalizing the quality of simulated data, are extremely important future projects.

Finally, in the way that data processing pipelines are currently set up, researchers will often be forced to select markers without the knowledge of a ground truth set of markers. Thus, it may be valuable to consider metrics such as the ones devised in this work when performing marker selection. Combining the values of several of the metrics may help to aid researchers in deciding when they have selected enough markers to adequately describe their cell types (so that they are not considering genes that were chosen at random), for example. The question of how to stop selecting markers is another important consideration for future work.

### The relationship between marker selection and the process of defining cell types

The marker selection framework considered in this work is quite narrow. It is focused on discrete cell types, and (as shown in the Paul data set) does not handle trajectory patterns very well. Moreover, we assume that the genetic information that we supply to a marker selection algorithm consists of cells that are already partitioned into cell types. This is consistent the data processing pipeline that many researchers currently follow (cluster the scRNA-seq data with an algorithm, then find markers for the clusters that are produced); it seems more reasonable to allow for marker selection to help guide the process of finding cell types, however.

For example, future marker selection methods could find markers that are useful for identifying certain regions of the transcriptome space (in an unsupervised or semi-supervised manner). This would allow for clarity along a cell differentiation pathway - at any point on the trajectory, a researcher could see which of the selected markers identify that area, and to what degree. Thus, cell types (or differentiation pathways) could be suggested based on marker genes. These cell types might themselves reveal more informative markers, creating an iterative process: let the markers guide the clustering and vice versa. Such a method is known as an “embedded” feature selection method in the computer science literature.

## Methods

### Details of the RANKCORR algorithm

#### A fast algorithm for solving the optimization (4)

Algorithm Select finds the support of the solution to the optimization (4) given a measurement matrix *A* ∈ ℝ^*n*×*p*^, a vector *τ* ∈ ℝ*^n^*, and a sparsity parameter *s*. It runs as follows:

1. Let 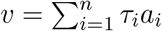 where *a*_*i*_ is the *i*-th row of *A*.
2. Sort *v* from highest to lowest and let *β* = *v*_1_ = ∥*v*∥*_∞_*.
3. Iteratively compute the values 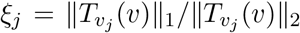 starting at *j* =1. If 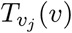 is the zero vector, move on to the next value of *j*. Stop computing when 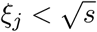.
4. Return the vector 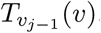. The support of this returned vector is the support of the solution to (4).

The algorithm relies on the fact that the solution 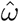 to (4) is given by a normalized soft-thresholding of a specific vector *v*; see Equations (1) and (6). For *s >* 1, note that the set 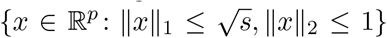 will look essentially like the set 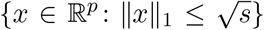 with the corners chopped off and rounded. Thus, we start by creating 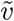, a non-zero soft-thresholding of *v* that has as few nonzero entries as possible (in the usual case, *v* has a unique largest entry, and thus 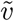 will have one nonzero entry so that it is pointing along one of the coordinate axes). We then soft-threshold *v* by smaller and smaller values so that 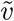 gains more non-zero coordinates and thus points further away from a coordinate axis. We stop when we find the point at which the 1-sphere 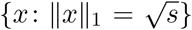 intersects the 2-sphere {*x* : ∥*x*∥ = 1}; the support of this vector are the features that we are interested in selecting.

There is also an even faster algorithm for solving a problem that is equivalent to (4) presented in [19]. This algorithm does not easily generalize to the multi-class problem in an interpretable way, however; see the upcoming work [14] for some discussions of these ideas.

#### Applying Select to rank transformed data

This section contains an algorithm RankBin for using Select along with the rank transformation. The inputs are a UMI counts matrix *X* ∈ ℝ^*n*×*p*^, a vector *τ* ∈ {±1}*^n^*, and a sparsity parameter *s*. Note that this is still a binary marker selection method, since the entries of *τ* are either +1 or −1. The extension to the multi-class case is handled in a one-vs-all manner and is explicitly described in the next section.

RankBin works as follows:

1. Construct the matrix *X*^*std*^ in the following manner: for all 1 ≤ *𝓁* ≤ *p*, let

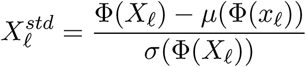
2. Let *τ*^*c*^ = Φ(*τ*) − *µ*(Φ(*τ*)).
3. Return Select(*X*^*std*^, *τ*^*c*^, *s*)

In the construction of *X*^*std*^, the columns of the data matrix *X* are standardized after they are rank transformed to more closely match the hypotheses of the theoretical results in [22]. In particular, the rows of the rank transformed and standardized data matrix *X*^*std*^ come from a bounded - and thus sub-Gaussian - distribution with mean 0 and variance 1 (the rows are not independent, however).

Moreover, motivated by the work in [13] and [18], the vector *τ* is replaced with Φ(*τ*) − *µ*(Φ(*τ*)) in RankBin. That is, the rank transformation is applied both to the data matrix *X* and the class indicator *τ*. In this case, the vector *v* that we soft threshold when we call Select (see 5 in Background) has entries given by

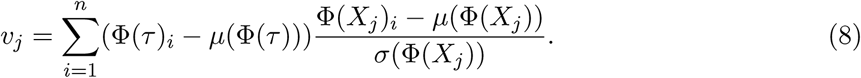

That is, entry *j* of *v* is (proportional to) the Spearman rank correlation between gene *j* and the vector *τ*. Thus, RankBin will select the genes that have the highest (absolute) Spearman correlation with the vector of class labels (compared to the method proposed in [13] that essentially selects the genes with the highest absolute Pearson corrlation with the vector *τ* - see (7)).

It is possible to show that replacing *τ* with 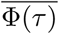 has no effect on the markers that are selected by the algorithm and thus the theoretical guarantees from [22] still apply. Algorithm RankBin is written with 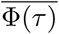 instead of *τ* to emphasize the connection with the Spearman rank correlation.

#### Rankcorr: Multi-class marker selection

RANKCORR works by fixing a parameter *s* and applying RankBin each of the cell types in the data set. Specifically, fix a sparsity parameter *s*; this parameter will be the same for all of the cell types. For cell type *j*, construct the vector*τ*^*j*^ with 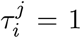 if cell *i* is in cell type *j* and 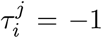 otherwise. Then run RankBin on the data matrix *X, τ_j_*, and the fixed sparsity parameter *s* to get the markers for cell type *j*. This will usually result in a different (informative) number of markers selected for each cell type.

When evaluating RANKCORR in this work, we take the union of all the markers selected for each cluster to get a set of markers that will represent all of the given cell types. This step is to allow for easier collection of benchmarking statistics - we would like to capture how well a selected set of markers informs us about an entire clustering. In practice, the sets of markers could be kept separate to give information about individual cell types. Note that there could still be duplicate markers in these sets - here, we do not address the problem of merging these sets in a smart way.

### Marker evaluation methods for experimental data

Below, we discuss two general procedures for the evaluation of a set of markers: supervised classification (that incorporates the given ground truth clustering as prior information) and unsupervised clustering (that does not). Both are discussed in more detail below.

Assuming that the data set contains *k* clusters, the result obtained by either classifying or clustering the data is a vector of predicted cell type labels 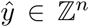. We would like to compare this to the “ground truth” cluster label vector *y* ∈ [*k*]*^n^*. The full information about the similarity between *ŷ* can be presented in terms of a confusion matrix; this is unwieldy when many such comparisons are required, however. For this reason, many summary statistics have been developed in the machine learning literature for the classification [23] and clustering [24] settings. We choose to examine several of these metrics in this work; the full list is summarized in Table 1.

#### Cross validation

In order to avoid overfitting, we perform all marker selection, classification, and clustering using 5-fold cross validation. Cross validation is commonly used in the computer science literature, as it allows for all of the data to be considered in test sets (we never test directly on the data that we trained with) - see Section 7.10 of [7]. See Figure 15 for a summary of this procedure.

**Figure 15:**
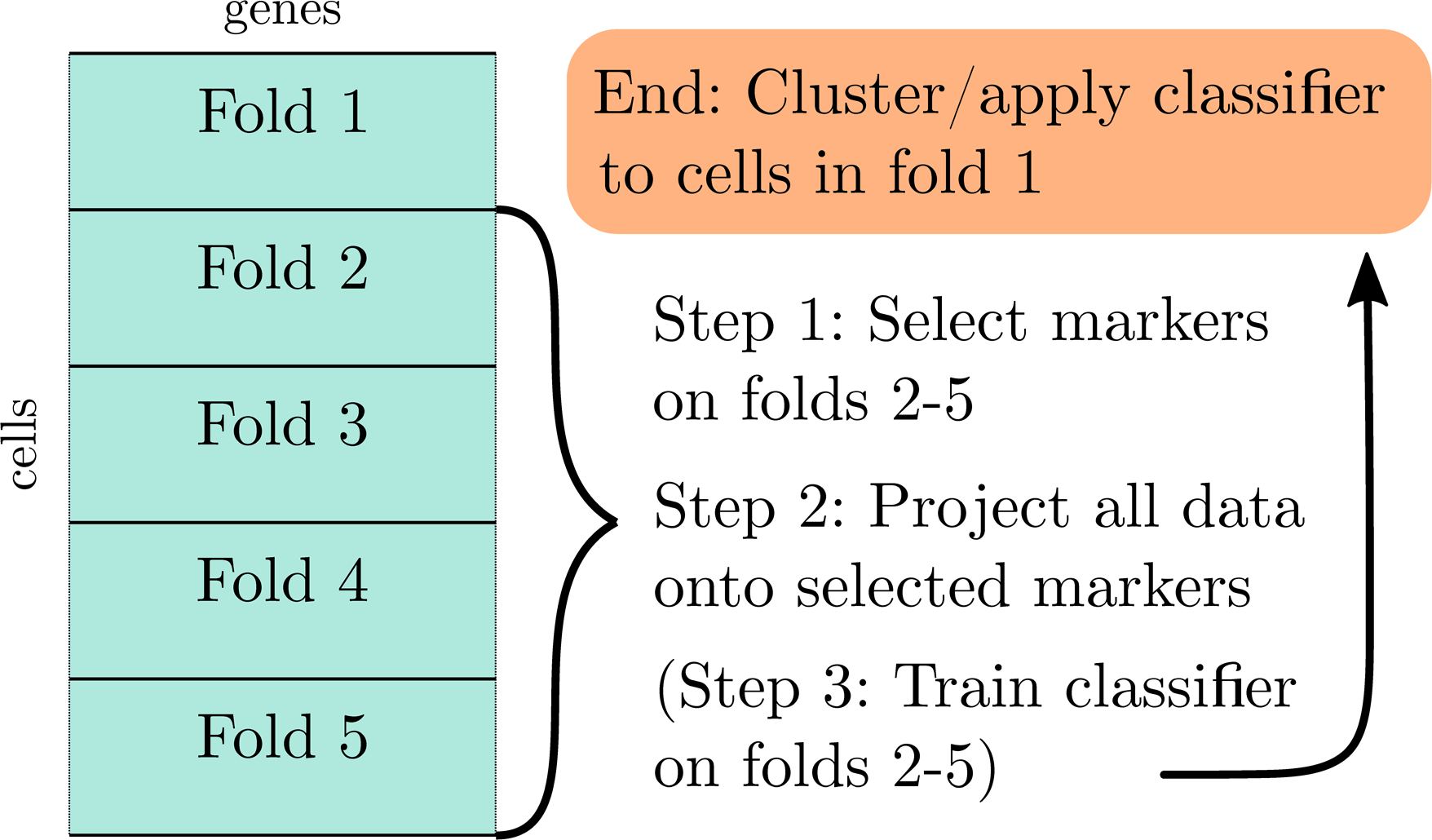
A visual description of 5 fold cross-validation

Specifically, we split the cells into five groups (called “folds”). For each fold, we combine the other four folds into one data set, find the markers on the dataset containing four folds, and train the classifier using the selected markers on the dataset containing four folds as the training data. We then apply the trained classifier to the initial fold and perform clustering on the initial fold using the markers that were selected on the other four folds. In this way, the initial fold is “test data” for the classifier/clustering metrics.

Repeating this process for all five folds creates a classification for the entire data set. On the other hand, we get a separate clustering for each fold, and these clustering solutions may be incompatible (they may contain different numbers of clusters, for example). See the section on clustering evaluation metrics for how we reconcile this.

#### Supervised classification

In order to incorporate information about the ground truth clustering into an evaluation metric, we train a multi-class classifier on the scRNA-Seq data using the cluster labels as the target output.

In order to evaluate the selected marker genes, we train the classifier using only the marker genes as the input data. When applied to a vector of counts (e.g. the counts of the markers in a cell), the classifier outputs a prediction of which cluster the vector belongs to.

#### Training a classifier

Given *y* ∈ ℕ*^n^*, a vector of cluster labels; *X* ∈ ℝ^*n*×*p*^, a scRNA-seq counts matrix; *S* = {*s*_1_, *…, s_R_*} ⊂ [*p*], a set of markers; we train a classifier Class in the following manner:

1. Normalize *X*.
2. Form a matrix Ξ ∈ ℝ^*n*×|*S*|^ from *X* by ignoring coordinates that aren’t in *S* (i.e. Ξ*_i_* = *X*_*s*_).
3. Train the classifier Class with Ξ as the input vectors and *y* as the target labels.
4. Return *h* : ℝ^|*S*|^ → ℕ, the classification function output by Class.train(Ξ, *y*).

In line 1 of this training procedure, we normalize the matrix *X*. It is possible to use any normalization for this step; for the purposes of our analysis we use a log normalization procedure that is commonly found in the scRNA-seq literature. Specifically, we perform a library-size normalization so that the sum of the entires in each row of *X* is 10, 000 and follow this by taking the base 2 logarithm of (1 plus) each entry of *X* to create a “log normalized” counts matrix.

Library size normalization was introduced in [28] to account for differences in capture efficiency between cells and taking a logarithm has it roots in bulk RNA-seq where it is used to attenuate technical variance (see [29]). Since log normalization of this type is often applied when clustering scRNA-seq counts data in a data processing pipeline, we apply log normalization when attempting to recover the information in the given clusters. It is important to note that the marker selection algorithms that we examine in this work do not assume that the input counts data are normalized (apart from when noted in their descriptions).

#### Classification evaluation metrics

We select markers and classify the cells using 5-fold cross-validation; see Figure 15. Once we have classified all cells in the data set, we examine how well the vector of classification labels matches the vector of ground truth cluster labels. Since we are in a classification framework, we use multi-class classification evaluation metrics for this purpose. In particular, we examine the classification error (1-accuracy) and precision of the classification compared to the known ground truth. For precision in a multi-class setting, we compute the precision for each class (as in a binary classification setting) and then take a weighted average of the per-class precision values, weighted by the class sizes. Finally, we also examine the Matthews correlation coefficient, which is a summary statistic that incorporates information about the entire confusion matrix. See [23] for more information about these statistics.

In all of the tests that we perform in this work, the precision and Matthews correlation coefficient curves look subjectively similar (though the actual values of the statistics do differ), while the classification error appears very similar to the other curves except it is flipped upside down. It is not clear why these summary statistics look as similar as they do.

#### Classifiers

We examine two classifiers to evaluate the marker sets (so that we are computing two classifications for each selected set of markers, and looking at all three metrics for both classifications).

The first is a simple (and fast) nearest centroids method that uses information about the original clustering to determine the locations of the cluster centroids. We refer to this as the Nearest Centroids Classifier (NCC). See the end of this section for a full description of the NCC. In the second, we use the Random Forest Classifier (RFC) that is implemented in the python package `scikit-learn` ([30]), version 0.20.0, with `nestimators` = 100. The summary statistics of the classifications produced when using the RFC are always better (more optimal) than the statistics that are produced when using the NCC. The overall shape of the curves produced using the RFC mostly mirror the curves produced using the NCC as well. The RFC is too slow to run on the largest data sets that we examine for testing. Since the RFC and NFC curves look similar for the smaller data sets, we are not concerned that we are missing information here.

Also note that, even with `nestimators` = 100, there is a significant amount of variability in the classification results obtained through the RFC. That is, running the RFC multiple times with the same set of markers will produce different classification results. See Figure 16 for a visualization of the differences in error rate that can be obtained when running the RFC twice on the same sets of markers (this example is created using the Paul data set; see the discussion of experimental data sets).

**Figure 16:**
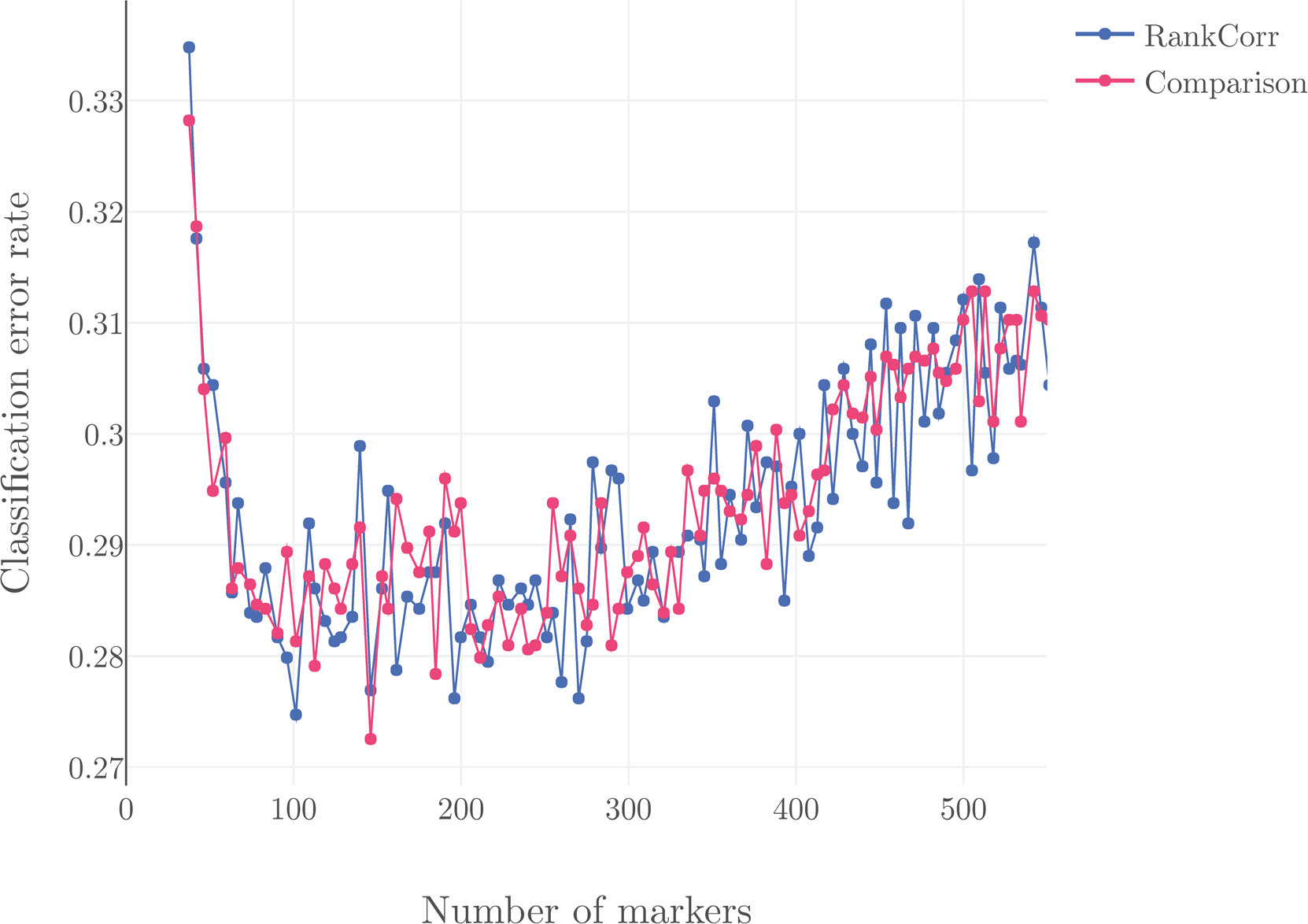
The classification accuracy under the RFC on the Paul data set (see Methods) run twice with the same markers used for each point. Significant variation is observed in the classification accuracy over the two classification attempts. Differences of nearly 2% are observed between the two curves.

#### The nearest centroids classifier (NCC)

The `NearestCentroids.train` method takes as input Ξ ∈ ℝ^*n*×*R*^, an scRNA-seq counts matrix; and *y* ∈ ℕ*^n^*, a vector of cluster labels. It runs in the following way:

1. Let *S*(*y*) = {*i* ∈ ℕ: *i* ∈ *y*} be the unique entires of *y*.
2. For each *k* ∈ *S*(*y*), let *C*_*k*_ = {*i* : *y*_*i*_ = *k*}.
3. For each *k* ∈ *S*(*y*), let 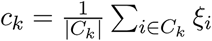, where *ξ*_*i*_ is the *i*-th row of Ξ.
4. Let *h* : [*n*] → ℕ be defined by *h*(*j*) = arg min*k* ∥*ξ*_*j*_ − *c*_*k*_∥_2_ (using Euclidean distance). Return *h*, the classification function.

#### Unsupervised clustering

Another natural way to measure the information in a selected set of markers is to cluster the data using only the selected coordinates in an unsupervised manner and compare this new clustering to the original clustering. Clustering scRNA-seq is itself a complicated problem that has inspired a great deal of study; here we restrict ourselves to Louvain clustering as implemented in the `scanpy` (version 1.3.7) package. Louvain clustering was introduced for use with scRNA-seq experiments in [31] and it is currently the recommended method for clustering scRNA-seq data in several commonly-used software packages including `scanpy` [32] and Seurat [33].

#### The clustering procedure

Louvain clustering on scRNA-seq data requires the input of *X* ∈ ℝ^*n*×*p*^, a scRNA-seq counts matrix; *S* = {*s*_1_, *…, s_R_*} ⊂ [*p*], a set of markers; *r*, a resolution parameter for Louvain clustering; and *k*, the number of nearest neighbors to consider in Louvain clustering.

It proceeds as follows:

1. Normalize *X*
2. Form a matrix Ξ ∈ ℝ^*n*×*|S|*^ from *X* by ignoring coordinates that aren’t in 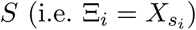.
3. For each cell *i*, find the *k* nearest neighbors to *i* according to Euclidean distance between the rows of Ξ.
4. Run Louvain clustering with resolution *r* on Ξ using the nearest neighbor data calculated in the previous step.
5. Let *h* : [*n*] → ℕ be a function that specifies the clustering. That is, *h*(*i*) = *j* means that cell *i* was placed into cluster *j*. Return *h*.

In line 1, we normalize the counts matrix *X*. As in the case of the supervised classification metrics, we apply log-normalization for this step. Also note that we do not perform any dimensionality reduction (e.g. PCA) before finding the nearest neighbors or performing the clustering. This is due to the fact that we projected the data onto the selected markers. These markers are meant to capture the important dimensions in the data - they are the features that have the most information about the clustering according to a marker selection algorithm. Thus, we work in the space spanned by these markers without performing any additional dimensionality reduction.

#### Clustering evaluation metrics

The unsupervised clustering is compared to the ground truth clustering using three metrics from the machine learning literature: the Adjusted Rand Index (ARI), Adjusted Mutual Information (AMI), and the Fowlkes-Mallows score (FMS). All three of these scores attempt to capture the amount of similarity between two groupings of one data set (e.g. the unsupervised clustering produced using a selected marker set and the ground truth clustering). They are also normalized scores: values near zero indicate that the cluster labels are close to random, while positive values indicate better performance. All of the scores have a maximum value of +1. Moreover, all three of these metrics do not make any assumption about the number of clusters: the unsupervised clustering can have a different number of clusters from the ground truth clustering and these indices can still be computed. See [24] for more information about these metrics.

We again use 5-fold cross-validation to compute the cluster performance markers discussed above. Note that the clustering solutions for the different folds may be incompatible: for example, the number of clusters in the Louvain cluster solution for the first fold may be different from the number of clusters in the Louvain cluster solution for the second fold, and there may be no obvious way to relate the clusters in the first fold to the clusters in the second fold. For this reason, we compute the clustering performance metrics separately on each fold, comparing the Louvain cluster solution to the ground truth clustering restricted to the fold. The scores that we report are averaged over all of the folds (and when we optimize over the resolution parameter *r*, discussed below, we find the optimal value of the average over the folds).

Note that some of the fine structure from the ground truth clustering may not be maintained in a specific fold and thus it is impossible to capture this structure when performing Louvain clustering on the fold. This means that the actual values of these metrics are not particularly informative - it is more useful to compare the different methods along a metric. In addition, in all of the Louvain clusterings for a specific data set, we fix the the value of *k*, the number of nearest neighbours that we consider. Thus, small differences between the scores are not particularly informative, as they could disappear if *k* was selected perfectly for each method. Nonetheless, it is useful to get an idea as to how well the markers selected by different algorithms could be used in an unsupervised manner to recover a given clustering.

#### Choice of Louvain clustering parameters

Louvain clustering requires the input of a number *k* of nearest neighbors and a resolution parameter *r*. It would be ideal to optimize both *k* and *r* for each set of markers on each data set for each clustering comparison metric; then we would be comparing the “optimal” performances of the marker selection algorithms under each metric. This is not computationally realistic for all of the data sets in consideration here.

Empirically, on the Paul and Zeisel data sets, we observed that all three of the clustering metrics are robust to changes in the value of *k* as long as *k* is chosen to be large enough (see Figures 17 and 18). On the other hand, changing the resolution parameter altered the metrics by large amounts, and different marker sets required different resolution parameters to obtain the optimal performance. Thus, for each dataset, we fix *k*. Then, for each of the three metrics and each set of markers, we optimize over *r* (we examine a grid from *r* = 0.1 to *r* = 3.0 with a step size of 0.1). This allows us to compute nearly optimal values for each metric and each set of selected markers. Importantly, the resolution parameter is not selected to optimize the performance of any one marker selection algorithm - the resolution is optimized for each marker selection algorithm separately.

**Figure 17:**
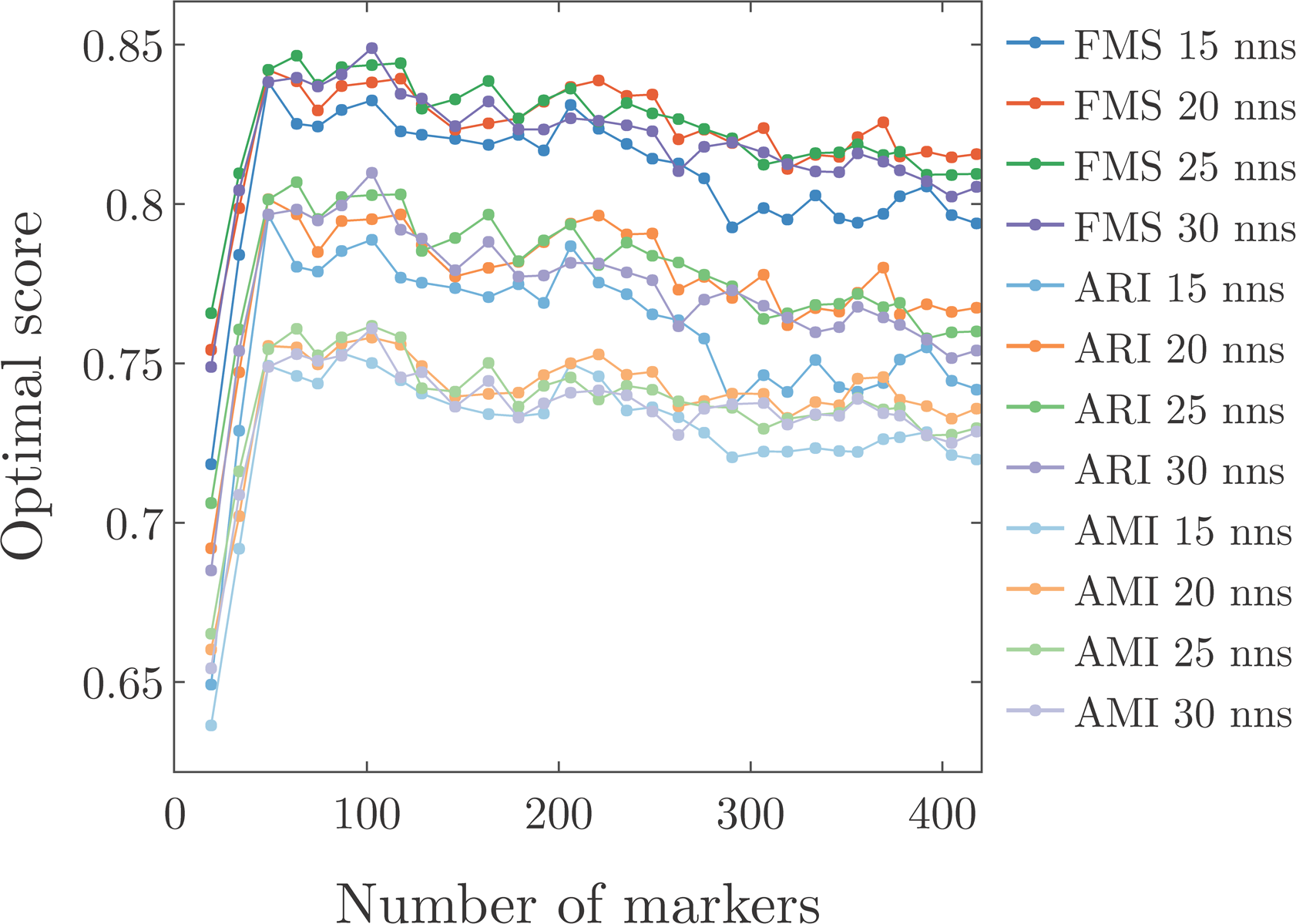
Effect of changing the number of nearest neighbors on the ARI, AMI, and FM scores for the Zeisel data set using RANKCORR to select markers. Clustering was performed with Louvain and the scores were optimized over the resolution. It appears that 15 nearest neighbors is too few, while 30 nearest neighbors is too many.

**Figure 18:**
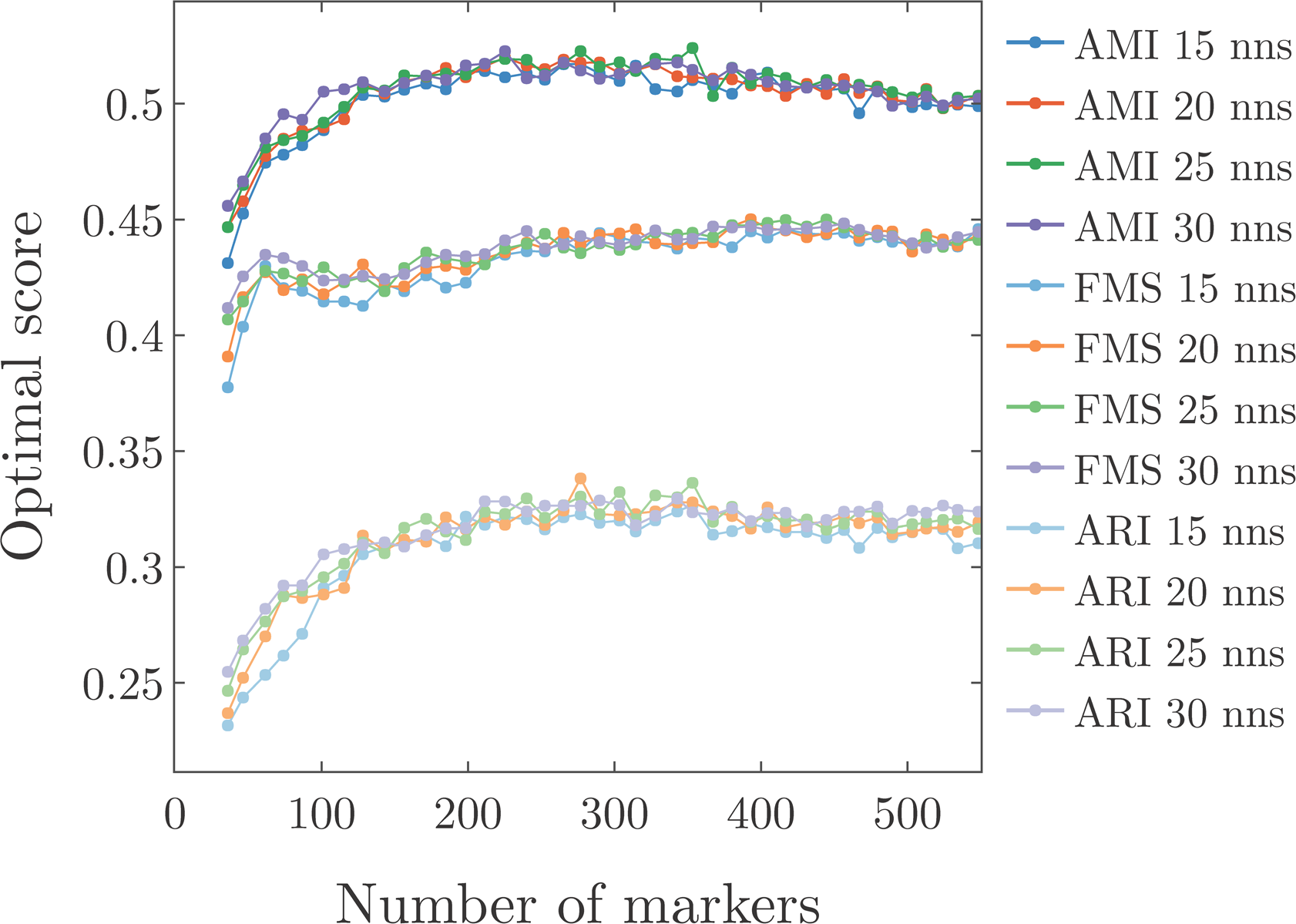
Effect of changing the number of nearest neighbors on the ARI, AMI, and FM scores for the Paul data set using RANKCORR to select markers. Clustering was performed with Louvain and the scores were optimized over the resolution. All of the choices of numbers of nearest neighbors produce similar curves for all three scores. Choosing 30 nearest neighbors appears to provide increased performance for small numbers of markers.

For a given dataset, the fixed value of *k* is obtained by a rough optimization strategy. On Zeisel and Paul, we optimize over both *k* and *r* (*r* varying from *r* = 0.1 to *r* = 3.0 with a step size of 0.1 and *k* varying from 15 to 30 with a step size of 5), using the RANKCORR algorithm and varying the number of markers selected to get a decent picture of the entire parameter space. The value of *k* is chosen to be the one that subjectively appears to optimize the performance of the majority of the metrics (see Figures 17 and 18).

On the Zeisel dataset, it appears that *k* = 15 nearest neighbors does not capture quite enough of the cluster structure, while *k* = 30 nearest neighbors results in lower scores than *k* = 25. We thus fix *k* at 25 for the unsupervised clustering evaluation on the Zeisel data set. See Figure 17 for the data that was used for this determination.

For the Paul data set, we observed that changing the number of nearest neighbors used in the Louvain clustering has little effect on the ARI, AMI, or FM scores. It appeared that the scores were slightly improved for *k* = 30 when small numbers of markers were selected, thus we fixed *k* at 30 for the Paul data set. See Figure 18.

The ZhengFull and ZhengSim data sets are large, and thus we focus on the ZhengSim data set when considering the unsupervised clustering metric. To estimate a value of *k*, the fixed number of nearest neighbours that we use for all of the clusterings, we computed a Louvain clustering that looks quite similar to the bulk labels in a UMAP plot. This clustering used 25 nearest neighbors (and used the top 50 PCs); thus we fix *k* at 25 for the Zheng data sets. See Figure 19 to see a comparison of the bulk labels and the generated Louvain clustering in UMAP space. Note that we still optimize over the resolution parameters separately for each method.

### Experimental data sets

We examine four publicly available experimental scRNA-seq data sets in this work. We focus on data sets that have been clustered, with clusters that have been biologically verified in some way. In addition, we mostly examine data sets that were collected using microfluidic protocols (Drop-seq, 10X) with UMIs. This is due to the fact that these protocols tend to collect a smaller number of reads in a larger number of cells (producing large amounts of sparse data). These data sets are summarized in Table 3. We discuss them further below. See the statement on data availabilty for how to obtain these data.

#### Zeisel

We work with one well-known reference fluidigm data sets. This is Zeisel, a data set consisting of mouse neuron cells that was introduced in [34]. Neuron cells are generally well-differentiated, and thus this data set contains distinct clusters that should be quite easy to separate. In [34], the authors have additionally used in-depth analysis with known markers to painstakingly label each cluster as a specific cell type. This labeling is the closest to an actual ground truth clustering of a dataset in the scRNA-seq literature - this fact makes Zeisel a valuable data set for our benchmarking purposes.

For our ground truth clustering, we consider only the nine major classes that the authors define in [34]. In addition, we pre-process the data set by selecting the top 5000 most variable genes, using the `cell_ranger` flavor of the `filter_genes_dispersion` function in the `scanpy` python package after library size normalization. We perform this pre-processing to speed up the marker selection process for the slower methods.

#### Paul

The smallest data set that we examine is Paul, a data set consisting of 2730 mouse bone marrow cells that was introduced in [35] and collected using the MARS-seq protocol. As opposed to Zeisel, bone marrow cells consist of progenitor cells that are in the process of differentiating. Thus, there are no well-defined cell types in the Paul data - the data appear in a continuous trajectory. The authors of Paul define discrete cell types along this trajectory based on known markers, however.

#### Zheng data sets

We perform an analysis of the data set introduced in [25] that consists of around 68 thousand human PBMCs from a single donor. These data were collected using 10x protocols; we refer to this full data set as ZhengFull. The cells were clustered (using k-means), and the clusters were assigned biological types based on known markers. The authors of [25] then took more cells (from the same donor) and isolated a set cells of each cell type that they found in their clustering of ZhengFull. They then sequenced the cells from the individual types. Finally, they used these pure samples to cluster the ZhengFull data set again: each cell is assigned to the type whose (bulk) profile correlates most strongly with the cell’s profile. We treat these bulk labels as the ground truth clustering for our experiments in this work. They can be found on the scanpy usage GitHub repository at https://github.com/theislab/scanpy_usage/blob/master/170503_ZHENG17/data/ZHENG17_bulk_lables.txt (we use commit `54607f0`).

We additionally generate a data set ZhengFilt from ZhengFull by restricting to the top 5000 most variable genes. We select the 5000 most variable genes by performing a library size normalization on the ZhengFull data set and then using the `cell_ranger` flavor of the `filter_genes_dispersion` function in the scanpy python package (see the data availabilty disclosure for more information about the scripts used to pre-process the data). We would like to see if restricting to highly variable genes hampers the marker selection process, or if the markers are mostly counted as highly variable genes.

#### 1.3 million mouse neurons

Finally, we examine 10xMouse, a data set consisting of 1.3 million mouse neurons generated using 10x protocols [26]. As noted above, neurons are well-differentiated into cell types, so this data set should contain well-separated clusters. The “ground truth” clustering that we consider for this data set is a graph-based (Louvain) clustering performed on the full 10xMouse dataset by the team behind `scanpy`. It can be found from the scanpy usage GitHub repository (https://github.com/theislab/scanpy_usage/tree/master/170522_visualizing_one_million_cells; we consider commit ba6eb85) As far as we know, this clustering has not been verified in any biological manner.

### Marker selection methods

A summary of the full set of marker selection methods that we consider in this work are found in Table 9 (a summary of the performance characteristics of the methods can be found in Table 2). We discuss the precise implementation details below.

**Table 9:** Differential expression methods tested in this paper. The top block consists of general statistical tests. The second block consists of methods that were designed specifically for scRNA-seq data. The third block consists of standard machine learning methods; Log. Reg. stands for Logistic Regression. The final block are the methods that are presented in this work; implementations of these methods can be found in the repository linked in the data availabilty disclosure.

| Method | Language | Package | Version | Ref |
| --- | --- | --- | --- | --- |
| t-test | Python 3.7 | **scanpy** | 1.3.7; see text |  |
| Wilcoxon | Python 3.7 | **scanpy** | 1.3.7; see text |  |
| edgeR | R (3.5.0) via Python | edgeR via rpy2 v2.9.4 | 3.24.1 | [5] |
| MAST | R (3.5.0) via Python | MAST via rpy2 v2.9.4 | 1.8.1 | [2] |
| scVI | Python 3.7 | Source from GitHub | 0.2.4 | [4] |
| SCDE | R (3.4.1) | SCDE | 2.6.0 | [3] |
| D<sup>3</sup>E | Python 2.7 and 3.5 | Source on GitHub | Commit efe21d1 | [1] |
| Elastic Nets | Python 3.7 | scikit-learn [30] | 0.20.0 | [11] |
| Log. Reg. | Python 3.7 | **scanpy** | 1.3.7; see text | [36] |
| RANKCORR | Python 3.7 | custom implementation |  |  |
| SPA | Python 3.7 | custom implementation |  | [13] |

#### Wilcoxon and the t-test

The t-test and Wilcoxon rank sum methods are general statistical methods that aren’t specifically designed for RNA-seq data, but they are still often used for the purposes of differential expression testing in the scRNA-seq literature. We use the Python `scanpy` package implementation to find Wilcoxson rank sum and t-test p-values with some editing to the file `_rank genes_groups.py` to fix several bugs (that are now fixed in the main release). See the data availability disclosure for how to find this file.

We use the version of the t-test in `scanpy` that overestimates the variance of the data. Both of these methods produce a score for each gene: when choosing the markers for the clusters, we use the absolute value of this score (so we would chose markers that have a large negative score as well). This is for more direct comparison to the RANKCORR method in which we choose markers by the absolute value of their coefficients. Finally, both of these methods correct the p-values that they produce using Benjamini-Hochberg correction.

#### edgeR and MAST

The methods edgeR, MAST, and SCDE were originally implemented in R. In order to run them with our existing framework, we use the `rpy2` (version 2.9.4) python package to access the methods through python.

Based on the results and scripts from [6], edgeR was run using the quasi-likelihood approach (QLF method) on the un-normalized scRNA-seq counts matrix *X*. For MAST, the data matrix *X* was normalized: the rows of *X* were scaled so that each row summed to 1 million (to approximate something that looks like “transcripts per millon”) to create a scaled matrix *X*^*s*^ and then each entry 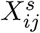 of *X*^*s*^ was replaced by log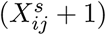.

Again following [6], we ran both edgeR and MAST in two ways. In the first way, we only consider the cluster label when fitting the statistical model; in the second way, we additionally include the fraction of genes that are detected in each cell (“detection rate”) as a covariate. In the rest of this paper, we refer to edgeR and MAST run the second way by edgeRdet and MASTdet respectively. According to the benchmarks considered in [6], edgeR (MAST) performs differently from edgeRdet (MASTdet).

#### scVI

scVI is implemented in python and utilizes GPUs for faster training of their model. Although the authors provide evidence that their code can handle a data set with one million cells (they test on the 10xMouse data set from [4]), scVI requires steep computational resources - around 75 GB of RAM to go with one core and one GPU. We have currently been unable to obtain this large amount of memory attached to a GPU, so we have been unable to reproduce their results here. One issue is that this method does not work with sparse data structures (or it makes them dense after loading them); thus, it has been computationally infeasible for us to run scVI on the larger data sets like 10xMouse

Another issue with scVI is that the differential expression methods included in the package are somewhat difficult to use (from a computational perspective). As far as we can tell, requesting information about differentially expressed genes from scVI produces a matrix of size larger than (10 *· n*) × *p*, where *n* is number of cells and *p* is the number of genes in the original data set (we were unable to precisely determine how the size of the matrix is computed). Even restricting to the top 3000 variable genes in the 10xMouse data set, this model matrix would require around 250 GB of memory to load into storage - in addition to the storage required for the 10xMouse dataset itself. Thus, although it may be possible to train the model on the 10xMouse data set, it will be nearly impossible with our computational resources to actually acquire the differential expression information from the trained model.

(An example of the extreme memory used by scVI: the Zeisel dataset takes approximately 5 MB to store in a dense format. The matrix produced during the differential expression computation method requires 4.1 GB. The actual computation of the Bayes factors - the generalization of a p-value produced by scVI - requires a peak of 15-16GB of memory during processing.)

#### SCDE

The SCDE package was too slow to be used on the real data sets: in testing, it was taking approximately one minute per cell to fit the model (on one core). Since we are performing 5-fold cross-validation, we would need to fit the model approximately 5 times. On one of the smaller data sets (Paul or Zeisel), this would require approximately 250 hours of computer time; it would be infeasible to train on the larger data sets. Since we are specifically developing methods for use with the large data sets that are appearing more often, we have excluded SCDE from our final analysis.

#### D^3^E

D^3^E is also implemented in python, but it has no support for sparse data structures; thus, running on the 10xMouse data set would require a very large amount of memory. Although the method allows for splitting the data into smaller segments (to allow for parallel computation), the full data set needs to be loaded into memory when initializing the process. In addition, when running on the Paul data set using the faster method-of-moments mode, D3E took about 25 minutes running on 10 cores (about 4 hours and 10 minutes total CPU time) to find markers for one cluster (vs the rest of the population). Since we need the p-values for all (*∼* 20) clusters for all 5 folds, this method would require approximately 40 hours on 10 cores. Although this is faster than SCDE, this would still be infeasible on the larger data sets, and thus we exclude D3E from our final analysis as well.

#### Elastic nets

The Spa method introduced in [17] is essentially an *L*_1_- and *L*_2_- regularized SVM without an offset (i.e. it finds a sparse separating hyperplane that assumed to pass though the origin, the instinct for this is given near Equation (3) in the background information). Thus, we also compare the performance of RANKCORR to that of the Elastic Nets version of LASSO: a least squares method with both *L*_1_ and *L*_2_ regularization. Elastic Nets has the two regularization parameters that need to be tweaked in order to find the optimal set of features; this requires extra cross-validation and therefore we are only able to run on the smaller Paul and Zeisel data sets. Although the `sklearn` package contains a method for finding the regularization parameters by cross-validation, it still takes a significant manual effort in order to find a range of the regularization parameters that capture the full possible behavior of the system but will also allow for the objective function to converge (in a reasonable number of iterations) the majority of the time. The timing information presented in Tables 5 and 4 only represents the run time of the method, and does not take into account this (time consuming) process of manipulating the data.

Another feature to note about the cross-validated elastic nets method is that it is (intentionally) a sparse method. Thus, scores are only generated for a small number of genes in each cluster - the genes that are specifically deemed “markers” for that cluster. It is not possible to compare the relative utilities of the genes that are not considered markers - each of those genes are given a score of 0. Thus, beyond a certain number of genes, it is not possible to get any more information from the markers selected by the elastic nets method. (You cannot, for example, request a “bad” marker in order to combine it with the information from other “good” markers).

#### Logistic regression

In a similar fashion to the method proposed here, logistic regression was proposed as a method for marker selection in [36]. In this, a regression is performed on each gene using the cluster label as the response variable. This is translated into a p-value via a likelihood ratio using the null model of logistic regression on the gene. This has been incorporated into the `scanpy` package, and thus we are able to run it on sparse data. We again have made some updates to the file `_rank genes_groups.py` in the `scanpy` package to fix some slight errors; see the data availability disclosure for where to find this edited file.

#### Spa

We also examine the performance of the method Spa introduced in [13] and analyzed further in [18]. As discussed in the background information, Spa was the original inspiration for this work, and also selects markers based on a sparsity parameter *s*. Spa also has two hyperparameters that we are required to optimize over, and this causes Spa to take considerably longer than RANKCORR to run for a fixed value of *s*. Moreover, since we are in a situation with no known ground truth, it is unclear what metric we would like to optimize when selecting these hyperparameters. For the current evaluation, we have minimized the classification error rate using the NCC (see information about the marker set evaluation metrics), but it is not clear that this would be the best metric to optimize in general. We choose the NCC classifier since the RFC exhibits a significant amount of variance - thus, optimizing the classification error rate according to the RFC classifier would produce an unstable set of markers (performing the optimization again would result in a different set of markers). We choose to optimize the supervised classification error (rather than one of the unsupervised clustering metrics) for the sake of speed - optimizing a slower evaluation metric would increase the time needed for the Spa marker selection method.

Another inconvenience of the Spa method is that the hyperparamters affect the number of markers that are selected for a fixed value of *s*. This makes the number of markers selected by Spa method more inconsistent and unpredictable. For example, it has occurred that the “optimal” (in terms of minimizing the classification error rate using the NCC, as discussed above) choice of hyperparameters for sparsity parameters *s*_1_ *> s*_2_ has resulted in a smaller number of markers selected for *s*_1_ than the “optimal” choice of hyperparameters for *s*_2_. That is, increasing *s* can lead to selecting smaller numbers of markers.

#### Rankcorr

The final method in the comparison is RANKCORR, the method introduced in this paper. It is important to note that the implementation of RANKCORR that we use here has not been fully optimized. Note that the major step (2) of the Select algorithm (see the details of RANKCORR) essentially consists of computing the dot product of each column of a data matrix with the cluster labels *τ*. The only other time consuming portion of Algorithm Select is computing the *𝓁*_2_ norm of a vector. These types of linear algebraic computation have fast implementations that are accessible from python (e.g. `numba`). We have not yet optimized the method to take advantage of all possible speed ups since RANKCORR runs quickly enough in our trials.

#### Random marker selection

Finally, as a sanity check, we choose markers uniformly at random (the same number of markers for each cluster).

### Generating marker sets of different sizes from algorithms other than RANKCORR

We wish to examine the relationship between the number of markers selected and the marker set performance metrics. In addition, for a fixed data set, we need to select markers for a given clustering - not just markers for a single cluster. Here we describe how we select a specific number of markers and how we merge lists of markers for individual clusters to make a marker list for the entire clustering.

For a differential expression method, we proceed in a one-vs-all fashion: letting *C* denote the number of clusters in the given clustering, we use the differential expression methods to find *C* vectors of p-values; the *i*-th vector corresponds to the comparison between cluster *i* and all of the other cells. For the sake of simplicity, we then include an equal number of markers for each cluster to create a set of markers for the clustering.

For example, we consider the classification error rate when the marker list consists of the three genes with the smallest p-values from each cluster (with duplicates removed). As mentioned in the introduction, this is a vast oversimplification of a tough problem - how to merge these lists of p-values in an optimal way, making sure that we have good representation of each cluster - but it allows for us to quickly and easily compare the methods that we present here. (Note that we would probably want to choose more markers for a cluster for which all p-values were large - we probably need more coordinates to distinguish this cluster from all of the others, even if those coordinates are not extremely informative. Thus, setting a p-value threshold could potentially perform worse than the method outlined here, as we may not select any markers for a certain cluster with a thresholding method.) The process of merging lists of p-values is left for future work.

For the elastic nets method, which selects one list of markers as optimal (without giving a score for all of the markers), we apply a similar strategy to approximate selecting a small number of markers. In particular, we choose an equal number of markers with the highest score for each cluster until we run out of markers to select. For example, when attempting to select 20 markers per cluster, we may include the top 20 markers for one cluster and all 18 of the markers that are selected for a different cluster.

### Generating synthetic data based on scRNA-seq data

In order to generate synthetic data that is made to look like an experimental droplet-based scRNA-seq data set, we use the `Splat` method from the R Splatter package (version 1.6.1) [27] in R version 3.5.0. The `Splat` method essentially works by selecting mean gene expression levels from a gamma distribution (with some “high expression outliers” included). Then, using these gene expression means, the actual counts for each cell are sampled from a modified Poisson distribution. Dropout can be added by randomly setting some of the counts to 0 at the end of the simulation process, though genes with higher average expression levels will experience less dropout. The required parameters for simulating counts in this way can be estimated from true experimental data using the Splatter package.

We use a samples from the data set consisting of purified (CD19+) B cells that is introduced in in order to estimate the `Splat` simulation parameters. In [25], the authors analyzed this dataset and saw only one cluster, suggesting that it consists mostly of one cell type. We have also combined it with the full ZhengFull dataset from [25] (see the descriptions of the experimental data sets) and observed good overlap with the cluster that the authors identified as B cells in ZhengFull when looking at a two dimensional UMAP visualization. This overlap appears in Figure 20. (The isolated cytotoxic T cell data set from [25] did not overlap with the T cells in the original data set ZhengSim as well, so we only considered the B cells.)

**Figure 19:**
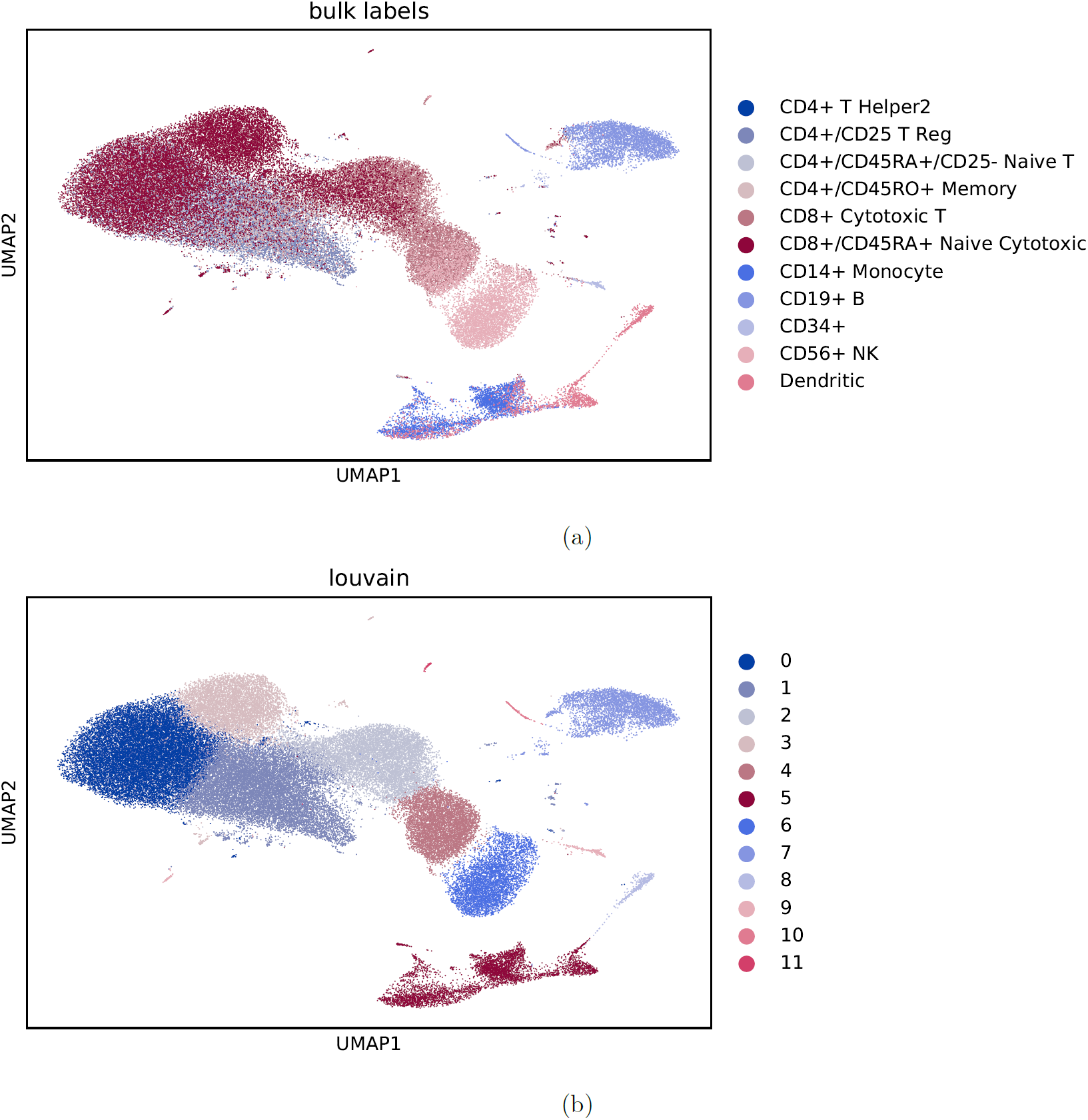
Clustering the 68k PBMC from [25] with Louvain clustering. (a) contains a UMAP plot of the bulk labels. (b) is a UMAP plot of a Louvain clustering of the data set. It was created by first filtering to the 1000 most variable genes (see the Methods). The Louvain algorithm was run on the top 50 PCs and used 25 nearest neighbours for each cell with a resolution parameter of 0.3. The Louvain clustering solution subjectively look similar to the bulk labels. The ARI for the clustering compared to the bulk labels is 0.345, the AMI is 0.565, and the FMS is 0.462 (these values have been rounded to 3 significant digits).

**Figure 20:**
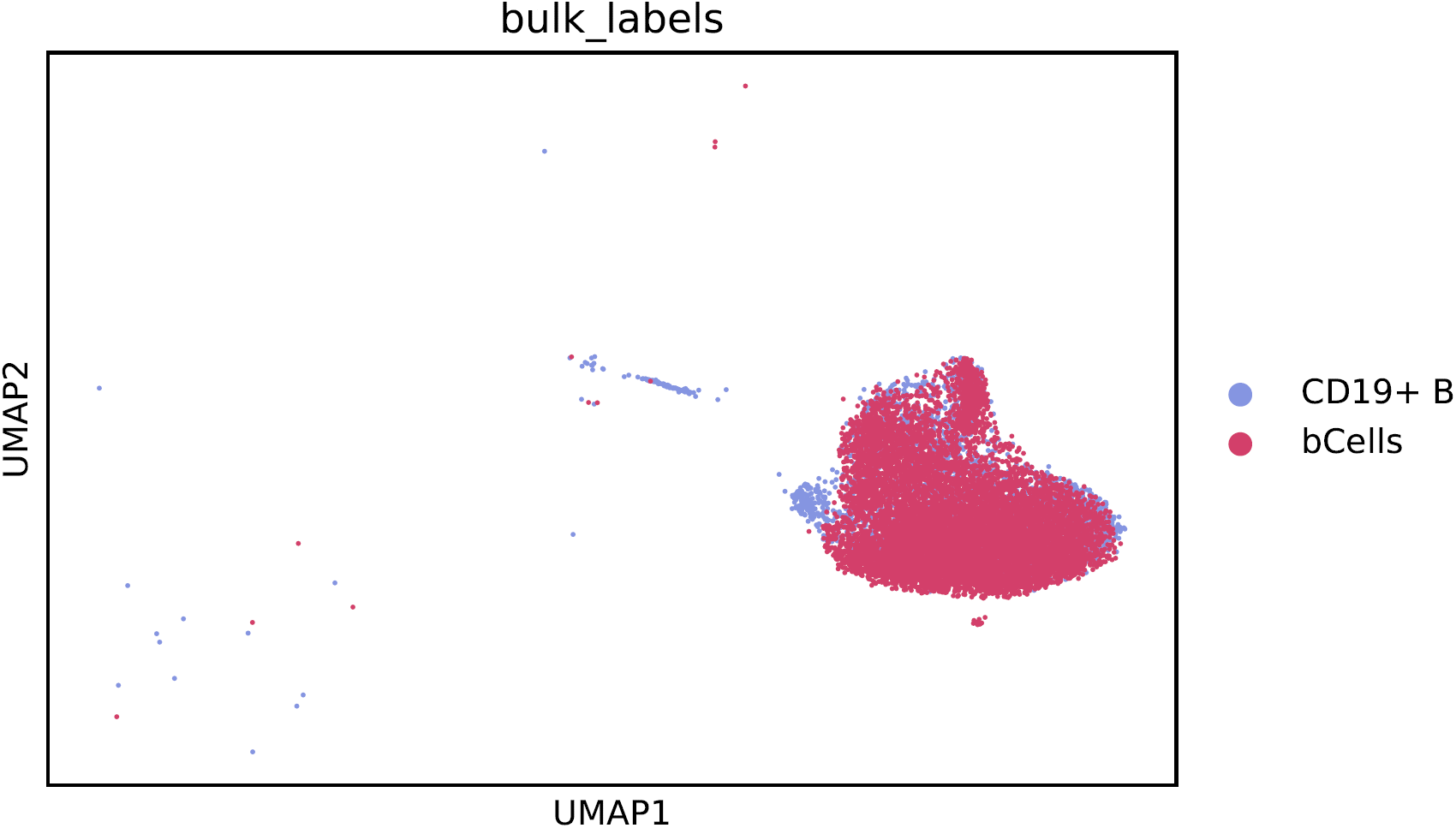
UMAP projection of the data consisting of ZhengFull combined with the isolated CD19+ B cell data set from [25] that was used to estimate parameters in Splatter simulations for generating synthetic data. We show only the isolated CD19+ sample (labeled “bCells”) and the cluster of B cells from ZhengFull. The overlap between the two clusters is quite good.

Some testing with Splatter showed that including dropout in the `Splat` simulation resulted in a simulated data set with a higher fraction of entries that are 0 than the original dataset. On the other hand, not including dropout resulted in similar fractions of entries that are 0 in the simulated and original datasets. Taking into account the fact that the Splat dropout randomly sets entries to 0 regardless of the size of those entries (a practice that we would argue is an unrealistic representation of actual dropout), we do not include additional dropout in our `Splat` simulations.

In the `Splat` method, differential expression is simulated by generating a multiplicative factor for each gene that is applied to the gene mean before the cell counts are created - a factor of 1 means that the gene is not differentially expressed. These multiplicative factors come from a lognormal distribution with location 0.1 and scale 0.4 - the default values in the Splatter package. We have not attempted to tweak these default parameters in this work. Using the default parameters, many of the “differentially expressed” genes have a differential expression multiplier that is between 0.9 and 1.1; for these genes, the gene mean is barely different between the two clusters. This creates a significant number of differentially expressed genes that are difficult to detect. See the simulated data results for further discussion.

For differential expression, we ask for Spatter to simulate two groups with 10% of the genes differentially expressed between the two groups: 10% of the genes in the first group are differentially expressed (i.e. have a differential expression multiplier not equal to 1), and none of the genes in the second group are differentially expressed. In this way, all differentially expressed genes can be considered to be marker genes for the first group - there are no overlaps between markers for the first and second groups. The direction of differential expression is randomly determined for each gene.

We use Splatter to generate 20 different simulated data sets from the CD19+ B cells dataset. See Figure 11 for a diagramme of the set-up. For all 20 simulated data sets, we simulate 5000 cells and the same number of genes that we input. The first 10 data sets are created by using the full (unfiltered) information from 10 random samples of 5000 cells from the B cell data set. This procedure results in sparse input data set of 5000 cells and about 20000 genes (the number of nonzero genes depends on the subsample). The output from this simulation is also very sparse. Since the differentially expressed genes are chosen at random, this means that many of the genes that are labeled as differentially expressed in the output data show low expression levels (often they are expressed in less than 10 cells).

To attempt avoid the issue of extreme low expression levels in the majority of the “differentially expressed” genes, we filter the genes of the simulated data via the method introduced in [37]: namely, place the genes in 20 bins based on their mean expression levels and select the genes with the highest dispersion from each bin. Using this method, we select the top 5000 most variable genes from the simulated data and we then use only genes these for marker selection. In the figures, we report these data under the heading “filtering after simulation.”

In order to explore this further, the second 10 simulated data sets are created by using only the top 5000 most variable genes in the original data as the input to Splatter. In this way, we are forcing the differentially expressed genes to look like genes that were originally highly variable.

## Declarations

### Availability of data and materials

The experimental data sets analysed during the current study are publicly available. They can be found in the following locations:

- Zeisel is found on the website of the authors of [34]: http://linnarssonlab.org/cortex/. The data are also available on the GEO (GSE60361).
- Paul is found in the `scanpy` python package - we consider the version obtained by calling the `scanpy.api.datasets.paul15`() function. The clustering is included in the resulting `Anndata` object under the heading `paul15 clusters`. The data are also available on the GEO (GSE72857).
- ZhengFull and ZhengFilt are (subsets) of the data sets introduced in [25]. The full data set can be found on the 10x website (https://support.10xgenomics.com/single-cell-gene-expression/datasets/1.1.0/fresh_68k_pbmc_donor_a) as well as on the SRA (SRP073767). The biologically motivated bulk labels can be found on the `scanpy usage` GitHub repository at https://github.com/theislab/scanpy_usage/blob/master/170503_ZHENG17/data/ZHENG17_bulk_lables.txt (we use commit `54607f0`).
- 10xMouse is available for download on the 10x website (https://support.10xgenomics.com/single-cell-gene-expression/datasets/1.3.0/1M_neurons. The clustering analysed in this manuscript can be found on the `scanpy_usage` GitHub repository (https://github.com/theislab/scanpy_usage/tree/master/170522_visualizing_one_million_cells; we consider commit `ba6eb85`)

The synthetic data analysed in this manuscript is based on the CD19+ B cell data set from [25]. It can be found on the 10x website at https://support.10xgenomics.com/single-cell-gene-expression/datasets/1.1.0/b_cells. The synthetic data sets themselves are available from the author on request.

All scripts that were used for marker selection and data processing can be found at the GitHub repository located at https://github.com/ahsv/marker-selection-code. This includes the implementations of Spa and RANKCORR.

## Acknowledgments

The authors wish to thANK Umang Varma for contributing his valuable programming experience and his mathematical insights. ThANKs also to the members of the Michigan Center for Single-Cell Genomic Data Analytics for their advice and feedback. Special thANKs to Jun Li and Justin Colacino for their biological wisdom and ideas, as well as to Xiang Zhou and his students Lulu Shang and Shiquan Sun for their comments on a previous draft of this work. This research was supported in part through computational resources and services provided by Advanced Research Computing at the University of Michigan, Ann Arbor.

## Additional Files

### Additional file 1 — Supplementary Figures

Additional figures containing data that is similar to the data already shown in the manuscript.

