## Additional File 1 for "Comparison of marker selection methods for high throughput scRNA-seq data"

Anna C. Gilbert Alexander Vargo

June 14, 2019

This file contains pictures to supplement the content in the manuscript

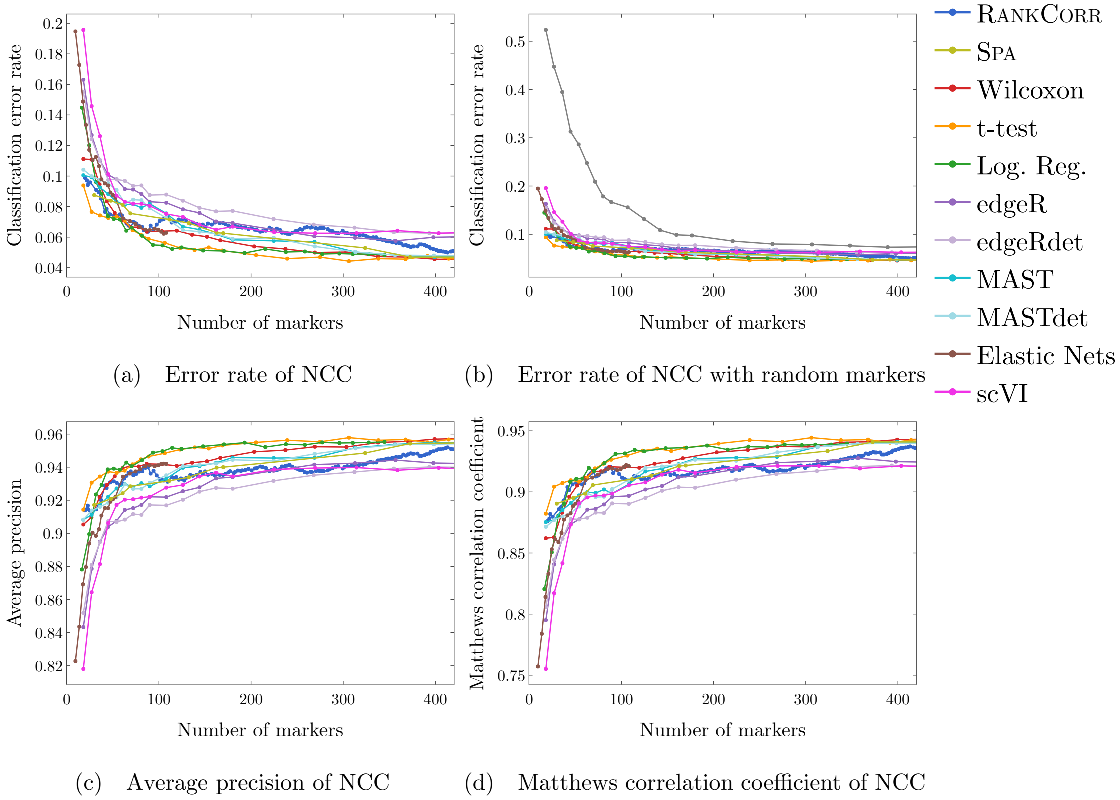


Figure 1: Supervised classification metrics for the Zeisel data set using the nearest centroid classifier (NCC). (a) is the classification error rate figure found in Figure 2(a) of the main manuscript, for reference here. (b) is the same classification error rate including a curve corresponding to random marker selection (the curve that starts at around 50% error, in grey). (c) contains the precision and (d) contains the Matthews correlation coefficient. Data from random marker selection is not included in figures (c) and (d).

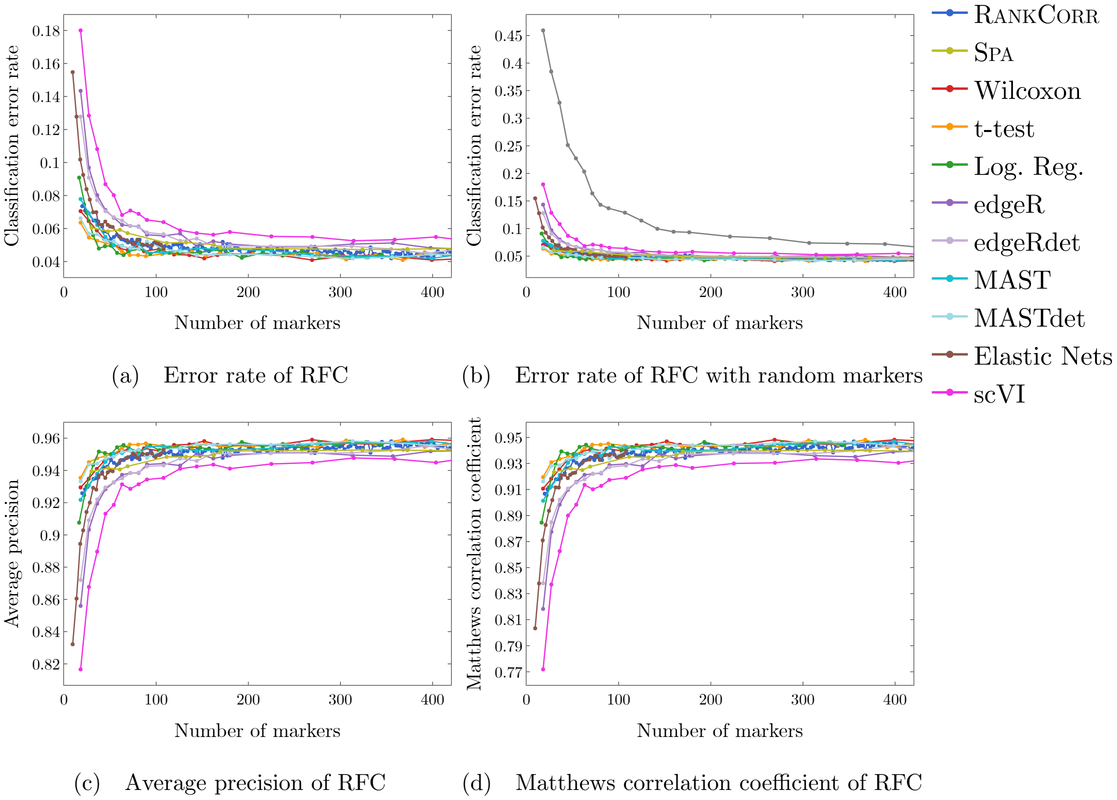


Figure 2: Supervised classification metrics for the Zeisel data set using the random forests classifier (RFC). (a) is the classification error rate figure found in Figure 2(c) of the main manuscript, for reference here. (b) is the same classification error rate including a curve corresponding to random marker selection (the curve that starts at around 45% error, in grey). (c) contains the average precision and (d) contains the Matthews correlation coefficient. Data from random marker selection is not included in figures (c) and (d).

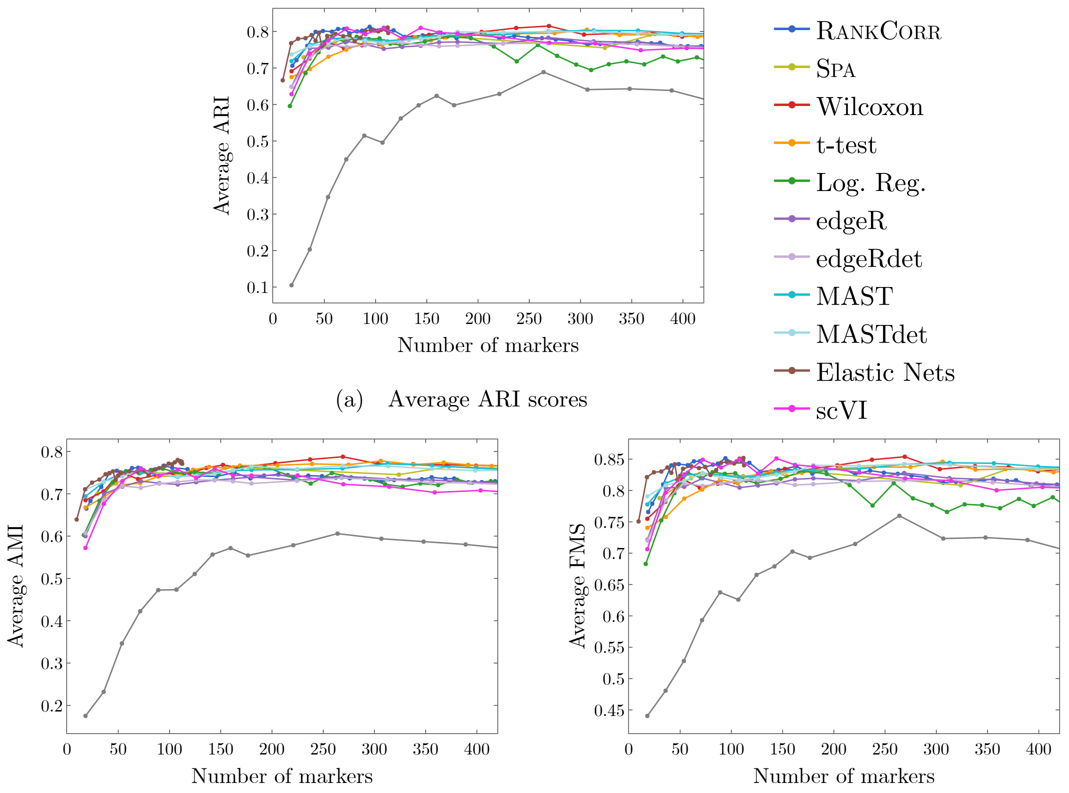


(a) Average ARI scores

(b) Average AMI scores

(c) Average Fowlkes-Mallows scores

Figure 3: Unsupervised clustering metrics for the Zeisel data set including data from random marker selection. The curve corresponding to random marker selection is shown in grey and is the lowest (worst) curve in all three plots. The ARI score is shown in (a), the AMI score is shown in (b), and the Fowlkes-Mallows score is shown in (c). The clustering is carried out using 5-fold cross validation and scores are averaged across folds.

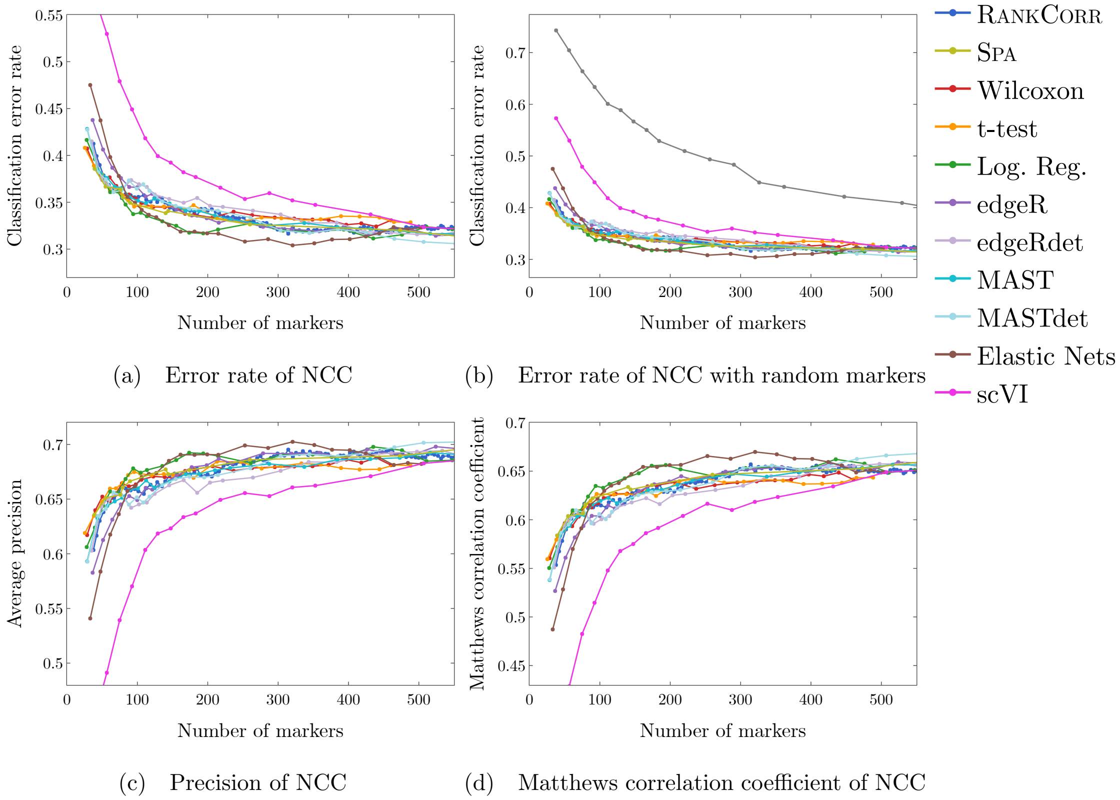


Figure 4: Supervised classification metrics for the Paul data set using the nearest centroid classifier (NCC). (a) is the classification error rate figure found in Figure 4(a) of the main manuscript, for reference here. (b) is the same classification error rate including a curve corresponding to random marker selection (the curve that starts at around 75% error, in grey). (c) contains the average precision and (d) contains the Matthews correlation coefficient. Data from random marker selection is not included in figures (c) and (d).

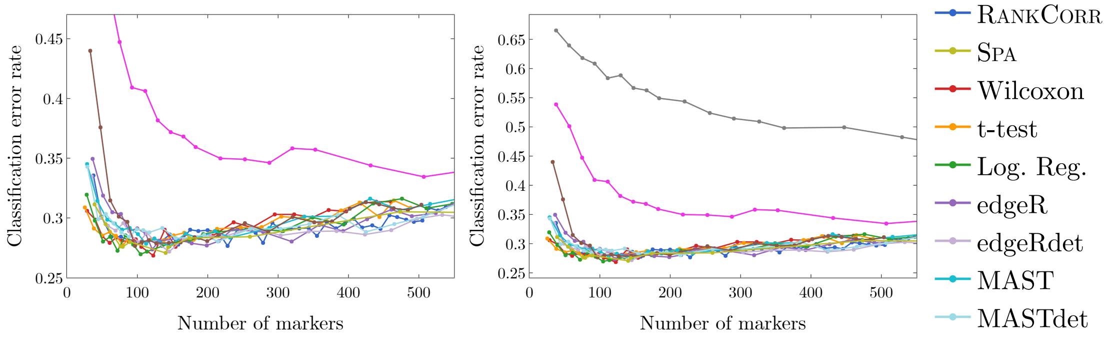


(a) Error rate of RFC

(b) Error rate of RFC with random markers

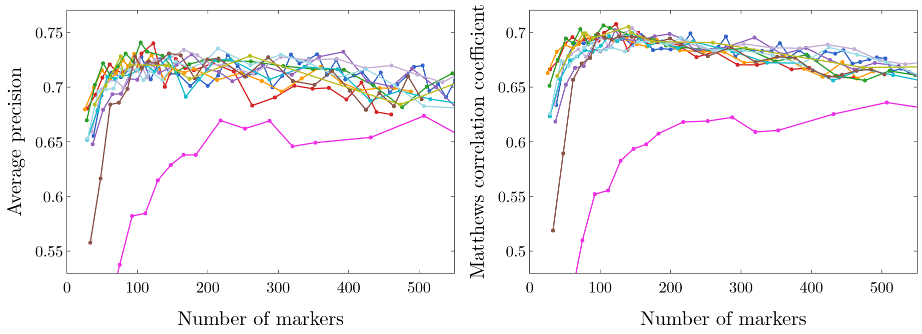


(c) Precision of RFC

(d) Matthews correlation coefficient of RFC

Figure 5: Supervised classification metrics for the Paul data set using the random forests classifier (RFC). (a) is the classification error rate figure found in Figure 4(c) of the main manuscript, for reference here. (b) is the same classification error rate including a curve corresponding to random marker selection (the curve that starts at around 65% error, in grey). (c) contains the average precision and (d) contains the Matthews correlation coefficient. Data from random marker selection is not included in figures (c) and (d).

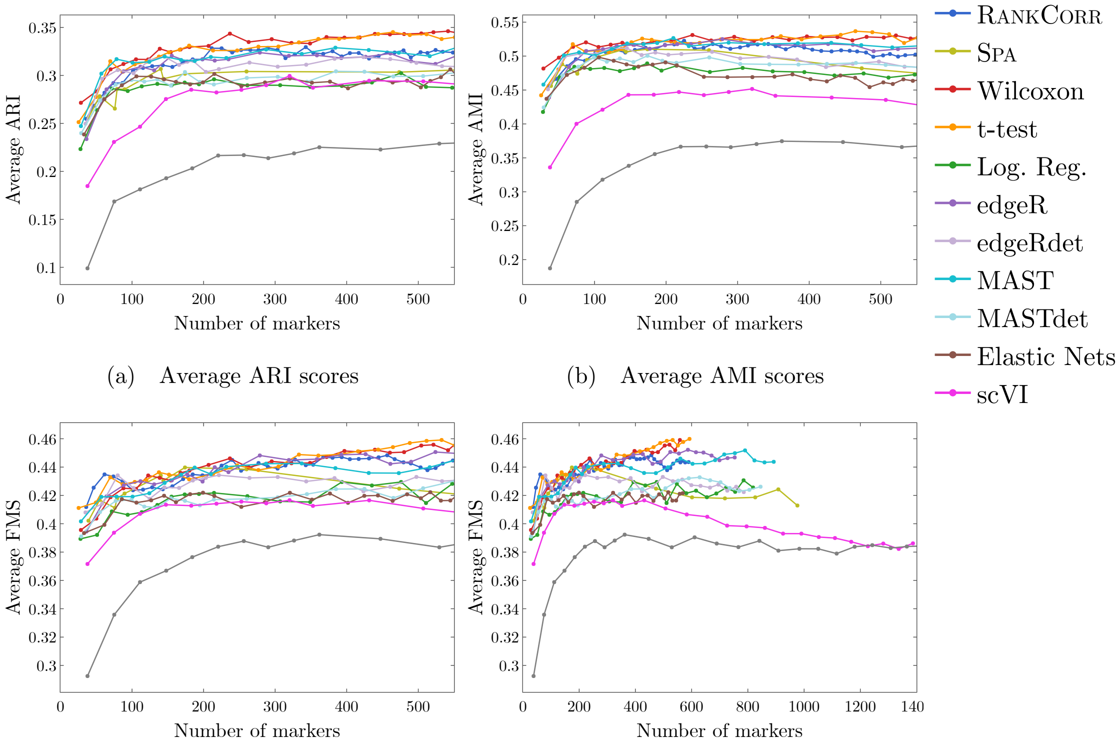


(a) Average ARI scores

(b) Average AMI scores

(c) Average Fowlkes-Mallows scores

(d) Tail behavior of Fowlkes-Mallows scores

Figure 6: Unsupervised clustering metrics for the Paul data set including data from random marker selection. The curve corresponding to random marker selection is shown in grey and is the lowest (worst) curve in all four plots. The ARI score is shown in (a), the AMI score is shown in (b), and the Fowlkes-Mallows score is shown in (c). (d) contains the FM scores for larger numbers of markers selected, showing the scVI does approach the behavior of random marker selection in this case. Each clustering is carried out using 5-fold cross validation and scores are averaged across folds.

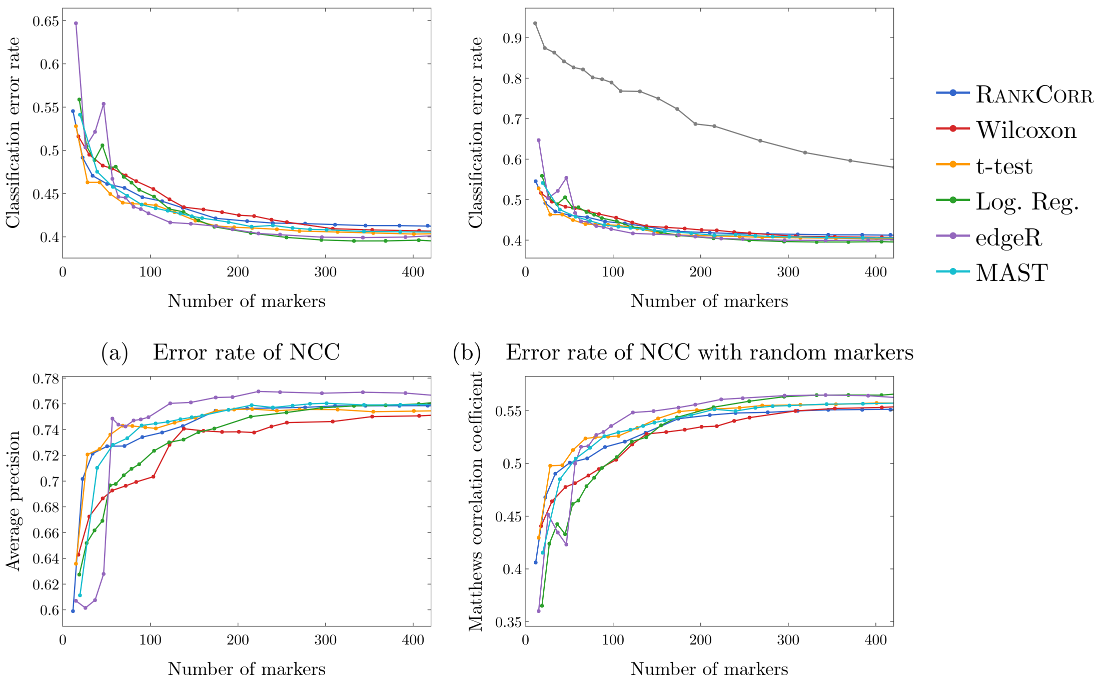


(a) Error rate of NCC (b) Error rate of NCC with random markers

(c) Average precision of NCC (d) Matthews correlation coefficient of NCC

Figure 7: Supervised classification metrics for the ZhengFilt data set using the nearest centroid classifier (NCC). (a) is the classification error rate figure found in Figure 6(a) of the main manuscript, for reference here. (b) is the same classification error rate including a curve corresponding to random marker selection (the curve that starts at around 90% error, in grey). (c) contains the average precision found in Figure 6(b) of the main manuscript, again for reference. (d) contains the Matthews correlation coefficient. Data from random marker selection is not included in (d). Note that (d) looks subjectively similar to the classification accuracy (1− classification error rate from (a)).

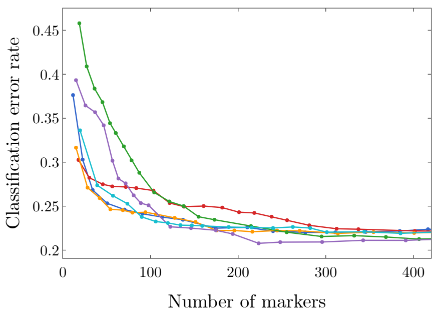


(a) Error rate of RFC

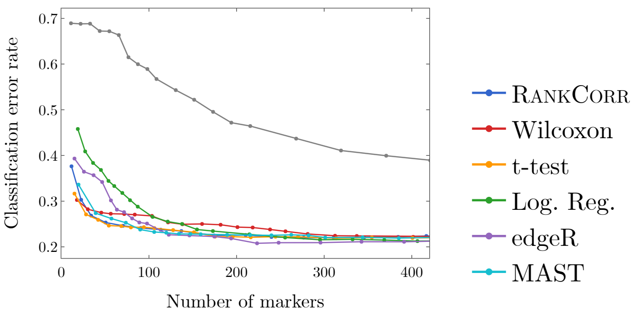


(b) Error rate of RFC with random markers

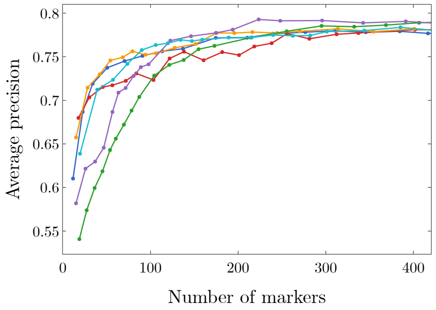


(c) Average precision of RFC

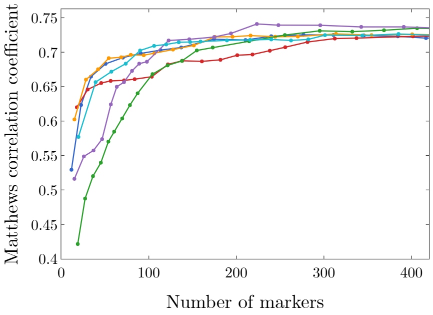


(d) Matthews correlation coefficient of RFC

Figure 8: Supervised classification metrics for the ZhengFilt data set using the random forests classifier (RFC). (a) is the classification error rate figure found in Figure 7(a) of the main manuscript, for reference here. (b) is the same classification error rate including a curve corresponding to random marker selection (the curve that starts at around 70% error, in grey). (c) contains the average precision data found in Figure 7(b) of the main manuscript, again for reference. (d) contains the Matthews correlation coefficient. Data from random marker selection is not included in (d). Note that (d) looks subjectively similar to the classification accuracy (1− classification error rate from (a)).

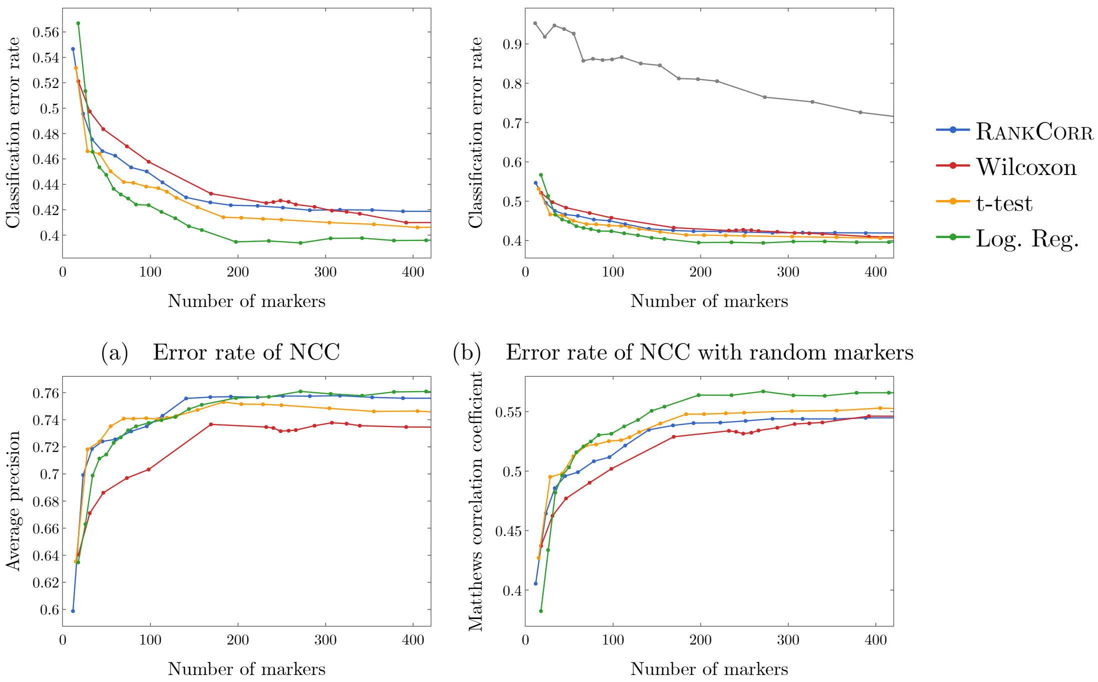


(a) Error rate of NCC

(b) Error rate of NCC with random markers

(c) Average precision of NCC

(d) Matthews correlation coefficient of NCC

Figure 9: Supervised classification metrics for the ZhengFull data set using the nearest centroid classifier (NCC). (a) is the classification error rate figure found in Figure 6(c) of the main manuscript, for reference here. (b) is the same classification error rate including a curve corresponding to random marker selection (the curve that starts at around 90% error, in grey). (c) contains the average precision found in Figure 6(d) of the main manuscript, again for reference. (d) contains the Matthews correlation coefficient. Data from random marker selection is not included in (d). Note that (d) looks subjectively similar to the classification accuracy (1− classification error rate from (a)).

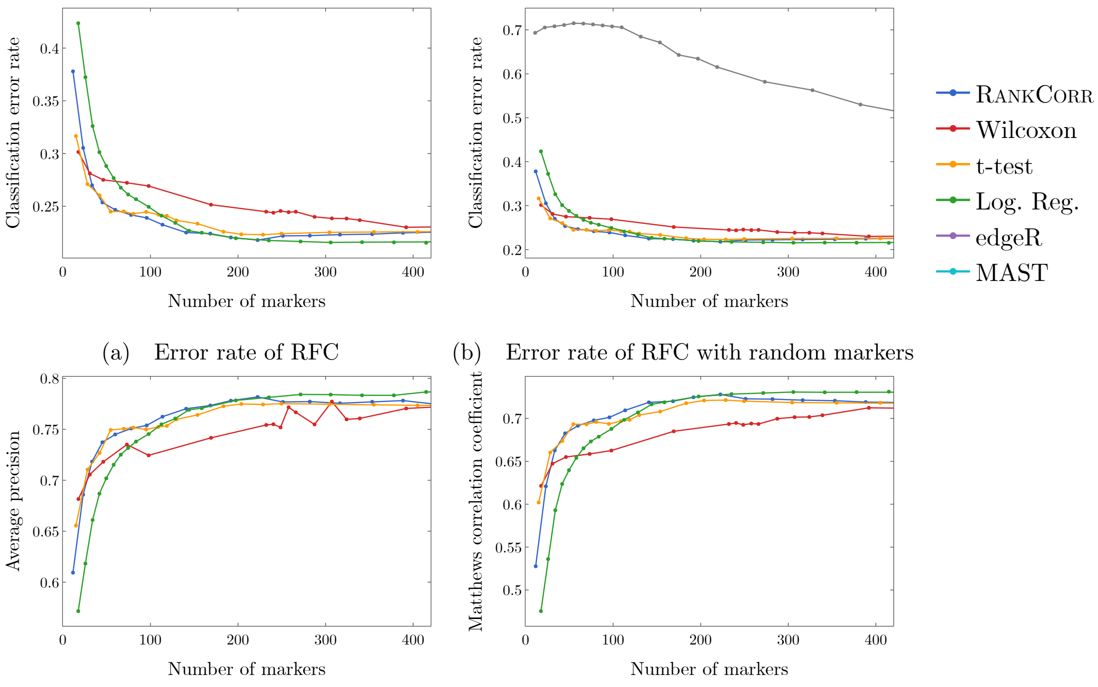


(a) Error rate of RFC

(b) Error rate of RFC with random markers

(c) Average precision of RFC

(d) Matthews correlation coefficient of RFC

Figure 10: Supervised classification metrics for the ZhengFull data set using the random forests classifier (RFC). (a) is the classification error rate figure found in Figure 7(c) of the main manuscript, for reference here. (b) is the same classification error rate including a curve corresponding to random marker selection (the curve that starts at around 70% error, in grey). (c) contains the average precision data found in Figure 7(d) of the main manuscript, again for reference. (d) contains the Matthews correlation coefficient. Data from random marker selection is not included in (d). Note that (d) looks subjectively similar to the classification accuracy (1− classification error rate from (a)).

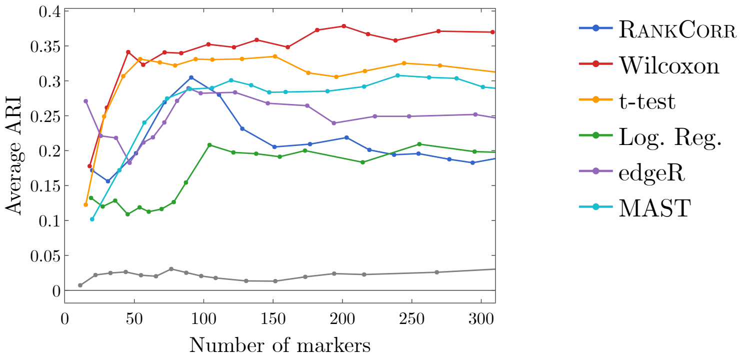


(a) Average ARI scores

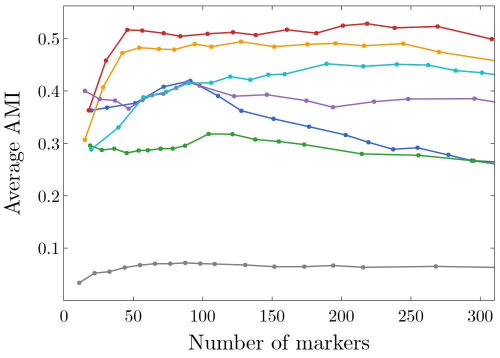


(b) Average AMI scores

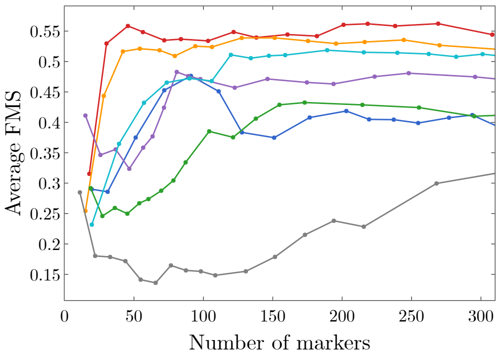


(c) Average Fowlkes-Mallows scores

Figure 11: Unsupervised clustering metrics for the ZhengFilt data set including data from random marker selection. The curve corresponding to random marker selection is shown in grey and is the lowest (worst) curve in all three plots. The ARI score is shown in (a), the AMI score is shown in (b), and the Fowlkes-Mallows score is shown in (c). Each clustering is carried out using 5-fold cross validation and scores are averaged across folds.

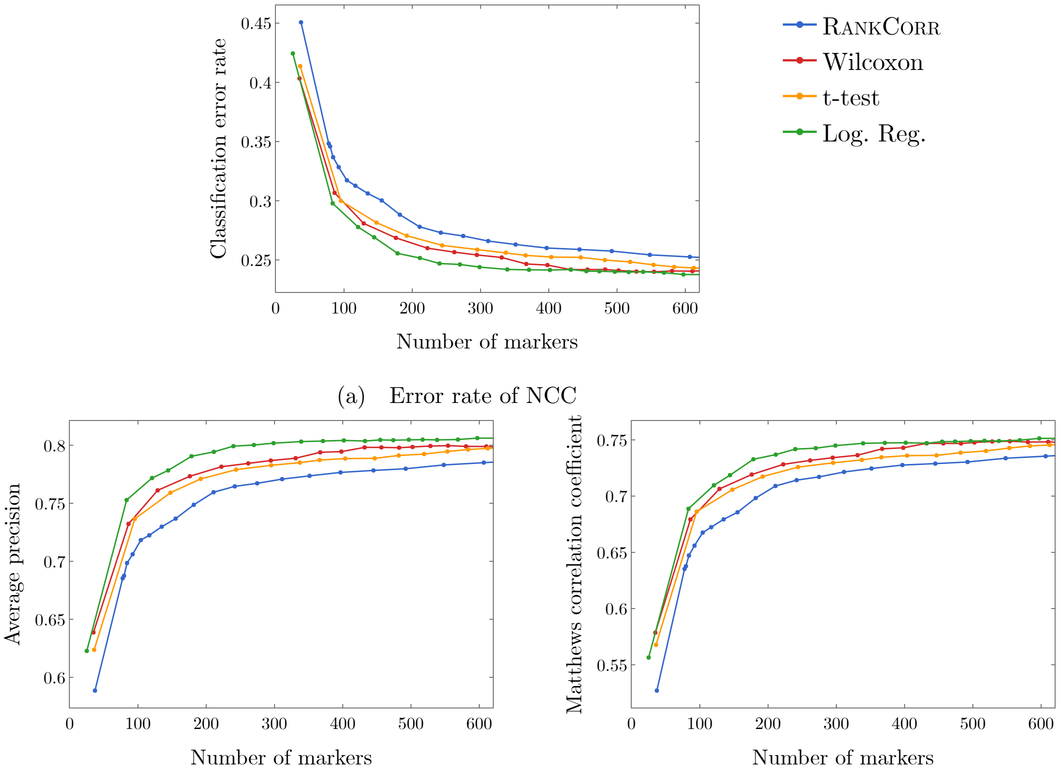


(a) Error rate of NCC

(b) Average precision of RFC

(c) Matthews correlation coefficient of NCC

Figure 12: Supervised classification metrics for the 10xMouse data set using the nearest centroid classifier (NCC). (a) is the classification error rate figure found in Figure 9 of the main manuscript, for reference here. (b) contains the average precision curves and (c) contains the Matthews correlation coefficient curves. We do not compare to random markers on this data set.
